# Proteomic and single-cell transcriptomic dissection of human plasmacytoid dendritic cell response to influenza virus

**DOI:** 10.1101/2021.11.01.466839

**Authors:** Mustafa H. Ghanem, Andrew J. Shih, Houman Khalili, Emily Werth, Jayanta K. Chakrabarty, Lewis M. Brown, Kim R. Simpfendorfer, Peter K. Gregersen

## Abstract

Plasmacytoid dendritic cells [pDCs] represent a rare innate immune subset uniquely endowed with the capacity to produce substantial amounts of type-I interferons [IFN-I]. This function of pDCs is critical for effective antiviral defenses and has been implicated in autoimmunity. While IFN-I and select cytokines have been recognized as pDC secreted products, a comprehensive agnostic profiling of the pDC secretome in response to a physiologic stimulus has not been reported. We applied LC-MS/MS to catalogue the repertoire of proteins secreted by pDCs in response to challenge with live influenza H1N1. Additionally, using single-cell RNA-seq [scRNA-seq], we perform multidimensional analyses of pDC transcriptional diversification following stimulation. Our data reveal an abundance of protein species released by pDCs in addition to IFN-I, and evidence highly specialized roles within the pDC population ranging from dedicated cytokine super-producers to cells with APC-like functions. Moreover, dynamic expression of transcription factors and surface markers characterize activated pDC fates.

## Introduction

pDCs were presumptively identified in 1958 as cells with a ‘plasma cell-like’ morphology observed in human splenic white pulp and lymph nodes (Lennert and Remmele, 1958). Decades later, these cells were isolated from the circulation as the ‘natural IFN-producing cells’ (Siegal et al., 1999). pDCs are notable for their constitutive high expression of the IFN-inducer IRF7 and the nucleic acid sensors TLR7 and TLR9. Together, with their highly developed endoplasmic reticulum [ER], pDCs are poised to rapidly respond to viral nucleic acid by producing massive amounts of IFN-I. This response is aided by the lack of introns in IFN-I genes, circumventing the steps required for pre-mRNA splicing. Secretion of IFN-I is critical to antiviral defense: signaling downstream of the ubiquitous IFN-α/β receptor results in the induction of antiviral programs, collectively known as the ‘interferon signature.’ The aberrant activation of the IFN pathway is a hallmark of several autoimmune diseases, most notably systemic lupus erythematosus (Baechler et al., 2003; Crow and Ronnblom, 2019).

Since their initial description, pDCs have mostly been studied in the context of IFN-I production. IFN-I comprises of IFN-α (itself a family of 13 genes encoding 12 distinct polypeptides), IFN-β, IFN-κ, IFN-ω and IFN-ε. Traditional approaches have also shown that activated pDCs secrete varying amounts of type-III interferon [IFN-III], TNF-α, IL-6 and CXCL10. Historically, these methods have relied on pDC stimulation with well-characterized synthetic TLR agonists and used analyte-specific ELISAs for readout. To our knowledge, a comprehensive profiling of the primary human pDC secretome in response to a physiologically relevant stimulus has not been reported. Moreover, whether pDCs maintain production of secreted proteins in the baseline ‘idle’ state has not been explored. Plasma cells, whose morphology bears striking similarity to pDCs, are likewise a rare immune subset, but are notable for their baseline secretion of massive amounts of immunoglobulin necessary for humoral immunity. To address these gaps in knowledge, we catalogued the pDC protein secretome from primary human pDCs using LC-MS/MS for both unstimulated and influenza H1N1 treated conditions. Our data reveal an abundance of known pDC products in addition to novel pDC-sourced proteins.

Another characteristic feature of pDCs is their ‘diversity’ following stimulation. It has long been known that upon stimulation, only a minority of pDCs ultimately produce IFN-I, with the remainder assuming alternative or unrecognized roles. These studies have been limited by the monoclonal antibodies used in flow cytometry, which cannot capture the breadth of pDC heterogeneity. We therefore endeavored to characterize pDC transcriptional states immediately *ex vivo* as well as following stimulation with influenza H1N1 using scRNA-seq. Our data suggest the presence of preexisting pDC transcriptional variation, and further define clearly specialized cell fates post-stimulation.

## Results

### pDC secretome profiling by LC-MS/MS

A major challenge to proteomic profiling of serum or culture media is the phenomenon of ionization suppression, whereby species dominant in a solution (such as albumin and immunoglobulin in serum) compete for ionization energy with less abundant species thereby impeding their detection (Polson et al., 2003). To minimize the necessary media protein supplement, we tested a variety of culture conditions assaying for pDC viability and IFN-I output as a measure of media compatibility. pDCs remained viable and produced significant quantities of IFN-α with fetal calf serum (FCS) levels down to 1%. However, we observed that even trace amounts of FCS accounted for the majority of media protein following pDC culture. Therefore, we opted to use serum-free media supplemented with minimal amounts of albumin, insulin and holo-transferrin. pDCs maintained IFN-I production capacity and viability in this media similar to media containing FCS **(Figure S1)**. Media completely free of exogenous protein was incompatible with pDC interferon production (data not shown).

pDCs were enriched from three female donors. In order to maximize pDC recovery, pDC yield was prioritized over pDC purity **(Figure S2)**. Cells were cultured at high density for 24 hours in the presence or absence of influenza H1N1 (A/PR/8/34). Following stimulation, cells were collected for viability assessment, culture media was cleared of cellular debris and media protein was precipitated for proteomic processing by LC-MS/MS. As has been previously established, pDC viability is enhanced *ex vivo* with stimulation (Thomas et al., 2014). Roughly 94% of the detected peptides corresponded to the exogenous media additives BSA, human transferrin and porcine trypsin (used in proteomic processing), reinforcing the imperative to minimize non-pDC derived media proteins *a priori* **(Figure S3A)**. The remainder of peptides were attributable to the cultured cells, identifying a total of 1,241 protein species, 819 of which were identified with high confidence, represented by 2 or more peptides **(Figure S3B-C)**. A list of identified proteins can be found in the supplementary appendix **(Tables S1 and S4)**.

Detected proteins which did not differ in abundance between stimulated and unstimulated conditions generally include exogenous media additives and human serum proteins not known to be produced by hematopoietic cells **(Figure S4A)**. The latter likely represent human serum proteins carried over through pDC isolation from whole blood. The protein secretome of unstimulated pDCs was enriched for proteins not known to be secreted, including proteins with various roles in the cytoskeleton and cellular metabolism **(Figure S4B)**. These proteins likely represent leakage from dying cells, and is consistent with the observed relative impairment of pDC viability in culture in the absence of stimulation.

Several proteins were present in a manner consistent with baseline secretory function of pDCs, however. These include pigment epithelium-derived factor/*SERPINF1*, alpha 2-antiplasmin/*SERPINF2*, and prostaglandin D2 synthase/*PTGDS*. These proteins are annotated as ‘secreted’ in the UniProt database and are recognized as enriched for expression in pDCs among circulating leukocytes in the Human Protein Atlas (http://www.proteinatlas.org/) (Thul et al., 2017). IL-16 is also enriched in the unstimulated condition; however, this cytokine is expressed by numerous cell types and thus may represent a contribution from contaminating non-pDC leukocytes in our cultures.

Stimulation with influenza virus is noted to induce the robust upregulation of numerous cytokines, including all 12 members of the IFN-α protein family **(Figure 1, Figure S4C-D)**. Other detected IFN-I members include IFN-β and IFN-ω. IFN-κ and IFN-ε were not detected. The IFN-III family was represented by IFN-λ1 and IFN-λ3. The sole IFN-II candidate, IFN-γ was observed at low levels and is likely is derived from contaminating cells, since this cytokine is not known to be produced by pDCs (see below scRNA-seq data for follow up). Other strongly induced cytokines include TNF-α and IL-6, and the chemokines CXCL9, CXCL10, CCL4 and CCL19. Several proteins encoded by secreted interferon-stimulated genes [ISGs] were detected, including ISG15 and SRGN. Notably, a number of proteins lacking a signal peptide and not annotated as ‘secreted’ elsewhere were overrepresented in the influenza stimulated group, including the proteases cathepsin c, cathepsin z, legumain, and the transcription factor IRF8. Finally, granzyme B is released from pDCs in response to influenza virus.

**Figure 1.**
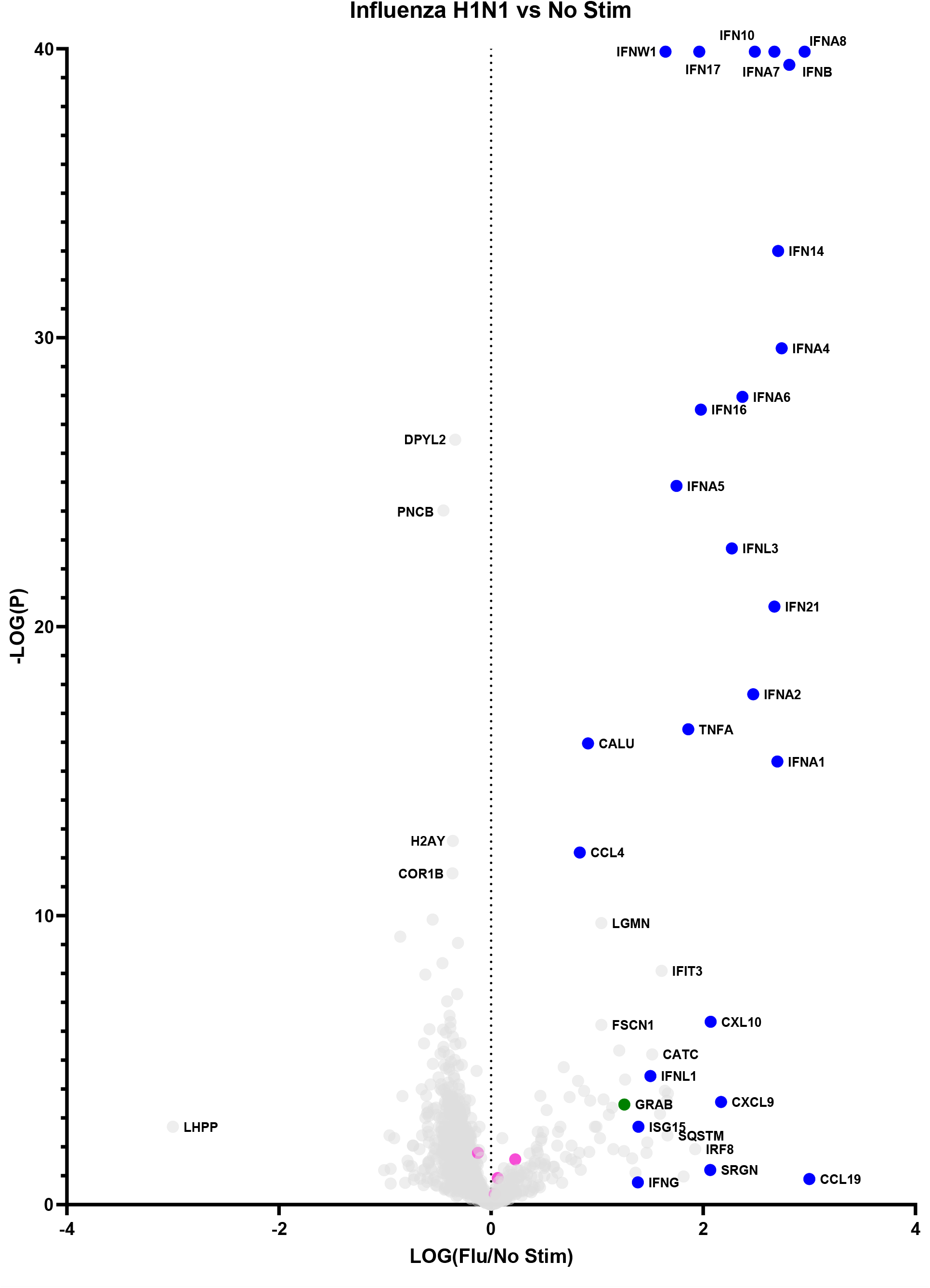
The pDC secretome: baseline vs influenza induced. Data points represent a single protein. Blue: cytokines/chemokines. Pink: exogenous proteins (bovine albumin, porcine trypsin, human transferrin). Green: granzyme B.

These data confirm IFN-α as the chief secreted product of stimulated pDCs. Moreover, they represent a more comprehensive and agnostic analysis of non-interferon secreted products, both upon stimulation, and in the unperturbed condition.

### pDC diversification in response to influenza virus mapped by scRNA-seq

Our above data are consistent with the well-established robust IFN-α secretion capabilities of pDCs. In our hands, we calculated the 24hr IFN-α production capacity of pDCs to 4.5-7pg/cell when averaged over a population of pure pDCs **(Figure S5)**. Making this value more astonishing is the reported observation that upon stimulation, only a minor fraction of pDCs stain positive for IFN-α when assayed by flow cytometry. This observation is limited by the fact that anti-IFN-α mAbs can only detect one to several of the IFN-α subtypes at a time, leaving open the possibility that while some pDCs stain positive for IFN-α, others account for the production of the uncaptured subtypes. Nevertheless, these reported findings suggest that what is traditionally thought of as a homogenous pool of pDCs may diverge in function following stimulation. pDC heterogeneity pre- and post-stimulation has been described. These studies were similarly limited, however, by the selection of mAbs used to delineate the pDC subpopulations. Therefore, we undertook a *de novo* characterization of pDC diversity, at baseline and in response to influenza virus using scRNA-seq.

pDCs were isolated from three genotyped female donors to near total purity **(Figure 2A)**. A third of pDCs from each donor was immediately partitioned for baseline pDC transcriptional profiling. The remainder were cultured with influenza virus for 6 and 24hrs; with cells partitioned at the close of each time point. To avoid batch effects, cells from the different donors were pooled upon partitioning by time point (*ex vivo*, 6hrs/flu, 24hrs/flu), with subsequent library preparation and Illumina sequencing steps shared among donors. Cell yields and sequencing output are reflected in **Table S2**. Single cells were assigned to their respective donors using Demuxlet (Kang et al., 2017). Donor-specific signals were controlled using Harmony (Korsunsky et al., 2019). Clusters were generated using Seurat (Stuart et al., 2019) **(Figure 2B)**. The data reveal that the cells segregate predominantly by time point **(Figure 2C)**, with clusters evident within each major condition. Coloring cells by donor reveals that the donor cells are well-interspersed among the clusters **(Figure 2D)**, negating the possibility that donor-specific characteristics are main drivers of cell diversity. 18 clusters are identified by Seurat. The top differentiating genes among the clusters are listed in the supplementary appendix **(Table S3)**.

**Figure 2:**
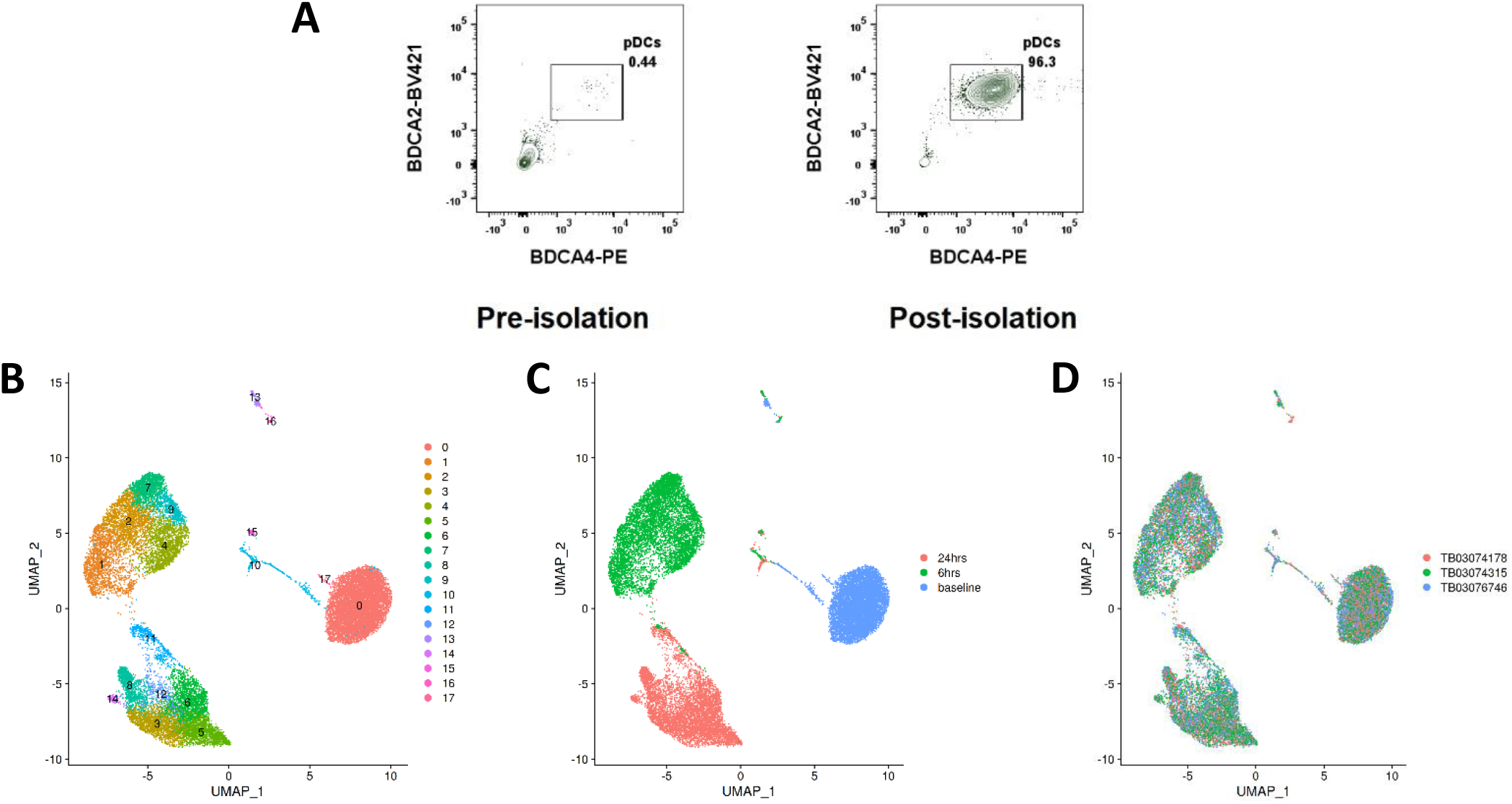
pDC heterogeneity determined by scRNA-seq. A) Representative pre- and post-pDC isolation purities. Data represent ungated PBMCs stained for the pDC markers BDCA4 and BDCA2. B) Cell clusters identified by Seurat. C) Cells colored by time point at partitioning. D) Cells colored by donor.

Of the clusters, clusters 13 and 16 represent minor B and T cell contamination of the pDC cultures, respectively. Cluster 17 represents AXL^+^ pDCs, a recently described dendritic cell with an intermediate phenotype between pDCs and cDCs (Villani et al., 2017). The remaining clusters represent bona fide pDCs with distinct transcriptional profiles.

Clusters 1 and 11, in the 6hr and 24hr influenza virus-treated conditions, respectively, represent the IFN producing cells **(Figure 3A)**. These clusters mapped with close proximity to each other across the two time points. Notably, these clusters expressed the totality of IFN-alpha genes and other IFN-Is produced by pDCs **(Figure 3B)**. TCF4, the transcription factor instructing pDC identity, is depleted in these clusters, whereas ID2, the transcription factor opposing pDC identity, is upregulated (see next paragraph). Moreover, the classical pDC markers NRP1, CLEC4C and CD123 are depleted in the IFN producing cells **(Figure 4A)**. These data confirm that among a population of pDCs treated with the same stimulus, only a minor portion mature into the IFN producing population. These cells appear to adopt a dedicated role, losing canonical transcriptional features of pDCs. Alternative surface markers suggested by the data for the IFN producers include IL18RAP, LILRA5 and LILRB2, among others **(Figure 4B)**.

**Figure 3:**
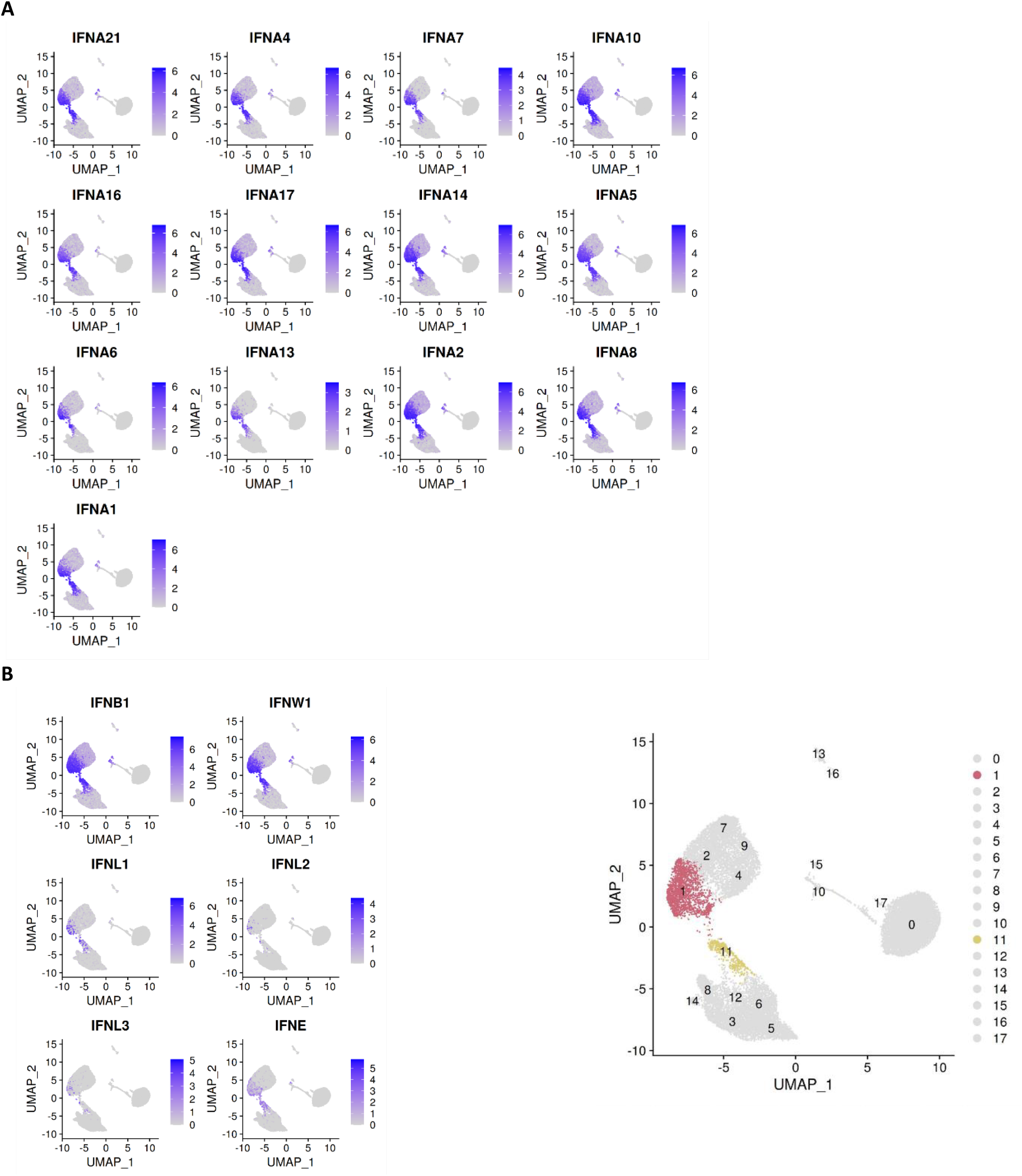
Clusters colored by IFN-I gene expression. A) Cells colored by IFN-α subtype gene expression. B) Cells colored by non-IFN-α type I interferon gene expression. Bottom right panel: highlight of clusters 1 and 11 identified by Seurat.

**Figure 4:**
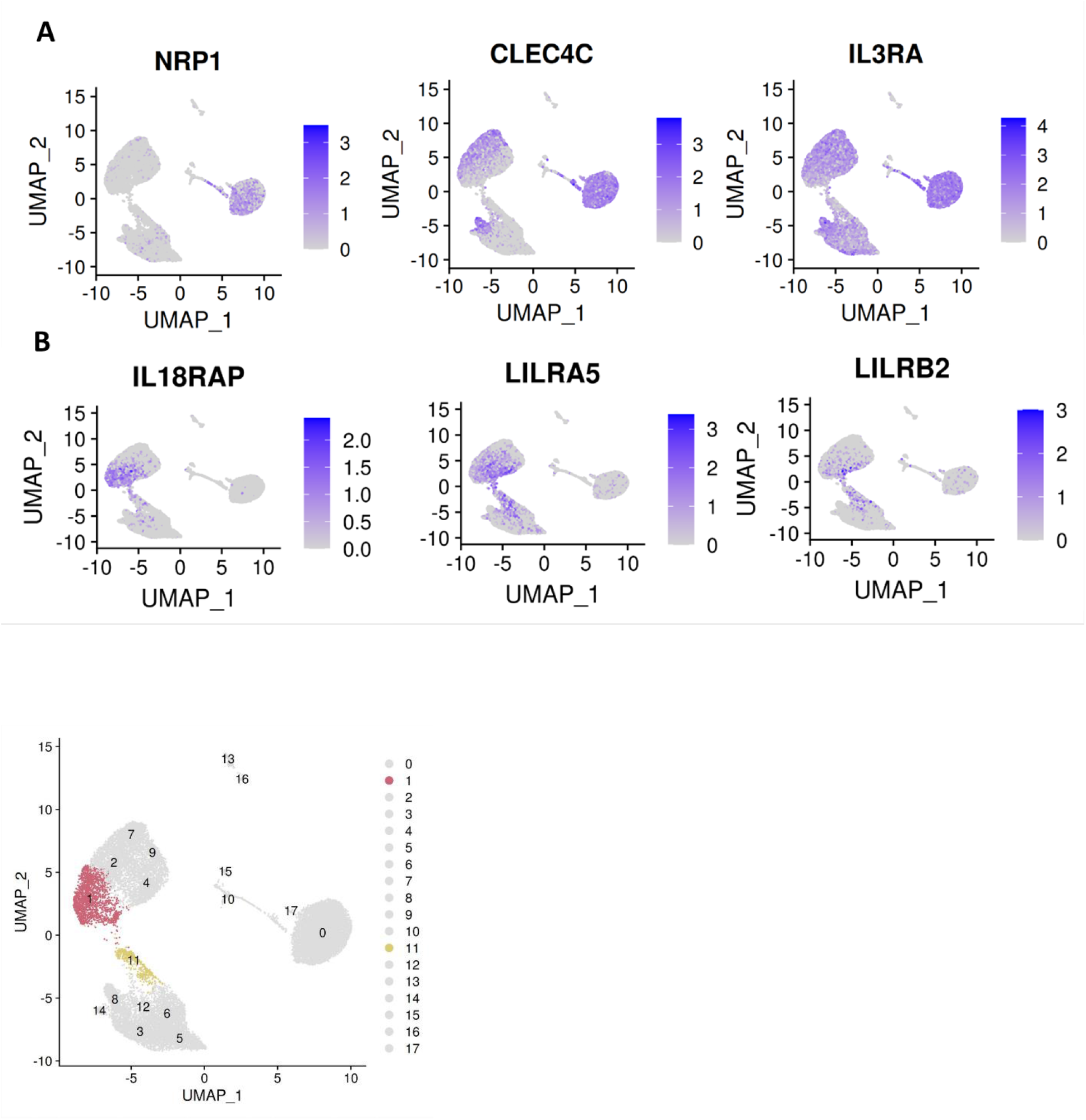
Dropout of classical pDC markers and novel surface markers on the IFN-α^+^ cells. A) Dropout of the classical pDC surface antigens NRP1 [BDCA4], CLEC4C [BDCA2] and IL3RA [CD123] in the IFN-producers. B) Novel suggested surface markers in the IFN-producers. Bottom panel: highlight of clusters 1 and 11 identified by Seurat.

Another notable phenomenon within the single cell data was the dynamism of expression of the transcription factors TCF4 and ID2. **(Figure 5A)**. TCF4 dictates pDC identity (Cisse et al., 2008), and its continuous expression is required for the maintenance of this identity (Ghosh et al., 2010). ID2 is an antagonist of TCF4 which instructs a cDC profile. All pDCs *ex vivo* were TCF4^+^ID2^-^, whereas upon stimulation, expression of these transcription factors varied widely by cluster. Cluster 5 is particularly prominent for loss of TCF4 and gain of ID2. This change is accompanied by upregulation of the T cell co-stimulatory markers CD80/CD86, cDC markers FSCN1 and lysozyme, and the secondary lymphoid organ T cell zone homing receptor CCR7 **(Figure 5B)**. These observations suggest that cluster 5 is a cDC/APC-like derivative of pDCs whose principal role is in antigen presentation to T cells.

**Figure 5.**
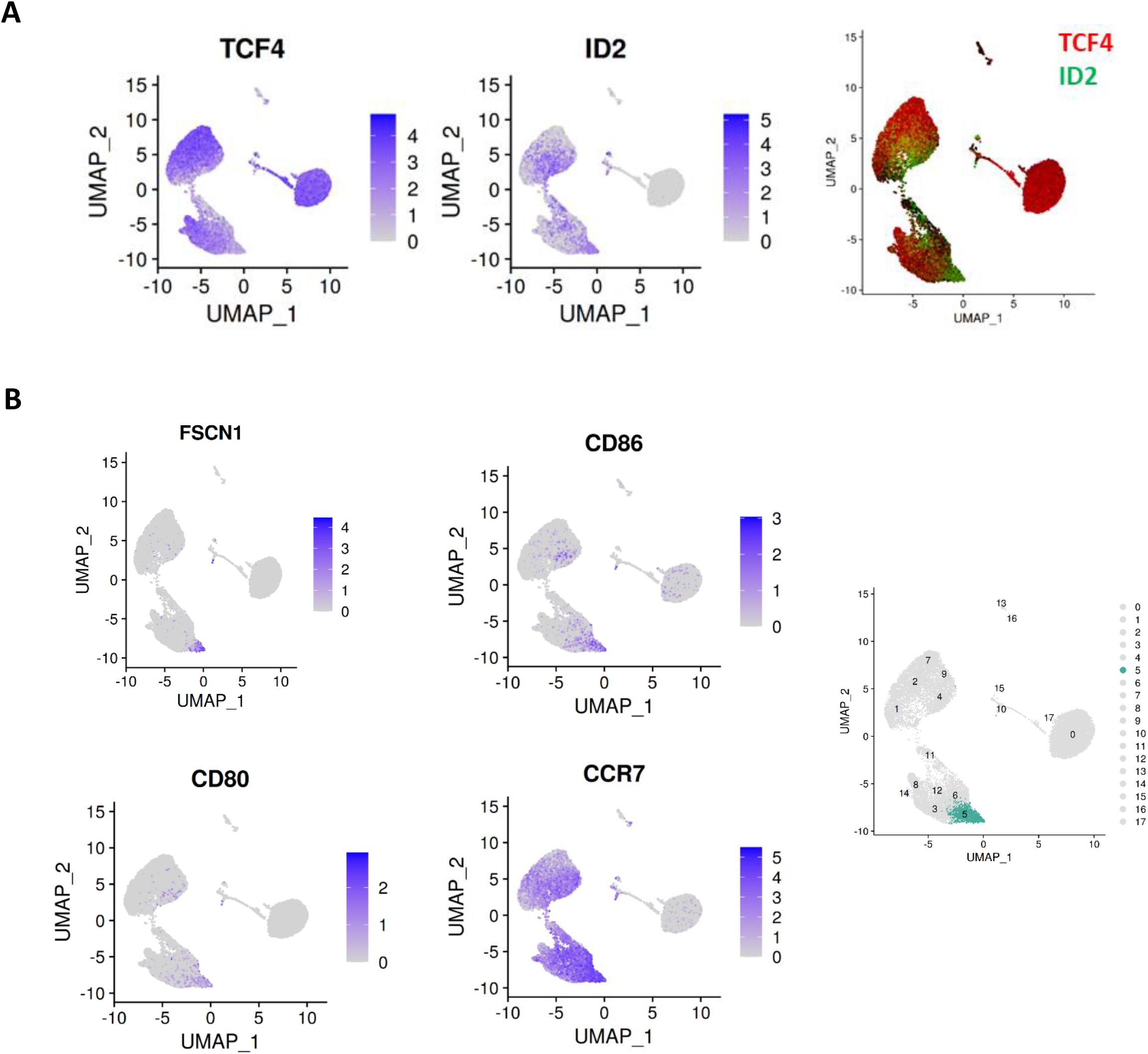
Dynamic regulation of master transcription factors across pDC clusters. A) An inverse relationship is observed between the pDC master regulator TCF4 and the TCF4 antagonist ID2. The cluster 1/11 isthmus is notable for loss of TCF4 and gain of ID2. B) Cluster 5 (highlighted in the right panel) is notable for upregulation of ID2, and the cDC/APC markers CD86, CD80, FSCN1 and CCR7.

Cluster 8 represents a group of cells retaining classical pDC features even following 24hr culture with influenza virus. These markers include CLEC4C and CD123, as well as pDC specific genes such as SCAMP5, GZMB, TCF4 and IRF7 **(Figure S6)**. Whether this cluster represents a group of cells resistant to stimulation, or a ‘return to baseline’ phenotype, remains to be determined.

Finally, pDC heterogeneity is evident at baseline **(Figure 6)**. On fine grain clustering, 9 subclusters are identified within the unstimulated group partitioned immediately *ex vivo*. Importantly, the classical pDC markers NRP1 and CLEC4C are evenly distributed among these clusters, as is the master pDC regulator TCF4. ID2 is absent. PTGDS, TCL1A, TCL1B and IGLC2 clearly distinguish these clusters. Whether this preexisting heterogeneity informs maturation into IFN producers or other phenotypes following stimulation is the subject of future investigations. Taken together, this dataset represents a multidimensional, discovery-led approach revealing pDC transcriptional diversity at baseline and further upon stimulation with influenza virus.

**Figure 6:**
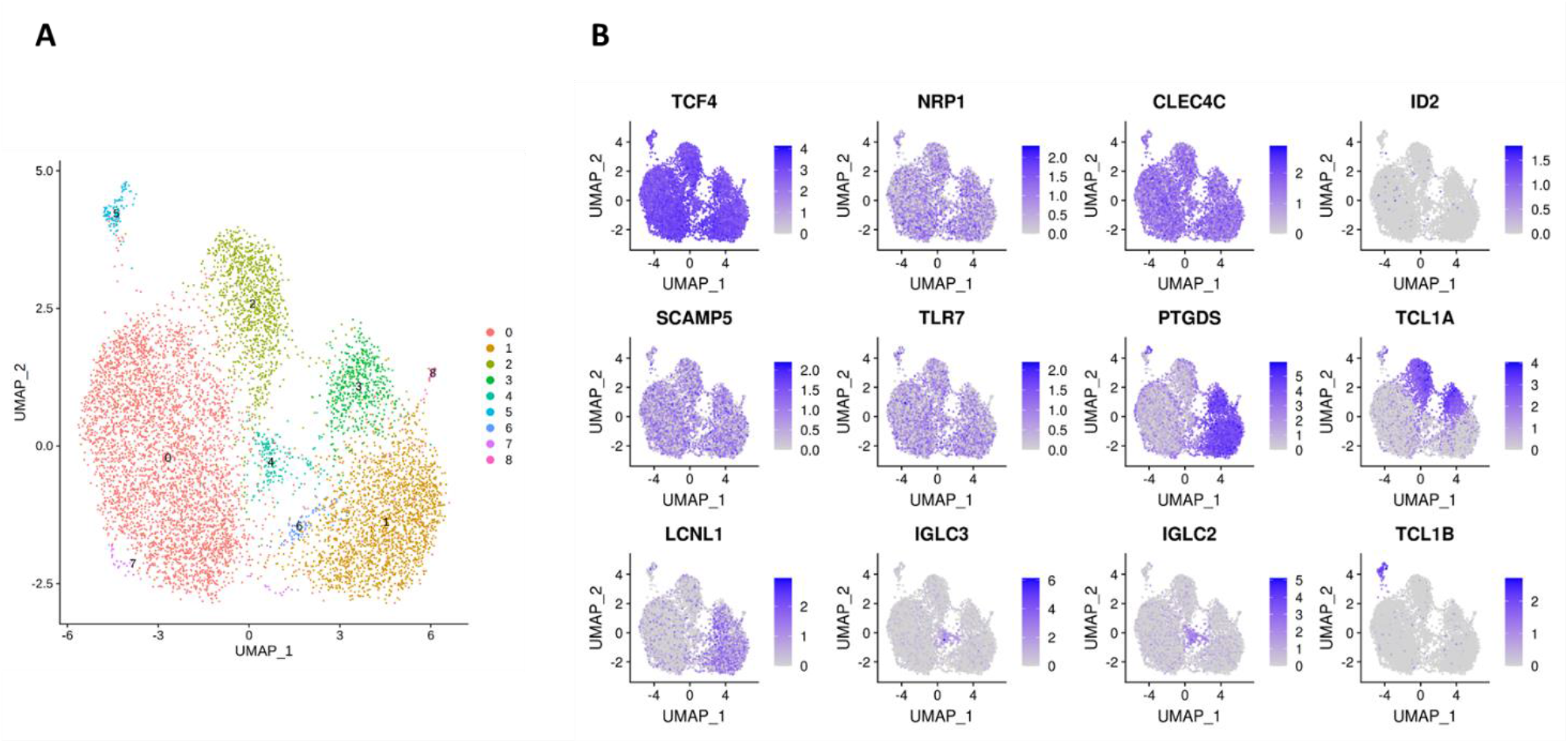
pDC heterogeneity at baseline. Cluster 0 is further divided into 9 subclusters by Seurat, highlighting preexisting pDC heterogeneity as the cells are isolated ex vivo. TCF4, NRP1 and CLEC4C are evenly distributed within these subclusters, as expected. ID2 is absent. pDC enriched genes SCAMP5 and TLR7 are also uniformly expressed, whereas PTGDS, TCL1A, LCLN1, IGLC3, IGLC2 and TCL1B distinguish various subclusters.

### Integration of LC-MS/MS and scRNA-seq datasets

The scRNA-seq data also allow for a more complete understanding of the secretome data discussed previously. As mentioned, pDCs in latter experiments were only ~ 60% pure, versus near total purity in the scRNA-seq dataset. Many identified cytokines, such as IFN-I and IFN-III, are readily attributable to pDCs. Other identified cytokines and chemokines, such as IL-16, CXCL9, CCL4 and CCL9 may or may not be pDC-derived. As the secretome and scRNA-seq experiments similarly involved pDC stimulation with influenza virus over a course of 24hrs, the scRNA-seq data may be leveraged to determine whether pDCs are in fact contributing the identified proteins in the secretome dataset. As shown in **Figure 7**, IL-16, CCL4 and CXCL10 are confirmed to be pDC-derived, whereas IFNG is confirmed to come from non-pDCs. Interestingly, IL16 is expressed predominantly among cells of the *ex vivo* time point, and is lost upon stimulation, confirming its status as a member of the ‘baseline’ pDC secretome discussed above.

**Figure 7.**
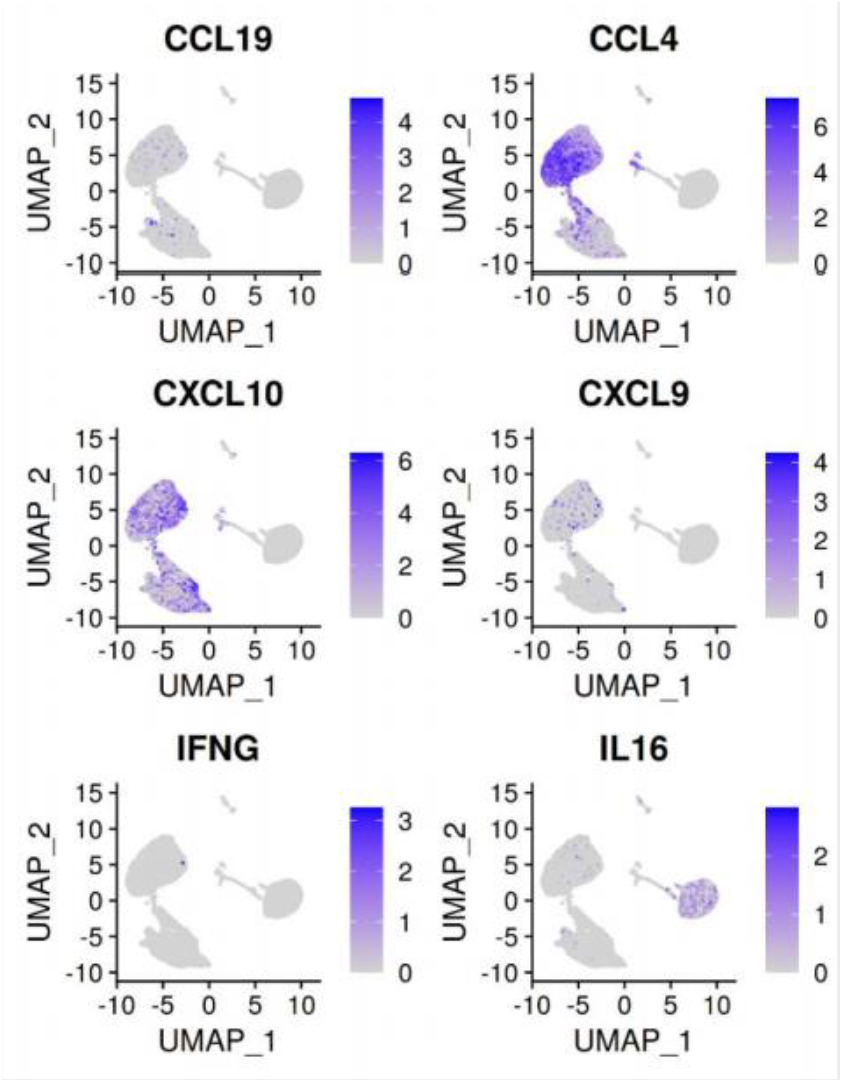
Cytokines/chemokines identified by LC-MS/MS and corresponding transcript presence in the scRNA-seq dataset. CXCL10 is a known pDC chemokine and is upregulated in pDCs upon flu virus treatment. CCL4 can be attributed to pDCs on the basis of upregulation with flu virus seen here. IFNG is not known to be a pDC cytokine, the absence of IFNG in the scRNA-seq dataset confirms the non-pDC origin of this cytokine in the secretome experiment. IL-16 is identified in the secretome data as enriched in the unstimulated condition. This finding is supported by the observation that ex vivo pDCs express IL16, whereas flu virus treated pDCs do not.

### *In vitro* validation of pDC markers suggested by scRNA-seq

The above findings merit validation and further exploration in the appropriate experimental and clinical contexts. For example, the finding that IFN-α^+^ pDCs gain a novel surface marker profile can be validated by flow cytometry. LILRB2 is noted to be enriched for expression in the IFN-α producers. Gating on the IFN-α^+^ cells in a population of pDCs stimulated with influenza virus demonstrates significant surface staining for LILRB2 relative to the IFN-α^-^ cells **(Figure 8A)**. Similarly, other clusters can be further explored. Cluster 5, shown to adopt a ‘cDC-like’ transcriptional profile, is noted to upregulate expression of CD80, CD86, and CCR7. Cells of this cluster can be detected using mAbs against these respective markers after overnight culture with influenza virus, whereas cells cultured in the absence of virus do not stain for any of these markers **(Figure 8B)**. Future validation in samples from patients with SLE, influenza or COVID-19 will be necessary to follow up on the clinical relevance of these data.

**Figure 8.**
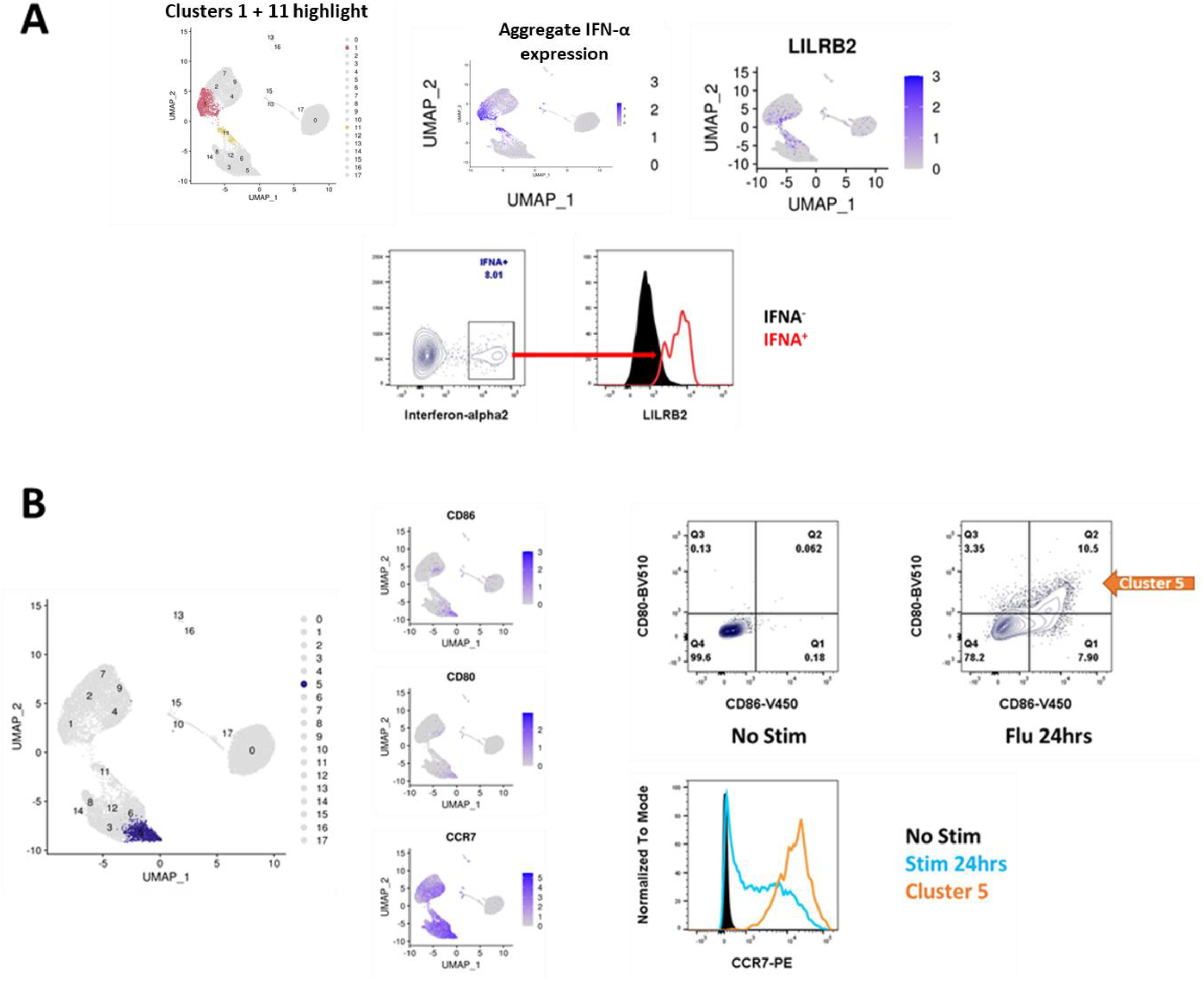
Validation of surface markers suggested by scRNA-seq. A) pDCs in the IFN-α^+^ cluster lose canonical pDC markers and adopt new ones (see also, Figure 6), including IL18RAP, LILRA5 and LILRB2. Upon stimulation with influenza virus, the population of IFN-α^+^ cells stains positive for LILRB2 whereas IFN-α^-^ cells are negative for LILRB2. B) Cluster 5 is notable for the adoption of a cDC like state, including upregulation of ID2 (Figure 9), CD80, CD86 and CCR7. Stimulation of pDCs with influenza virus produces a population of cells which stain for CD80 and CD86, these cells also upregulate expression of CCR7 at the protein level as determined by flow cytometry.

## Discussion

### pDC secretory activity

pDCs were definitively identified as the ‘natural-IFN producing cell’ in the circulation, and IFN-I has been the principle focus of pDC investigations since. Besides IFN-I, traditional approaches have shown that pDCs also produce substantial amounts of IFN-III, TNF-alpha, IL-6 and CXCL10, typically using single-analyte ELISAs and often in response to synthetic stimuli. An early attempt towards characterizing the pDC secretome utilized a 12-plex Luminex assay to show that pDCs also produce CCL2, CCL3, CCL4, CCL5, IL-1RA and IL-8 in response to stimulation with influenza virus or ODN-2216 (Decalf et al., 2007). Another group attempted to characterize DC secretion via 2D-PAGE followed by shotgun proteomics from monocyte-differentiated DCs (Gundacker et al., 2009). The resemblance of in vitro differentiated DCs to primary DCs is questionable, however, as even the most advanced techniques produce cells transcriptionally distant from their natural counterparts (Martin Jakobsen, personal communication).

To our knowledge, a comprehensive and agnostic study of the pDC secretome has not been previously attempted. Moreover, secreted products of unperturbed pDCs have not been systematically catalogued. To accomplish these goals, we subjected media protein preparations from high density pDC cultures to LC-MS/MS analysis. To increase the odds of detecting pDC-derived proteins, we used the minimal amount of exogenous protein possible. In addition, we did not enrich the pDCs to total purity, as this would inevitably reduce the yield of pDCs. The finding that bovine albumin was the dominant species in the culture media was expected, the extent of which corroborated the need to minimize the presence of media protein additive. Methods exist to deplete albumin from sample preparations; we elected not to use these methods for risk of introducing identification bias (BSA depletion may also deplete non-specifically).

We performed a pilot secretome analysis on unstimulated and ODN-2216 stimulated pDCs from 3 donors, identifying 293 proteins (data not shown), serving as proof-of-concept. Our follow-up analysis, presented here, using unstimulated and influenza virus stimulated pDCs from an additional 3 donors netted 1,241 protein identifications. Our approach to analyzing the list of identified proteins prioritized species annotated as ‘secreted’ in the Uniprot database. To ascribe pDCs as the source of a particular protein in the unstimulated condition, we referred to the Human Protein Atlas to determine the degree of pDC-specific expression. Secreted proteins identified in the influenza virus treated conditions were ascribed to pDCs on the basis of historical association or upregulation in the scRNA-seq data, which also used influenza virus as the stimulus.

Our results suggest the presence of a ‘baseline’ pDC secretome. Interleukin-16 (IL-16/*IL16*), pigment epithelium-derived factor (PEDF/SERPINF1), alpha-2-antiplasmin (A2AP/SERPINF2) and prostaglandin-D2-synthase (PGD2/PTGDS) are identified as top candidates for pDC-specific secreted proteins in unperturbed conditions. The biological significance of PEDF and A2AP secretion by pDCs has not been explored. PGD2 catalyzes the formation of the arachidonic acid metabolite prostaglandin D2, a neuromodulator and potent vasodilator. A single report (Shimura 2010) describes pDCs as a hematopoietic source of PGD2 but does not further examine the role of PGD2 from these cells in particular (Shimura et al., 2010). Other baseline pDC secreted products were also identified, including several additional members of the SERPIN family, although pDCs are likely not the sole source of these proteins among leukocytes. For example, IL-16, a T cell chemoattractant, was identified in the secretome data as enriched in the unstimulated condition. Our scRNA-seq data points to expression of IL-16 in *ex vivo* pDCs which is lost upon stimulation. The Protein Atlas suggests that non-pDCs serve as IL-16 producers as well. Together, these proteins represent underexplored baseline pDC secretory activity.

In the induced secretome, IFN-I was clearly the dominant category of secreted protein, including all 12 IFN-α subtypes, IFN-β and IFN-ω. The IFN-III category was represented via IFN-λ2 and 3. TNF-α, IL-6 and CXCL10—known pDC cytokines, were also well represented. Cytokines less commonly associated with pDCs, such as CCL4 and IFN-gamma, were respectively ruled-in and ruled-out as pDC-sourced on the basis of scRNA-seq data. Not all cytokines reported by Decalf et al. were identified, however. CCL3 and CCL5, for example, were absent in the secretome dataset, but highly upregulated in the IFN-producers in the scRNA-seq dataset. This likely indicates that our secretome analysis missed important identifications, reflecting a technical limitation as currently performed. On the other hand, the cytokines CCL2 and IL-1RA were absent in both secretome and scRNA-seq datasets, suggesting non-pDC contamination as the source of these proteins in Decalf et al.

Granzymes are conventionally thought of in terms of cytotoxic lymphocyte degranulation and represent a major antiviral tool of adaptive immunity (Chowdhury and Lieberman, 2008). The Protein Atlas illustrates preferential expression of granzymes A, H, K and M in CD8 T cells and NK cells, as expected. Granzyme B, however, is by far most highly expressed in pDCs **(Figure S7)**. Granzyme B is enriched in the culture media of influenza stimulated pDCs, as observed in the secretome analysis. In the scRNA-seq dataset, GZMB expression is noted at baseline and is lost upon stimulation, counter to the protein level observation. This finding is consistent with granzyme release via fusion of preformed granules with the plasma membrane, a process distinct from IFN secretion, which is induced *de novo* at the transcript level for rapid transit through the secretory pathway.

Overall, the induced pDC secretome confirms the robust reactionary nature of these cells, whose chief product is IFN-α, and argues against baseline hypersecretory function akin to the namesake plasma cells. Other secreted protein products represent a more diverse picture of virus-induced pDC activity. A striking component of the induced secretome consisted of proteins not annotated as secreted yet which are pDC-specific, including IRF8, legumain and various cathepsins. Leakage from dying cells is unlikely to explain the enrichment of these proteins in the media of stimulated cells, as stimulation enhances pDC survival in vitro. We hypothesize that pDC stimulation results not only in secretion through the classical ER-Golgi pathway, but also results in the release of protein-loaded exosomes. These exosomes may serve as an alternative method for transferring antiviral programs to recipient cells. Recently it was demonstrated that pDCs form ‘interferogenic synapses’ with viral infected cells (Assil et al., 2019). Communication of an antiviral program via transcription factor and protease loaded exosomes in a paracrine fashion is an exciting avenue for future exploration. This work marks the first agnostic description of the primary human pDC secretome, at baseline, and in response to a physiologic stimulus. Our data support multiple mechanisms of regulated protein release from pDCs, including the classical secretory pathway, degranulation and, putatively, exosome formation.

### Transcriptional diversity of pDCs

pDC heterogeneity pre- and post-stimulation has been described. A CD2^hi^Lysozyme^+^ population of pDCs was reported to possess enhanced T cell activation properties (Matsui et al., 2009). A similar report describes CD2^hi^CD5^+^CD81^+^ pDCs as potent inducers of T cell proliferation (Zhang et al., 2017). That a minority of pDCs produce IFN-α upon stimulation has long been known. The evidence for this, however, typically involves intracellular antibody staining for IFN-α towards flow cytometric analyses. As there are 12 IFN-α subtypes, no single IFN-α antibody could detect all subtypes simultaneously. As such, it remained possible that a minority of pDCs produced particular subtypes of IFN-α, with other pDCs producing the remaining subtypes. Alculumbre et al. report a detailed analysis of pDC diversification following stimulation with influenza virus (Alculumbre et al., 2018). Their results show that upon treatment with influenza virus, pDCs diversify along a 2-parameter, CD80/PDL1 axis. CD80^+^PD-L1^-^ cells adopt APC-like functions with dendritic morphology, whereas CD80^-^PD-L1^+^ cells secrete interferon and retain plasmacytoid morphology. These results were later recapitulated using SARS-CoV-2 as the stimulus (Onodi et al., 2021). Using mass cytometry, it was further shown that human pDCs mount distinct, heterogenous responses dependent on the stimulus (Yun et al., 2021).

To better understand pDC diversity pre- and post-stimulation in a highly multidimensional format, we analyzed the transcriptomes of pDCs from 3 donors immediately *ex vivo*, upon 6hrs of influenza virus stimulation and upon 24hrs of influenza virus stimulation at the single-cell level. To our knowledge, we are the first to demonstrate that a single population of IFN-producers is responsible not only for the totality of IFN-α production, but for the other IFN-I family members, IFN-III, and the majority of induced cytokines as well. This population represents roughly 24% of cells at the 6hr time point and 5.5% of cells at the 24hr time point. These clusters are notable for dropout of classical pDC markers, including BDCA4, BDCA2, CD123 and ILT7. pDCs have been reported to diminish in the circulation of SLE patients with active disease (Cederblad et al., 1998; Hancharou et al., 2012), patient affected by pandemic influenza (Lichtner et al., 2011), and in patients with severe COVID-19 (Peruzzi et al., 2020; Ren et al., 2021; Wilk et al., 2020). This has commonly been interpreted to indicate that pDCs in these patients infiltrate affected tissues, decreasing their presence in the circulation. pDCs have been shown to diminish from the bronchoalveolar fluid of COVID-19 patients(Chua et al., 2020), however, challenging this theory. Our findings suggest an alternative explanation: pDCs appear to be reduced in these conditions because they are no longer detectable using the standard pDC surface antigens. We suggest alternative surface markers for the detection of IFN-producing, activated pDCs. These markers can then be applied to characterize pDCs in the relevant clinical contexts.

Another notable phenomenon within the single cell data was the dynamism of TCF4/ID2 expression. TCF4 dictates pDC identity (Cisse et al., 2008), and its continuous expression is required for the maintenance of this identity (Ghosh et al., 2010). ID2 is an antagonist of TCF4 which instructs a cDC profile. All pDCs *ex vivo* were TCF4^+^ID2^-^, whereas upon stimulation, expression of these transcription factors varied widely by cluster. Cluster 5 is particularly prominent for loss of TCF4 and gain of ID2. This change is accompanied by upregulation of the T cell co-stimulatory markers CD80/CD86, cDC marker FSCN1, and the secondary lymphoid organ T cell zone homing receptor CCR7. These observations suggest that cluster 5 is a cDC/APC like derivative of pDCs whose principal role is in antigen presentation to T cells. A small population, cluster 8, is observed to maintain the panel of pDC specifying transcription factors, including TCF4, SPIB, and IRF8, along with pDC-typical genes including IRF7, GZMB, SCAMP5 and pDC surface markers BDCA2, ILT7 and CD123. Whether the presence of this cluster at 24hrs represents cells unaffected by stimulation or cells which lost and subsequently recovered pDC identity is not clear. It is apparent, nonetheless, that pDC identity reflects a continuous rebalancing of master regulators.

Preexisting heterogeneity is also evident from our data, with 9 baseline subclusters identified. Whether this heterogeneity determines the fate of a particular cell upon influenza virus stimulation remains to be determined. Unfortunately, the genes discriminating these clusters are not expressed at the cell surface, precluding the sorting of live cells towards fate-determination assays. Notably, CD2, CD5 and CD81, markers previously reported to define baseline pDC heterogeneity, did not discriminate these baseline pDC clusters. Rather, these markers were enriched in cluster 17, AXL^+^ pDCs. This population was recently identified as cells with an intermediate phenotype between pDCs and cDCs (Villani et al., 2017), and likely explains the enhanced T cell proliferation capacity of the cells reported earlier (Matsui et al., 2009; Zhang et al., 2017).

In summary, we have applied novel techniques to the study of pDC function and identity. Our results shed new light on pDC secretory activity at baseline and upon stimulation. Altogether, these findings suggest a need to address new aspects of pDC biology, which do not necessarily center on cell activation or IFN-I secretion. Furthermore, the pDC maturation trajectories described here bear clinical relevance in terms of use as biomarkers for disease states, towards pDC-centric immunotherapies and towards TLR7/9-based vaccine adjuvants.

## Materials and Methods

### pDC isolation and culture

Human donors were recruited from the Genotype and Phenotype Registry [GaP], a living biobank of genotyped individuals who have volunteered for recall on the basis of their genotype (Gregersen et al., 2015). Fresh whole blood was drawn from these donors into sodium-heparin vacutainers. PBMCs were extracted over Ficoll. pDCs were isolated from PBMCs using Miltenyi Plasmacytoid Dendritic Cell Isolation Kit II, human [Cat# 130-092-207] or EasySep Human Plasmacytoid DC Enrichment Kit [Cat # 19062]. pDC purity was determined by flow cytometry against BDCA2 and/or BDCA4.

For the secretome experiments, pDCs were cultured in a 1:1 mixture of Advanced RPMI-1640 [Gibco 12633012] and RPMI-1640 [Gibco 11875119], without serum supplementation in ultra-low protein binding 24-well plates. Where indicated, influenza virus H1N1 A/PR/8/34 [Advanced Biotechnologies] at 0.4 HAU/mL was used for stimulation.

For the scRNA-seq experiments, pDCs were cultured in Advanced RPMI-1640 [Gibco 12633012] supplemented with 5% FCS, with the same concentration of influenza virus as in the secretome experiment.

### Sample Preparation for Proteomics

Methods are summarized here with additional details described previously (Wobma et al., 2018). Secreted proteins from three biological replicates of each treatment were precipitated with chloroform/methanol, reduced, alkylated, and digested with trypsin. Samples were desalted using Nestgroup C18 Macrospin columns. Eluted peptides were lyophilized, and redissolved in 3% acetonitrile, 0.1% formic acid.

### Liquid Chromatography and Mass Spectrometry

Separations were performed with an Ultimate 3000 RSLCNano (Thermo Scientific) and a 75 μm ID x 50 cm Acclaim PepMap reversed phase C18, 2 µm particle size column. Flow rate was 300 nL/min with an acetonitrile/formic acid gradient and a column temperature of 40 °C with details as described previously(Wobma et al., 2018).

Mass spectra were collected with a Q Exactive HF mass spectrometer (Thermo Scientific) in positive ion mode using data-dependent acquisition (DDA). Settings included top 15, resolution 120,000 for MS, 15,000 for MS/MS, dynamic exclusion for 20 s and maximum injection time 30 ms for MS and 100 ms for MS/MS. NCE was 28.0, source 2.2kV and 250 °C and S-Lens RF at 55%.

Data processing was done with Elucidator software (Protein Expression Data Analysis System (Ver. 4.0.0.ror2.31, Ceiba Solutions/ PerkinElmer) for label-free quantitation based on comparison of MS1 feature volume. PeptideTeller was set to 0.01 predicted error. Protein identifications were obtained by searching against a database from UniProtKB Release 2020_01, 26-Feb-2020 with 43,392 sequences using Mascot server V. 2.5.1. The database included human reference proteome UP000005640 of reviewed canonical sequences with isoforms as well as the Influenza A virus (strain A/Puerto Rico/8/1934 H1N1) reference proteome UP000170967 with 25 sequences. Also included were porcine trypsin, bovine serum albumin, sheep keratin and other standard contaminants. Expression ratios reported were calculated by the software at the peptide level rather the protein level.

All mass spectrometry raw files are deposited in at the MassIVE public repository (https://massive.ucsd.edu/)

### Single cell RNA-seq

For the scRNA-seq experiments, isolated pDCs were cultured in Advanced RPMI-1640 [Gibco 12633012] supplemented with 5% FCS, with the same concentration of influenza virus as in the secretome experiment where indicated. pDCs were counted, pooled at equal ratios from the 3 donors for each time condition, washed, resuspended in PBS/0.4% BSA at a final concentration of 1000 cells/ul and loaded onto the 10X Chromium Controller (10X Genomics). GEMs (Gel Bead-in Emulsion) were generated using Chromium Next GEM Single Cell 3’ GEM, Library & Gel Bead Kit v3.1, following the manufacturer’s recommendations. Single cell libraries were assessed and quantitated using a High Sensitivity DNA chip (Agilent). Libraries were loaded onto a High Output Cartridge v2 (PN # 15057931) at 1.8pM and run on a NextSeq 500 sequencer (Illumina). The raw data from the sequencer was demultiplexed into reads along with cell and unique molecular identified [UMI] barcodes which were then aligned to GrCH38 human reference and quantified via CellRangerV3 from 10x Genomics. For each condition, cells belonging to each donor were determined using Demuxlet (Kang et al., 2017). Further downstream analysis was done using Seurat (Stuart et al., 2019). Doublets and ambiguous cells identified by Demuxlet were removed along with any cell with < 200 unique genes, any cell expressing > 20% mitochondrial genes and any gene not identified in at least 3 cells. Clusters were identified using Seurat’s SCT transform analysis using 30 PCs with time point as an effect being removed using Harmony (Korsunsky et al., 2019). Statistical analysis and figure plotting was also done using R [https://www.r-project.org/] and Tidyverse (Wickham et al., 2019).

### Cytokine ELISAs and Flow Cytometry

Where indicated, IFN-α was quantified from culture media using a pan-subtype ELISA (MABTECH 3425-1H-6).

Intracellular staining for IFN-α2 was accomplished using BD Cytofix/Cytoperm in accordance with the manufacturer’s protocol, along with a monoclonal anti-IFN antibody (Abcam EPR19074). LILRB2 (564345), BDCA4 (565951) and BDCA2 (566427) antibodies were obtained from BD. Cells were acquired on the BD Fortessa flow cytometer and data were analyzed using FlowJo v10.

## Acknowledgments

We thank our blood donors for their contribution to this work. Gila Klein, Margaret DeFranco, Kristine Elmaliki and Dorean Flores coordinated GaP donor recall and blood draws. Drs. Sun Jung Kim, Betsy Barnes and Patricia Fitzgerald-Bocarsly provided helpful discussions. Christopher Colon and Barrett Waling assisted with flow cytometry. We appreciate help from Shahar Goeta for technical support with the proteomics work. LMB gratefully acknowledge the funding support from NYSTEM contract C029159 for the mass spectrometer (New York State Stem Cell Science Board, LMB). Matching funds (LMB) from Columbia University through the Department of Biological Sciences, Dean of Science, Executive Vice President for Arts and Sciences, Executive Vice President for Research, Department of Biomedical Engineering, Department of Medicine, the Fu Foundation School Engineering & Applied Science and the Columbia Stem Cell Initiative are also appreciated.

## Author contributions

MHG and PKG conceived the study. MHG, KRS, LMB and PKG designed the experiments. MHG, HK, EW, and JKC performed the experiments. MHG, AJS, EW, JKC, LMB, KRS and PKG analyzed the data. MHG, AJS, KRS, LMB and PKG wrote the paper. All authors reviewed and approved the final manuscript.

## References

Alculumbre, S.G., V. Saint-Andre, J. Di Domizio, P. Vargas, P. Sirven, P. Bost, M. Maurin, P. Maiuri, M. Wery, M.S. Roman, L. Savey, M. Touzot, B. Terrier, D. Saadoun, C. Conrad, M. Gilliet, A. Morillon, and V. Soumelis. 2018. Diversification of human plasmacytoid predendritic cells in response to a single stimulus. Nat Immunol 19:63–75.

Assil, S., S. Coleon, C. Dong, E. Decembre, L. Sherry, O. Allatif, B. Webster, and M. Dreux. 2019. Plasmacytoid Dendritic Cells and Infected Cells Form an Interferogenic Synapse Required for Antiviral Responses. Cell Host Microbe 25:730–745 e736.

Baechler, E.C., F.M. Batliwalla, G. Karypis, P.M. Gaffney, W.A. Ortmann, K.J. Espe, K.B. Shark, W.J. Grande, K.M. Hughes, V. Kapur, P.K. Gregersen, and T.W. Behrens. 2003. Interferon-inducible gene expression signature in peripheral blood cells of patients with severe lupus. Proceedings of the National Academy of Sciences 100:2610–2615.

Cederblad, B., S. Blomberg, H. Vallin, A. Perers, G.V. Alm, and L. Rönnblom. 1998. Patients with Systemic Lupus Erythematosus have Reduced Numbers of Circulating Natural Interferon-α-Producing Cells. Journal of Autoimmunity 11:465–470.

Chua, R.L., S. Lukassen, S. Trump, B.P. Hennig, D. Wendisch, F. Pott, O. Debnath, L. Thurmann, F. Kurth, M.T. Volker, J. Kazmierski, B. Timmermann, S. Twardziok, S. Schneider, F. Machleidt, H. Muller-Redetzky, M. Maier, A. Krannich, S. Schmidt, F. Balzer, J. Liebig, J. Loske, N. Suttorp, J. Eils, N. Ishaque, U.G. Liebert, C. von Kalle, A. Hocke, M. Witzenrath, C. Goffinet, C. Drosten, S. Laudi, I. Lehmann, C. Conrad, L.E. Sander, and R. Eils. 2020. COVID-19 severity correlates with airway epithelium-immune cell interactions identified by single-cell analysis. Nat Biotechnol 38:970–979.

Cisse, B., M.L. Caton, M. Lehner, T. Maeda, S. Scheu, R. Locksley, D. Holmberg, C. Zweier, N.S. den Hollander, S.G. Kant, W. Holter, A. Rauch, Y. Zhuang, and B. Reizis. 2008. Transcription factor E2-2 is an essential and specific regulator of plasmacytoid dendritic cell development. Cell 135:37–48.

Crow, M.K., and L. Ronnblom. 2019. Type I interferons in host defence and inflammatory diseases. Lupus Sci Med 6:e000336.

Decalf, J., S. Fernandes, R. Longman, M. Ahloulay, F. Audat, F. Lefrerre, C.M. Rice, S. Pol, and M.L. Albert. 2007. Plasmacytoid dendritic cells initiate a complex chemokine and cytokine network and are a viable drug target in chronic HCV patients. J Exp Med 204:2423–2437.

Ghosh, H.S., B. Cisse, A. Bunin, K.L. Lewis, and B. Reizis. 2010. Continuous expression of the transcription factor e2-2 maintains the cell fate of mature plasmacytoid dendritic cells. Immunity 33:905–916.

Gregersen, P.K., G. Klein, M. Keogh, M. Kern, M. DeFranco, K.R. Simpfendorfer, S.J. Kim, and B. Diamond. 2015. The Genotype and Phenotype (GaP) registry: a living biobank for the analysis of quantitative traits. Immunol Res 63:107–112.

Gundacker, N.C., V.J. Haudek, H. Wimmer, A. Slany, J. Griss, V. Bochkov, C. Zielinski, O. Wagner, J. Stockl, and C. Gerner. 2009. Cytoplasmic proteome and secretome profiles of differently stimulated human dendritic cells. J Proteome Res 8:2799–2811.

Hancharou, A.Y., K.A. Chyzh, L.P. Titov, N.F. Soroka, and L.M. DuBuske. 2012. Characterization Of Plasmacytoid Dendritic Cells From Blood Of Patients With Systemic Lupus Erythematosus. Journal of Allergy and Clinical Immunology 129:

Kang, H.M., M. Subramaniam, S. Targ, M. Nguyen, L. Maliskova, E. McCarthy, E. Wan, S. Wong, L. Byrnes, C.M. Lanata, R.E. Gate, S. Mostafavi, A. Marson, N. Zaitlen, L.A. Criswell, and C.J. Ye. 2017. Multiplexed droplet single-cell RNA-sequencing using natural genetic variation. Nature Biotechnology 36:89–94.

Korsunsky, I., N. Millard, J. Fan, K. Slowikowski, F. Zhang, K. Wei, Y. Baglaenko, M. Brenner, P.-r. Loh, and S. Raychaudhuri. 2019. Fast, sensitive and accurate integration of single-cell data with Harmony. Nature Methods 16:1289–1296.

Lennert, K., and W. Remmele. 1958. Karyometrische Untersuchungen an Lymphknotenzellen des Menschen. Acta Haematologica 19:99–113.

Lichtner, M., C.M. Mastroianni, R. Rossi, G. Russo, V. Belvisi, R. Marocco, C. Mascia, C. Del Borgo, F. Mengoni, I. Sauzullo, G. d’Ettorre, C. D’Agostino, A.P. Massetti, and V. Vullo. 2011. Severe and persistent depletion of circulating plasmacytoid dendritic cells in patients with 2009 pandemic H1N1 infection. PLoS One 6:e19872.

Matsui, T., J.E. Connolly, M. Michnevitz, D. Chaussabel, C.I. Yu, C. Glaser, S. Tindle, M. Pypaert, H. Freitas, B. Piqueras, J. Banchereau, and A.K. Palucka. 2009. CD2 distinguishes two subsets of human plasmacytoid dendritic cells with distinct phenotype and functions. J Immunol 182:6815–6823.

Onodi, F., L. Bonnet-Madin, L. Meertens, L. Karpf, J. Poirot, S.Y. Zhang, C. Picard, A. Puel, E. Jouanguy, Q. Zhang, J. Le Goff, J.M. Molina, C. Delaugerre, J.L. Casanova, A. Amara, and V. Soumelis. 2021. SARS-CoV-2 induces human plasmacytoid predendritic cell diversification via UNC93B and IRAK4. J Exp Med 218:

Peruzzi, B., S. Bencini, M. Capone, A. Mazzoni, L. Maggi, L. Salvati, A. Vanni, C. Orazzini, C. Nozzoli, A. Morettini, L. Poggesi, F. Pieralli, A. Peris, A. Bartoloni, A.M. Vannucchi, F. Liotta, R. Caporale, L. Cosmi, and F. Annunziato. 2020. Quantitative and qualitative alterations of circulating myeloid cells and plasmacytoid DC in SARS-CoV-2 infection. Immunology 161:345–353.

Polson, C., P. Sarkar, B. Incledon, V. Raguvaran, and R. Grant. 2003. Optimization of protein precipitation based upon effectiveness of protein removal and ionization effect in liquid chromatography–tandem mass spectrometry. Journal of Chromatography B 785:263–275.

Ren, X., W. Wen, X. Fan, W. Hou, B. Su, P. Cai, J. Li, Y. Liu, F. Tang, F. Zhang, Y. Yang, J. He, W. Ma, J. He, P. Wang, Q. Cao, F. Chen, Y. Chen, X. Cheng, G. Deng, X. Deng, W. Ding, Y. Feng, R. Gan, C. Guo, W. Guo, S. He, C. Jiang, J. Liang, Y.M. Li, J. Lin, Y. Ling, H. Liu, J. Liu, N. Liu, S.Q. Liu, M. Luo, Q. Ma, Q. Song, W. Sun, G. Wang, F. Wang, Y. Wang, X. Wen, Q. Wu, G. Xu, X. Xie, X. Xiong, X. Xing, H. Xu, C. Yin, D. Yu, K. Yu, J. Yuan, B. Zhang, P. Zhang, T. Zhang, J. Zhao, P. Zhao, J. Zhou, W. Zhou, S. Zhong, X. Zhong, S. Zhang, L. Zhu, P. Zhu, B. Zou, J. Zou, Z. Zuo, F. Bai, X. Huang, P. Zhou, Q. Jiang, Z. Huang, J.X. Bei, L. Wei, X.W. Bian, X. Liu, T. Cheng, X. Li, P. Zhao, F.S. Wang, H. Wang, B. Su, Z. Zhang, K. Qu, X. Wang, J. Chen, R. Jin, and Z. Zhang. 2021. COVID-19 immune features revealed by a large-scale single-cell transcriptome atlas. Cell 184:1895–1913 e1819.

Shimura, C., T. Satoh, K. Igawa, K. Aritake, Y. Urade, M. Nakamura, and H. Yokozeki. 2010. Dendritic cells express hematopoietic prostaglandin D synthase and function as a source of prostaglandin D2 in the skin. Am J Pathol 176:227–237.

Siegal, F.P., N. Kadowaki, M. Shodell, P.A. Fitzgerald-Bocarsly, K. Shah, S. Ho, S. Antonenko, and Y.-J. Liu. 1999. The nature of the principal type 1 interferon-producing cells in human blood. Science 284:1835–1837.

Stuart, T., A. Butler, P. Hoffman, C. Hafemeister, E. Papalexi, W.M. Mauck, Y. Hao, M. Stoeckius, P. Smibert, and R. Satija. 2019. Comprehensive Integration of Single-Cell Data. Cell 177:1888–1902.e1821.

Thomas, J.M., Z. Pos, J. Reinboth, R.Y. Wang, E. Wang, G.M. Frank, P. Lusso, G. Trinchieri, H.J. Alter, F.M. Marincola, and E. Thomas. 2014. Differential responses of plasmacytoid dendritic cells to influenza virus and distinct viral pathogens. J Virol 88:10758–10766.

Thul, P.J., L. Akesson, M. Wiking, D. Mahdessian, A. Geladaki, H. Ait Blal, T. Alm, A. Asplund, L. Bjork, L.M. Breckels, A. Backstrom, F. Danielsson, L. Fagerberg, J. Fall, L. Gatto, C. Gnann, S. Hober, M. Hjelmare, F. Johansson, S. Lee, C. Lindskog, J. Mulder, C.M. Mulvey, P. Nilsson, P. Oksvold, J. Rockberg, R. Schutten, J.M. Schwenk, A. Sivertsson, E. Sjostedt, M. Skogs, C. Stadler, D.P. Sullivan, H. Tegel, C. Winsnes, C. Zhang, M. Zwahlen, A. Mardinoglu, F. Ponten, K. von Feilitzen, K.S. Lilley, M. Uhlen, and E. Lundberg. 2017. A subcellular map of the human proteome. Science 356:

Villani, A.C., R. Satija, G. Reynolds, S. Sarkizova, K. Shekhar, J. Fletcher, M. Griesbeck, A. Butler, S. Zheng, S. Lazo, L. Jardine, D. Dixon, E. Stephenson, E. Nilsson, I. Grundberg, D. McDonald, A. Filby, W. Li, P.L. De Jager, O. Rozenblatt-Rosen, A.A. Lane, M. Haniffa, A. Regev, and N. Hacohen. 2017. Single-cell RNA-seq reveals new types of human blood dendritic cells, monocytes, and progenitors. Science 356:

Wilk, A.J., A. Rustagi, N.Q. Zhao, J. Roque, G.J. Martinez-Colon, J.L. McKechnie, G.T. Ivison, T. Ranganath, R. Vergara, T. Hollis, L.J. Simpson, P. Grant, A. Subramanian, A.J. Rogers, and C.A. Blish. 2020. A single-cell atlas of the peripheral immune response in patients with severe COVID-19. Nat Med 26:1070–1076.

Wobma, H.M., M.A. Tamargo, S. Goeta, L.M. Brown, R. Duran-Struuck, and G. Vunjak-Novakovic. 2018. The influence of hypoxia and IFN-gamma on the proteome and metabolome of therapeutic mesenchymal stem cells. Biomaterials 167:226–234.

Yun, T.J., S. Igarashi, H. Zhao, O.A. Perez, M.R. Pereira, E. Zorn, Y. Shen, F. Goodrum, A. Rahman, P.A. Sims, D.L. Farber, and B. Reizis. 2021. Human plasmacytoid dendritic cells mount a distinct antiviral response to virus-infected cells. Sci Immunol 6:

Zhang, H., J.D. Gregorio, T. Iwahori, X. Zhang, O. Choi, L.L. Tolentino, T. Prestwood, Y. Carmi, and E.G. Engleman. 2017. A distinct subset of plasmacytoid dendritic cells induces activation and differentiation of B and T lymphocytes. Proc Natl Acad Sci U S A 114:1988–1993.

