## Supplementary Materials for "Proteomic and single-cell transcriptomic dissection of human plasmacytoid dendritic cell response to influenza virus"

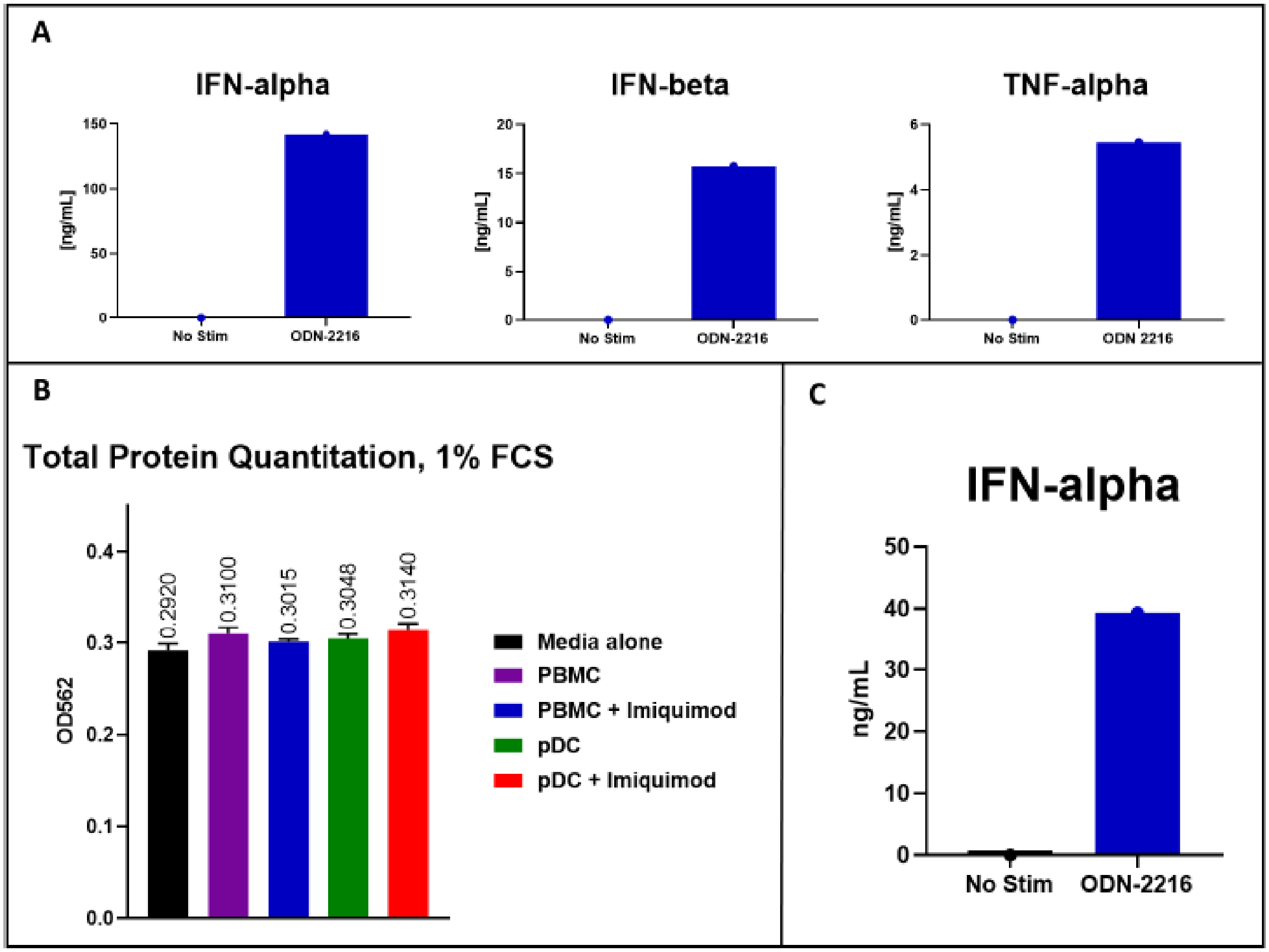

<u>Figure S1: Media optimization towards LC-MS/MS.</u>

A) pDCs cultured in the presence of 1% FCS are highly responsive to stimulation, producing robust amounts of IFN-alpha, IFN-beta and TNF-alpha. B) BCA assay for total protein content. Despite substantial cytokine production, pDCs only marginally enrich the protein content of the culture media, as >90% of media protein is derived from FCS alone. Stimulated pDCs were no different in this regard than unstimulated pDCs or PBMCs. C) pDCs cultured in Advanced RPMI, containing minimal amounts of albumin, insulin and holo-transferrin, without any serum supplement, remain highly functional. ODN2216, TLR9 agonist. Imiquimod, TLR7 agonist.

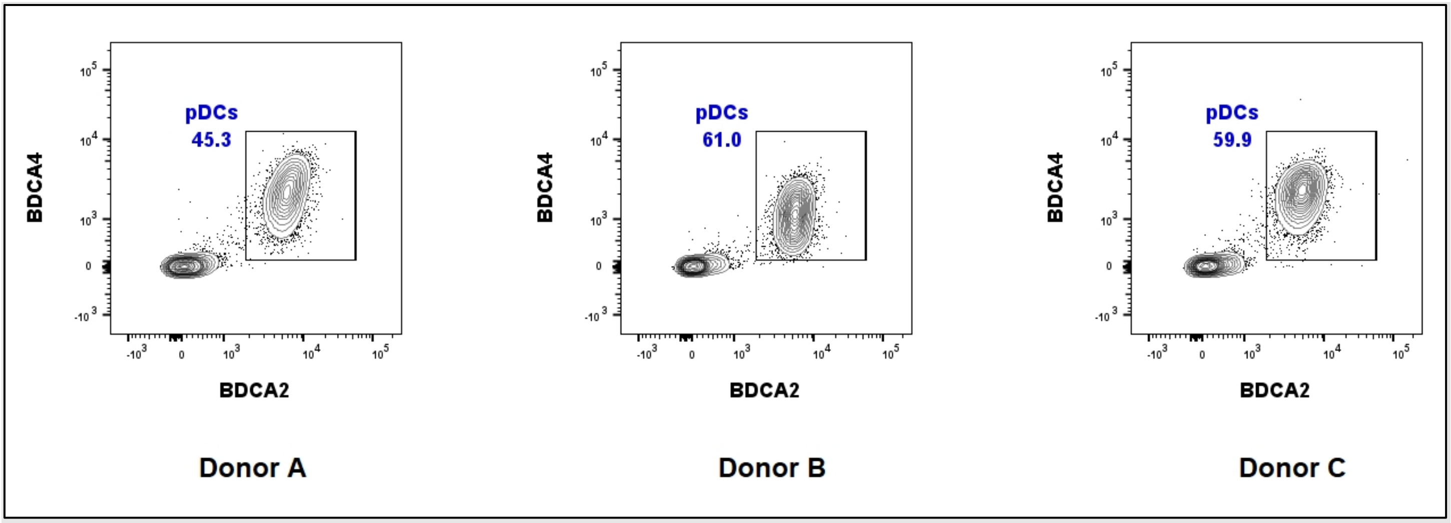

Figure S2: pDC purity for the secretome cultures.

pDCs were enriched from 3 donors emphasizing yield over purity. Purity was determined by staining for BDCA2 and BDCA4. Cell yields were: Donor A, 4.2e6; Donor B 3.7e6, Donor C 1.5e6 total cells. Cell viability was >95% by fluorescent cell counting using Acridine Orange / Propidium Iodide staining and the Cellometer Auto 2000 instrument. Pre-enrichment pDC frequency was 0.2-0.3% among PBMCs in all donors.

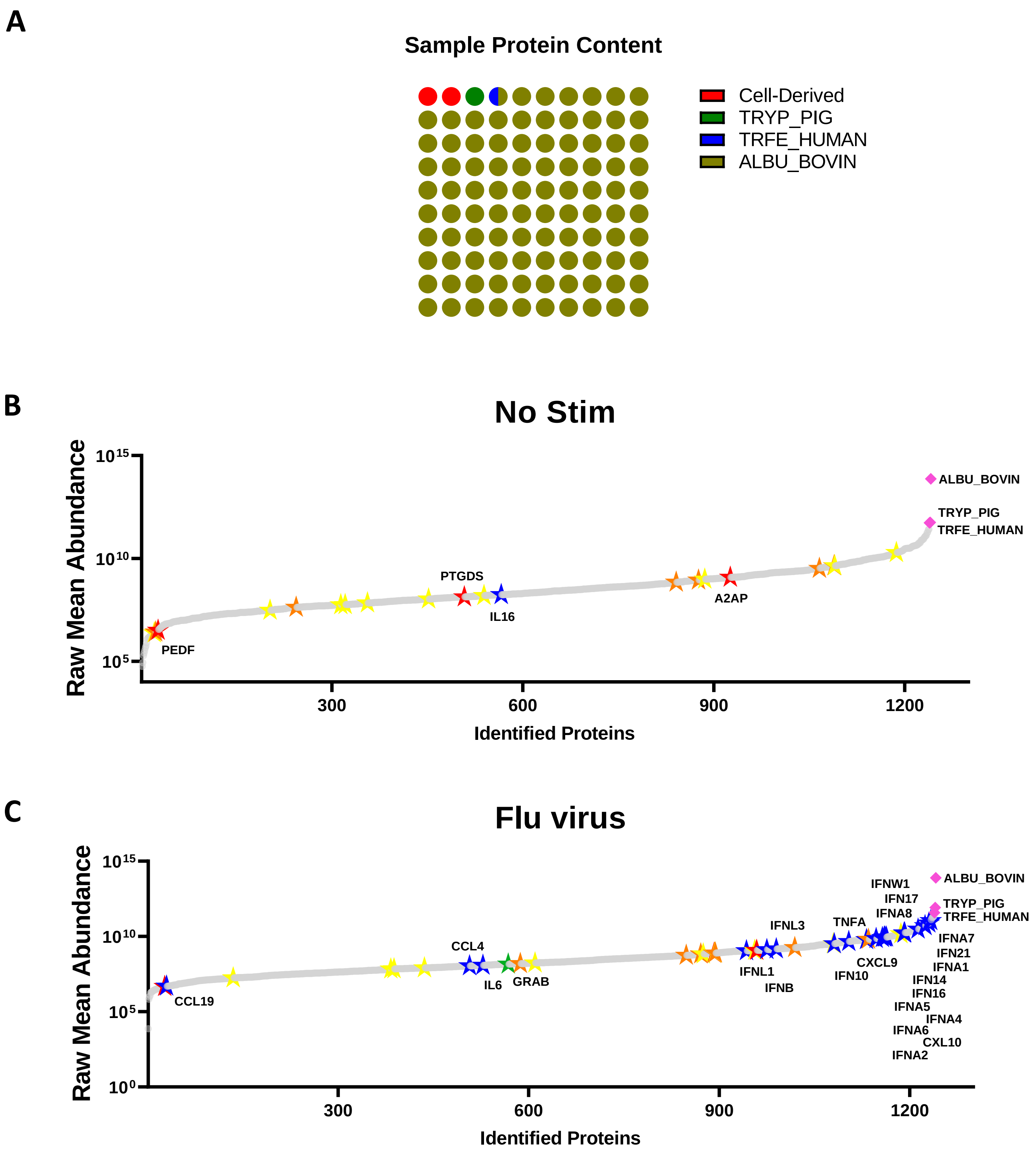
A
Sample Protein Content
Cell-Derived
TRYP_PIG
TRFE_HUMAN
ALBU_BOVIN
B
No Stim
Raw Mean Abundance
Identified Proteins
ALBU_BOVIN
TRYP_PIG
TRFE_HUMAN
PTGDS
IL16
A2AP
PEDF
C
Flu virus
Raw Mean Abundance
Identified Proteins
ALBU_BOVIN
TRYP_PIG
TRFE_HUMAN
IFNW1
IFN17
IFNA8
TNFA
IFNL3
CCL4
IL6
GRAB
CCL19
IFNL1
IFNB
CXCL9
IFN10
IFNA7
IFN21
IFNA1
IFN14
IFN16
IFNA5
IFNA4
IFNA6
CXL10
IFNA2

Figure S3. The pDC baseline and influenza-induced secretomes.

A) Protein content in the samples by relative abundance. B) Proteins identified in the unstimulated group ranked by average abundance. C) Proteins identified in the Influenza H1N1-stimulated group ranked by average abundance. For B and C: Data points represent unique proteins. Pink diamonds; exogenous proteins. Yellow, orange, red stars: proteins annotated as 'secreted' in the UniProt database, with relative enrichment of expression in pDCs among circulating leukocytes according to the Protein Atlas (http://www.proteinatlas.org/), with yellow=mild, orange=moderate, and red=significant pDC enrichment. Blue stars; cytokines/chemokines. Green star; granzyme B. Gray data points, with all points fused into a single line, represent proteins not annotated as 'secreted' in UniProt or not significantly enriched for expression in pDCs according to the Protein Atlas. Select proteins of interest are directly labeled.

## A

### HEMO_HUMAN, Hemopexin

Example Peptide Abundances

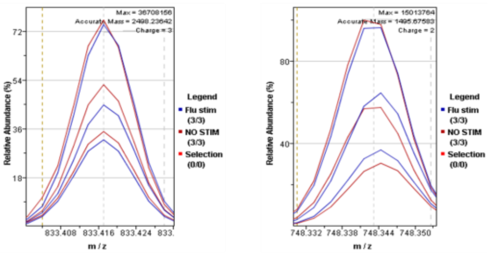

EVGTPHGIILDSVDAA FIC[160.0307]PGSSR (3+)

YYC[160.0307]FQGNQFLR (2+)

| Peptide Count | Mean Ion Score | Mean Protein Score | Ratio (Flu stim/ NO STIM) | P-value (Flu stim vs. NO STIM) |
|---|---|---|---|---|
| 20 | 57 | 1,133 | 0.99 | 0.89 |

### HEMO_HUMAN, Hemopexin

Example MS/MS Spectra

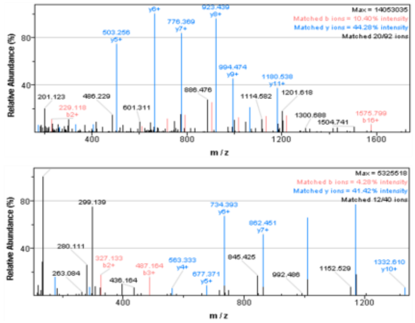

EVGTPHGIILDSVDAA FIC[160.0307]PGSSR (3+)

YYC[160.0307]F QGNQFLR (2+)

## B

### DPYL2_HUMAN, Dihydropyrimidinase-related protein 2

Example Peptide Abundances

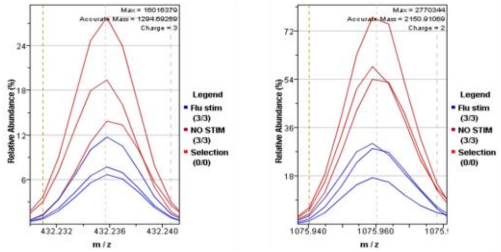

MVIPGGIDVHTR (3+)

FQMPDQGMTSADDFFQGTK (2+)

| Peptide Count | Mean Ion Score | Mean Protein Score | Ratio (Flu stim/ NO STIM) | P-value (Flu stim vs. NO STIM) |
|---|---|---|---|---|
| 21 | 61 | 1,065 | 0.46 | 3.4E-27 |

### DPYL2_HUMAN, Dihydropyrimidinase-related protein 2

Example MS/MS Spectra

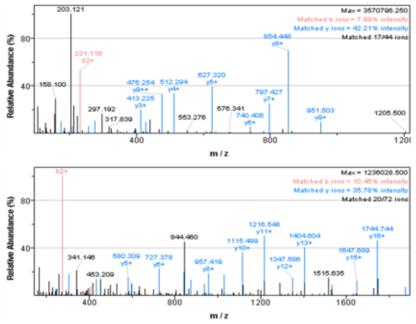

MVIPGGIDVHTR (3+)

FQMPDQGMTSA DDFFQGTK (2+)

## C

### TNFA_HUMAN, Tumor necrosis factor

Example Peptide Abundances

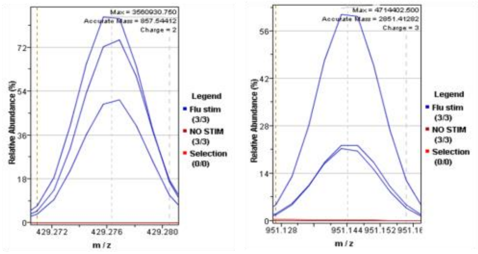

VNLLSAIK (2+)

ETPEGAEAKPWYEPIYL GGVFQLEK (3+)

| Peptide Count | Mean Ion Score | Mean Protein Score | Ratio (Flu stim/ NO STIM) | P-value (Flu stim vs. NO STIM) |
|---|---|---|---|---|
| 5 | 38 | 137 | 72.1 | 3.5E-17 |

### TNFA_HUMAN, Tumor necrosis factor

Example MS/MS Spectra

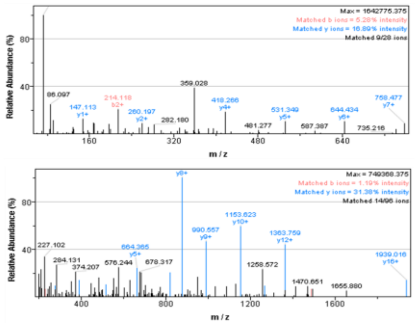

VNLLSAIK (2+)

ETPEGAEAKPWYEPIYL GGVFQLEK (3+)

## D

### IFN21_HUMAN, Interferon alpha-21

Example Peptide Abundances

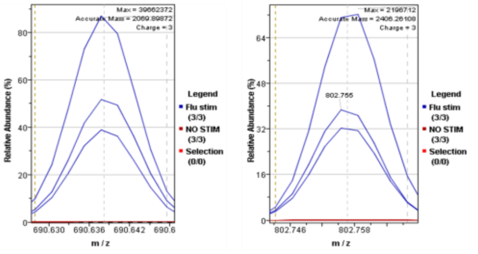

HDFGFPQEEFDG NQFQK (3+)

AQAISVLHEMIQ QTFNLFSTK (3+)

| Peptide Count | Mean Ion Score | Mean Protein Score | Ratio (Flu stim/ NO STIM) | P-value (Flu stim vs. NO STIM) |
|---|---|---|---|---|
| 7 | 64 | 1,432 | 468 | 2.0E-21 |

### IFN21_HUMAN, Interferon alpha-21

Example MS/MS Spectra

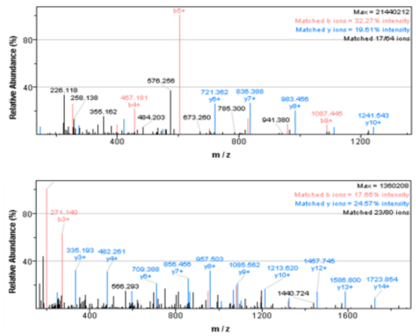

HDFGFPQEEFDG NQFQK (3+)

AQAISVLHEMIQ QTFNLFSTK (3+)

Figure S4. Representative protein species identified in the baseline and induced pDC secretomes.

For each protein, example graphics for two peptides are shown, including a monoisotopic MS peak (feature plotted as relative % abundance) as generated by the Elucidator program (charge state indicated), and an ms/ms spectrum for each peptide. Blue peaks are y-ions, red peaks are b-ions, and black are unmatched peaks.The feature peaks (left) in each case are an overlay of aligned monoisotopic peaks of MS spectra for the 12 LC-MS/MS runs in the experiment. Below the MS feature graphics is a table representing quantitation and identification details for each protein. A. Hemopexin, a protein produced by the liver and not known to be produced by circulating leukocytes, is not significantly different in preparations from unstimulated and flu stimulated cells. B. DPYL2, a metabolic enzyme, is enriched in the media of unstimulated cells compared to flu stimulated cells. C-D. Interferon α-1/13 and TNF-α are tremendously upregulated in the flu stimulated condition.

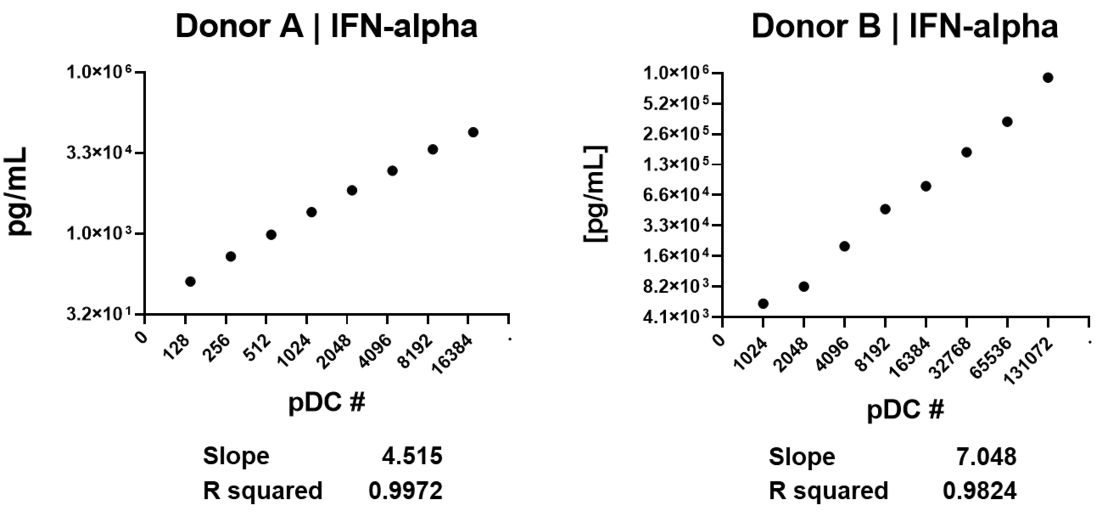

<u>Figure S5. Relationship between pDC input in IFN-α output using influenza H1N1 as the stimulus.</u>

Data derived from pDCs cultured for 24hrs in 96-well plates, with cells in a fixed 200uL culture volume per well. IFN-alpha levels determined using pan-subtype ELISA.

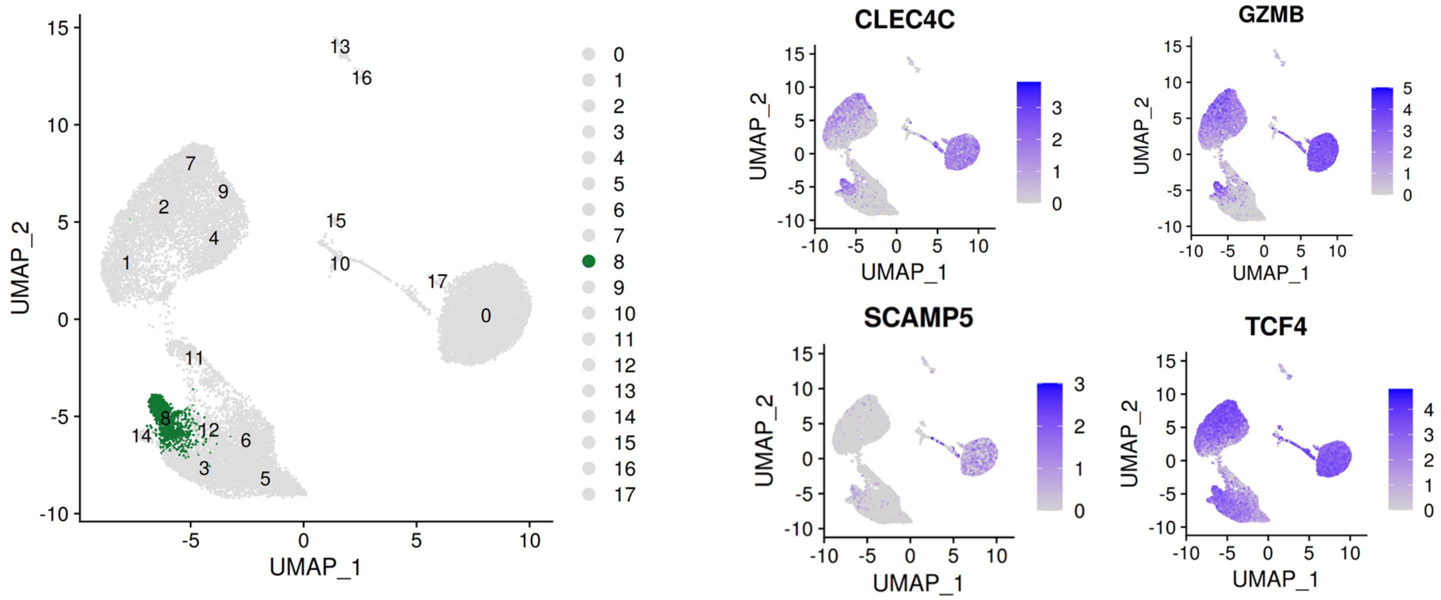

Figure S6. Cluster 8 retains features of unperturbed pDCs to 24hrs in the presence of influenza virus.

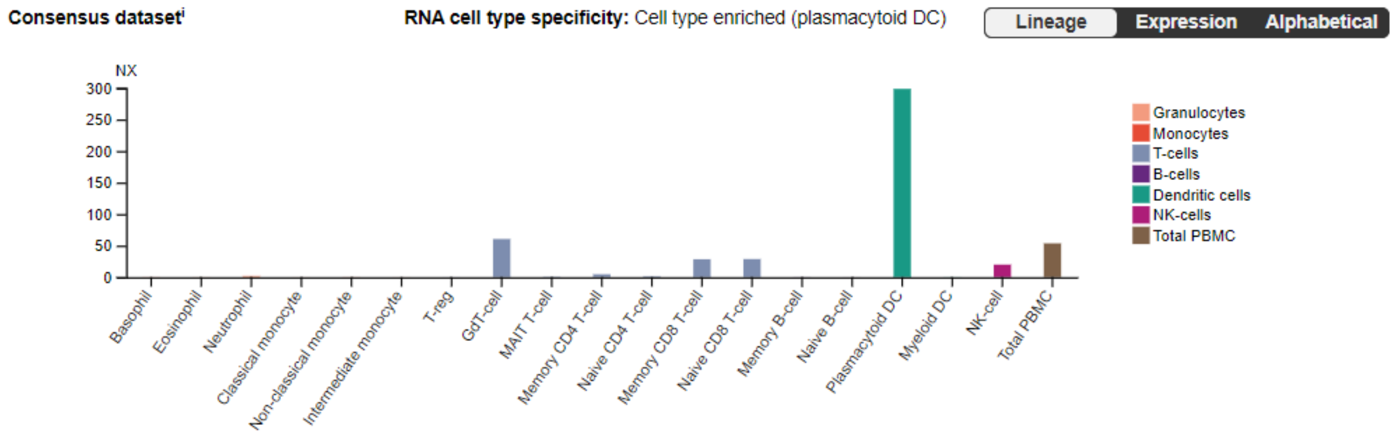

<u>Figure S7. Expression of Granzyme B across leukocyte subsets in the Human Protein Atlas.</u>

Table S1 LC-MS/MS Identified Proteins in the pDC Secretome with normalized expression data.

| UniProt ID | Peptide Count | Ratio (Flu stim:NO STIM) | P-value (Flu stim vs. NO STIM) | -LOG(P) |
|---|---|---|---|---|
| IFN10 | 2 | 306 | 0 | >40 |
| IFN17 | 2 | 91.6 | 0 | >40 |
| IFNW1 | 3 | 44.2 | 0 | >40 |
| IFNA8 | 4 | 901 | 0 | >40 |
| IFNA7 | 10 | 470 | 0 | >40 |
| IFNB | 3 | 647 | 3.6E-40 | 39.4432 |
| IFN14 | 4 | 508 | 1.0E-33 | 33.0001 |
| IFNA4 | 4 | 550 | 2.3E-30 | 29.6356 |
| IFNA6 | 5 | 234 | 1.1E-28 | 27.9531 |
| IFN16 | 6 | 94.9 | 3.1E-28 | 27.5147 |
| DPYL2 | 21 | 0.46 | 3.4E-27 | 26.472 |
| IFNA5 | 2 | 56.1 | 1.3E-25 | 24.8755 |
| PNCB | 4 | 0.36 | 9.5E-25 | 24.0244 |
| IFNL3 | 2 | 186 | 2.0E-23 | 22.71 |
| IFN21 | 7 | 468 | 2.0E-21 | 20.6942 |
| IFNA2 | 4 | 296 | 2.2E-18 | 17.6657 |
| TNFA | 5 | 72.1 | 3.5E-17 | 16.4548 |
| CALU | 1 | 8.21 | 1.1E-16 | 15.9626 |
| IFNA1 | 10 | 499 | 4.6E-16 | 15.3368 |
| H2AY | 12 | 0.44 | 2.6E-13 | 12.5894 |
| CCL4 | 1 | 6.85 | 6.5E-13 | 12.1896 |
| COR1B | 5 | 0.43 | 3.4E-12 | 11.4641 |
| H1X | 5 | 0.28 | 1.4E-10 | 9.86614 |
| LGMN | 4 | 11.0 | 1.8E-10 | 9.7428 |
| PDCD4 | 2 | 0.14 | 5.3E-10 | 9.27597 |
| TCL1A | 2 | 0.49 | 8.8E-10 | 9.05453 |
| GLO2 | 2 | 0.35 | 4.4E-09 | 8.36002 |
| IFIT3 | 7 | 40.5 | 8.1E-09 | 8.09334 |
| HP1B3 | 2 | 0.24 | 1.1E-08 | 7.96098 |
| RHG17 | 6 | 0.48 | 5.2E-08 | 7.28802 |
| COR1C | 17 | 0.39 | 9.2E-08 | 7.03763 |
| MPRD | 2 | 0.41 | 2.9E-07 | 6.53745 |
| CXL10 | 6 | 118 | 4.7E-07 | 6.32873 |
| PACN1 | 12 | 0.41 | 4.8E-07 | 6.31623 |
| FSCN1 | 12 | 11.0 | 6.1E-07 | 6.21789 |
| PRP19 | 6 | 0.41 | 7.6E-07 | 6.11827 |

| | | | | |
|---|---|---|---|---|
| **PAF1** | **1** | **0.26** | **8.5E-07** | **6.07109** |
| **VAPA** | **2** | **0.35** | **8.7E-07** | **6.06053** |
| **VP13C** | **6** | **0.38** | **1.1E-06** | **5.95468** |
| **FLNB** | **56** | **0.44** | **1.5E-06** | **5.81446** |
| **VATB2** | **6** | **0.51** | **2.6E-06** | **5.58603** |
| **SC22B** | **3** | **0.23** | **2.6E-06** | **5.57873** |
| **CNDP2** | **21** | **0.46** | **2.8E-06** | **5.55705** |
| **RAB7A** | **5** | **0.36** | **3.5E-06** | **5.45817** |
| **CATB** | **5** | **16.1** | **4.6E-06** | **5.34008** |
| **LMNB2** | **15** | **0.36** | **5.0E-06** | **5.29722** |
| **RB11B** | **8** | **0.33** | **6.0E-06** | **5.21961** |
| **CATC** | **14** | **33.0** | **6.3E-06** | **5.19921** |
| **CLCA** | **1** | **0.40** | **6.7E-06** | **5.17594** |
| **H15** | **8** | **0.46** | **9.7E-06** | **5.01341** |
| **SC24C** | **9** | **0.42** | **1.1E-05** | **4.94348** |
| **RUXE** | **2** | **0.28** | **1.3E-05** | **4.87484** |
| **VATG1** | **3** | **0.53** | **1.4E-05** | **4.84285** |
| **U520** | **6** | **0.36** | **1.5E-05** | **4.83239** |
| **ROCK1** | **1** | **0.49** | **1.5E-05** | **4.81531** |
| **RU17** | **6** | **0.40** | **1.7E-05** | **4.76246** |
| **ISG20** | **2** | **4.84** | **1.7E-05** | **4.7602** |
| **PAXX** | **4** | **0.38** | **1.9E-05** | **4.71897** |
| **H14** | **8** | **0.40** | **2.1E-05** | **4.66979** |
| **RET4** | **5** | **0.73** | **2.4E-05** | **4.62599** |
| **HNRL2** | **7** | **0.45** | **3.5E-05** | **4.45556** |
| **IFNL1** | **3** | **31.7** | **3.5E-05** | **4.45432** |
| **ARHG7** | **3** | **0.32** | **3.9E-05** | **4.40506** |
| **FA49B** | **14** | **0.45** | **4.3E-05** | **4.36917** |
| **IC1** | **4** | **18.3** | **4.7E-05** | **4.33106** |
| **CAB45** | **3** | **6.61** | **5.2E-05** | **4.28108** |
| **SRRT** | **1** | **0.40** | **5.9E-05** | **4.22775** |
| **KTN1** | **3** | **0.34** | **6.8E-05** | **4.16577** |
| **CAPG** | **14** | **0.50** | **6.9E-05** | **4.15889** |
| **HDAC2** | **3** | **0.27** | **7.0E-05** | **4.15783** |
| **H12** | **1** | **0.43** | **8.1E-05** | **4.0913** |
| **ZC3HF** | **1** | **0.39** | **8.8E-05** | **4.05596** |
| **LGUL** | **1** | **0.33** | **9.8E-05** | **4.00997** |
| **TLN1** | **85** | **0.50** | **9.8E-05** | **4.00948** |
| **VPS25** | **1** | **0.22** | **1.0E-04** | **3.99396** |
| **HNRL1** | **1** | **0.36** | **1.0E-04** | **3.97964** |
| **NUMA1** | **15** | **0.34** | **1.1E-04** | **3.95195** |
| **HPLN3** | **5** | **43.5** | **1.1E-04** | **3.94539** |
| **STAT1** | **14** | **7.51** | **1.2E-04** | **3.9333** |
| **IFIT2** | **5** | **46.5** | **1.4E-04** | **3.84802** |

| | | | | |
|---|---|---|---|---|
| RBM25 | 2 | 0.38 | 1.4E-04 | 3.84436 |
| AN32A | 4 | 0.42 | 1.6E-04 | 3.80883 |
| ARP2 | 13 | 0.51 | 1.6E-04 | 3.79942 |
| RBP56 | 2 | 0.39 | 1.6E-04 | 3.79507 |
| AIMP1 | 3 | 0.24 | 1.7E-04 | 3.78068 |
| KT3K | 1 | 2.91 | 1.7E-04 | 3.76221 |
| AIFM1 | 5 | 0.40 | 1.7E-04 | 3.75945 |
| SRPRA | 1 | 0.15 | 1.7E-04 | 3.75721 |
| 11-Sep | 4 | 0.43 | 1.8E-04 | 3.744 |
| GNPI1 | 4 | 0.39 | 1.8E-04 | 3.73779 |
| MCTS1 | 1 | 0.43 | 1.9E-04 | 3.72862 |
| SEM7A | 8 | 5.49 | 1.9E-04 | 3.72538 |
| RSMB, RSMN | 3 | 0.45 | 2.1E-04 | 3.66776 |
| CAPZB | 15 | 0.49 | 2.2E-04 | 3.66314 |
| LYAM1 | 3 | 11.5 | 2.3E-04 | 3.64608 |
| AATC | 8 | 0.57 | 2.3E-04 | 3.63283 |
| WDR1 | 17 | 0.63 | 2.4E-04 | 3.61154 |
| IF4H | 1 | 8.60 | 2.5E-04 | 3.60206 |
| PNOC | 2 | 45.4 | 2.5E-04 | 3.59602 |
| ACTN4 | 46 | 0.50 | 2.7E-04 | 3.57675 |
| SRP14 | 3 | 0.42 | 2.7E-04 | 3.57528 |
| HCFC1 | 2 | 0.37 | 2.7E-04 | 3.56527 |
| RAB5C | 2 | 0.40 | 2.7E-04 | 3.56511 |
| CXCL9 | 8 | 147 | 2.8E-04 | 3.55705 |
| AP2M1 | 2 | 0.39 | 2.9E-04 | 3.5397 |
| KAD2 | 10 | 0.40 | 3.1E-04 | 3.50307 |
| SF01 | 4 | 0.42 | 3.3E-04 | 3.47912 |
| PTPRE | 8 | 0.37 | 3.3E-04 | 3.47599 |
| 7-Sep | 11 | 0.46 | 3.3E-04 | 3.47521 |
| GRAB | 2 | 18.1 | 3.4E-04 | 3.46916 |
| HNRPD | 9 | 0.45 | 3.4E-04 | 3.46332 |
| NH2L1 | 1 | 0.42 | 3.7E-04 | 3.43192 |
| PSB1 | 6 | 0.47 | 4.0E-04 | 3.3933 |
| SPS1 | 3 | 0.38 | 4.1E-04 | 3.39073 |
| PRDX2 | 8 | 0.56 | 4.1E-04 | 3.38977 |
| TBL1X | 1 | 0.25 | 4.1E-04 | 3.38595 |
| CAZA2 | 5 | 0.50 | 4.1E-04 | 3.38426 |
| GDIA | 5 | 0.48 | 4.3E-04 | 3.36957 |
| RAB2A | 3 | 0.30 | 4.4E-04 | 3.35793 |
| HMCS1 | 4 | 13.8 | 4.4E-04 | 3.35586 |
| LANC1 | 4 | 0.42 | 4.6E-04 | 3.33913 |
| HDHD2 | 2 | 0.41 | 4.8E-04 | 3.31912 |
| ESYT1 | 1 | 0.29 | 4.8E-04 | 3.31677 |
| DDAH2 | 5 | 0.37 | 5.0E-04 | 3.30417 |

| | | | | |
|---|---|---|---|---|
| COR1A | 20 | 0.56 | 5.1E-04 | 3.29183 |
| CATZ | 4 | 3.33 | 5.3E-04 | 3.27548 |
| DHX9 | 9 | 0.44 | 5.4E-04 | 3.26833 |
| 2-Sep | 8 | 0.41 | 5.6E-04 | 3.25524 |
| CAP1 | 24 | 0.56 | 5.6E-04 | 3.25065 |
| VINC | 35 | 0.51 | 5.6E-04 | 3.24841 |
| COPE | 5 | 0.49 | 5.8E-04 | 3.23897 |
| RBMX | 6 | 0.47 | 6.1E-04 | 3.21453 |
| PARK7 | 11 | 0.47 | 6.2E-04 | 3.20859 |
| RBP2 | 2 | 0.26 | 6.3E-04 | 3.19846 |
| PGK1 | 24 | 0.57 | 6.4E-04 | 3.19716 |
| ROAA | 4 | 0.48 | 6.5E-04 | 3.18595 |
| EIF3B | 7 | 0.47 | 6.9E-04 | 3.15939 |
| PSA1 | 11 | 0.52 | 7.0E-04 | 3.15633 |
| SMRC2 | 4 | 0.25 | 7.0E-04 | 3.15627 |
| SP16H | 4 | 0.40 | 7.0E-04 | 3.15596 |
| TRAD1 | 1 | 39.1 | 7.2E-04 | 3.14363 |
| COPA | 20 | 0.51 | 7.3E-04 | 3.13954 |
| CELF2 | 1 | 0.41 | 7.4E-04 | 3.12819 |
| SAHH | 10 | 0.50 | 7.6E-04 | 3.1197 |
| SF3B1 | 4 | 0.43 | 7.6E-04 | 3.11839 |
| ENPP2 | 5 | 12.9 | 7.7E-04 | 3.11283 |
| RL5 | 10 | 0.45 | 7.7E-04 | 3.11165 |
| UBC9 | 2 | 0.31 | 7.8E-04 | 3.10807 |
| APEX1 | 8 | 0.37 | 7.8E-04 | 3.10563 |
| IF4A2 | 4 | 0.43 | 8.0E-04 | 3.0968 |
| 6PGL | 8 | 0.45 | 8.3E-04 | 3.07956 |
| ILF3 | 10 | 0.46 | 8.4E-04 | 3.07603 |
| RMD3 | 1 | 0.26 | 8.5E-04 | 3.0689 |
| PRDX6 | 13 | 0.51 | 8.8E-04 | 3.05567 |
| AP1B1 | 9 | 0.51 | 9.0E-04 | 3.04793 |
| ABHEB | 3 | 0.43 | 9.1E-04 | 3.04206 |
| ARP3 | 20 | 0.52 | 9.5E-04 | 3.02452 |
| DDTL | 2 | 0.33 | 9.9E-04 | 3.00542 |
| TCRG1 | 1 | 0.36 | 1.0E-03 | 3 |
| PML | 1 | 0.35 | 1.0E-03 | 3 |
| RAB1A | 1 | 0.24 | 1.0E-03 | 3 |
| KYNU | 2 | 0.40 | 1.0E-03 | 3 |
| CUL4B | 3 | 0.46 | 1.0E-03 | 3 |
| RHOA | 4 | 0.51 | 1.0E-03 | 3 |
| PSB3 | 4 | 0.47 | 1.0E-03 | 3 |
| TSN | 4 | 0.45 | 1.0E-03 | 3 |
| PSA3 | 6 | 0.53 | 1.0E-03 | 3 |
| MK01 | 6 | 0.52 | 1.0E-03 | 3 |

| | | | | |
|---|---|---|---|---|
| PSA5 | 6 | 0.52 | 1.0E-03 | 3 |
| UBE2N | 6 | 0.47 | 1.0E-03 | 3 |
| SC23A | 7 | 0.46 | 1.0E-03 | 3 |
| SFPQ | 9 | 0.65 | 1.0E-03 | 3 |
| SNAA | 9 | 0.53 | 1.0E-03 | 3 |
| PSB4 | 9 | 0.49 | 1.0E-03 | 3 |
| MDHC | 11 | 0.53 | 1.0E-03 | 3 |
| SLAF7 | 1 | 9.51 | 2.0E-03 | 2.69897 |
| RUXGL | 1 | 0.45 | 2.0E-03 | 2.69897 |
| LSM3 | 1 | 0.44 | 2.0E-03 | 2.69897 |
| DYL1 | 1 | 0.37 | 2.0E-03 | 2.69897 |
| STRN3 | 1 | 0.25 | 2.0E-03 | 2.69897 |
| LHPP | 1 | 0.00 | 2.0E-03 | 2.69897 |
| AIP | 2 | 0.52 | 2.0E-03 | 2.69897 |
| LZIC | 2 | 0.40 | 2.0E-03 | 2.69897 |
| XRN2 | 2 | 0.34 | 2.0E-03 | 2.69897 |
| IL6 | 3 | 9.85 | 2.0E-03 | 2.69897 |
| UCHL3 | 3 | 0.52 | 2.0E-03 | 2.69897 |
| H2A1D | 3 | 0.51 | 2.0E-03 | 2.69897 |
| PSB10 | 3 | 0.50 | 2.0E-03 | 2.69897 |
| NIF3L | 3 | 0.43 | 2.0E-03 | 2.69897 |
| SRP09 | 4 | 0.41 | 2.0E-03 | 2.69897 |
| ISG15 | 5 | 24.5 | 2.0E-03 | 2.69897 |
| MA1A1 | 5 | 4.49 | 2.0E-03 | 2.69897 |
| FABP5 | 5 | 0.53 | 2.0E-03 | 2.69897 |
| IAH1 | 5 | 0.37 | 2.0E-03 | 2.69897 |
| CYC | 5 | 0.36 | 2.0E-03 | 2.69897 |
| PEBP1 | 5 | 0.32 | 2.0E-03 | 2.69897 |
| PDC6I | 6 | 0.57 | 2.0E-03 | 2.69897 |
| CAN1 | 6 | 0.50 | 2.0E-03 | 2.69897 |
| PNPH | 6 | 0.47 | 2.0E-03 | 2.69897 |
| HNRPL | 7 | 0.54 | 2.0E-03 | 2.69897 |
| DHX15 | 7 | 0.45 | 2.0E-03 | 2.69897 |
| NPM | 8 | 0.46 | 2.0E-03 | 2.69897 |
| ARPC2 | 9 | 0.54 | 2.0E-03 | 2.69897 |
| ILF2 | 9 | 0.48 | 2.0E-03 | 2.69897 |
| SF3B3 | 10 | 0.54 | 2.0E-03 | 2.69897 |
| NAGK | 12 | 0.57 | 2.0E-03 | 2.69897 |
| PSA6 | 12 | 0.54 | 2.0E-03 | 2.69897 |
| CAZA1 | 12 | 0.52 | 2.0E-03 | 2.69897 |
| ADA | 15 | 0.41 | 2.0E-03 | 2.69897 |
| CATA | 15 | 0.41 | 2.0E-03 | 2.69897 |
| GELS | 23 | 0.78 | 2.0E-03 | 2.69897 |
| GDIB | 27 | 0.52 | 2.0E-03 | 2.69897 |

| | | | | |
|---|---|---|---|---|
| IQGA1 | 28 | 0.51 | 2.0E-03 | 2.69897 |
| VIME | 34 | 0.66 | 2.0E-03 | 2.69897 |
| PP6R1 | 1 | 0.39 | 3.0E-03 | 2.52288 |
| HS105 | 2 | 3.10 | 3.0E-03 | 2.52288 |
| DNJA1 | 2 | 2.75 | 3.0E-03 | 2.52288 |
| PLIN3 | 2 | 0.55 | 3.0E-03 | 2.52288 |
| RBM8A | 2 | 0.50 | 3.0E-03 | 2.52288 |
| NUDT5 | 2 | 0.49 | 3.0E-03 | 2.52288 |
| LSM8 | 2 | 0.45 | 3.0E-03 | 2.52288 |
| SF3A1 | 2 | 0.42 | 3.0E-03 | 2.52288 |
| UBC12 | 2 | 0.41 | 3.0E-03 | 2.52288 |
| AP1S2 | 2 | 0.40 | 3.0E-03 | 2.52288 |
| PP14B | 2 | 0.36 | 3.0E-03 | 2.52288 |
| NLTP | 2 | 0.33 | 3.0E-03 | 2.52288 |
| HSP13 | 3 | 4.27 | 3.0E-03 | 2.52288 |
| AP2B1 | 3 | 0.44 | 3.0E-03 | 2.52288 |
| DDX1 | 3 | 0.43 | 3.0E-03 | 2.52288 |
| RS21 | 3 | 0.38 | 3.0E-03 | 2.52288 |
| RL22 | 3 | 0.34 | 3.0E-03 | 2.52288 |
| PSB9 | 4 | 0.53 | 3.0E-03 | 2.52288 |
| CBX3 | 4 | 0.45 | 3.0E-03 | 2.52288 |
| STX7 | 4 | 0.26 | 3.0E-03 | 2.52288 |
| A16A1 | 5 | 0.47 | 3.0E-03 | 2.52288 |
| F10A1 | 5 | 0.46 | 3.0E-03 | 2.52288 |
| LAP2A | 6 | 0.52 | 3.0E-03 | 2.52288 |
| TPR | 6 | 0.37 | 3.0E-03 | 2.52288 |
| EF1G | 10 | 0.62 | 3.0E-03 | 2.52288 |
| COF1 | 10 | 0.60 | 3.0E-03 | 2.52288 |
| DCPS | 10 | 0.42 | 3.0E-03 | 2.52288 |
| PSA7 | 11 | 0.52 | 3.0E-03 | 2.52288 |
| EIFCL | 11 | 0.51 | 3.0E-03 | 2.52288 |
| TWF2 | 11 | 0.45 | 3.0E-03 | 2.52288 |
| COPG1 | 12 | 0.55 | 3.0E-03 | 2.52288 |
| STIP1 | 13 | 0.71 | 3.0E-03 | 2.52288 |
| 1433Z | 14 | 0.55 | 3.0E-03 | 2.52288 |
| TCPA | 15 | 0.57 | 3.0E-03 | 2.52288 |
| SQSTM | 1 | 46.0 | 4.0E-03 | 2.39794 |
| ASC | 1 | 0.49 | 4.0E-03 | 2.39794 |
| THOC4 | 1 | 0.45 | 4.0E-03 | 2.39794 |
| H10 | 1 | 0.40 | 4.0E-03 | 2.39794 |
| CDC73 | 1 | 0.36 | 4.0E-03 | 2.39794 |
| LEO1 | 1 | 0.26 | 4.0E-03 | 2.39794 |
| DDRGK | 1 | 0.22 | 4.0E-03 | 2.39794 |
| TSNAX | 2 | 0.46 | 4.0E-03 | 2.39794 |

| | | | | |
|---|---|---|---|---|
| ERH | 2 | 0.11 | 4.0E-03 | 2.39794 |
| SYNC | 3 | 0.61 | 4.0E-03 | 2.39794 |
| PACN2 | 3 | 0.44 | 4.0E-03 | 2.39794 |
| 6-Sep | 4 | 0.53 | 4.0E-03 | 2.39794 |
| RO60 | 4 | 0.49 | 4.0E-03 | 2.39794 |
| SCFD1 | 4 | 0.45 | 4.0E-03 | 2.39794 |
| PA2G4 | 6 | 0.64 | 4.0E-03 | 2.39794 |
| PSB2 | 6 | 0.52 | 4.0E-03 | 2.39794 |
| ARPC3 | 7 | 0.59 | 4.0E-03 | 2.39794 |
| EF1D | 7 | 0.58 | 4.0E-03 | 2.39794 |
| RS12 | 7 | 0.55 | 4.0E-03 | 2.39794 |
| ARPC4 | 7 | 0.52 | 4.0E-03 | 2.39794 |
| 9-Sep | 8 | 0.35 | 4.0E-03 | 2.39794 |
| ESTD | 9 | 0.57 | 4.0E-03 | 2.39794 |
| PSB8 | 9 | 0.52 | 4.0E-03 | 2.39794 |
| LYSC | 9 | 0.48 | 4.0E-03 | 2.39794 |
| GLOD4 | 9 | 0.45 | 4.0E-03 | 2.39794 |
| H4 | 12 | 0.63 | 4.0E-03 | 2.39794 |
| EIF3A | 13 | 0.49 | 4.0E-03 | 2.39794 |
| TAGL2 | 13 | 0.48 | 4.0E-03 | 2.39794 |
| PSA4 | 14 | 0.49 | 4.0E-03 | 2.39794 |
| PLEC | 34 | 0.52 | 4.0E-03 | 2.39794 |
| VAS1 | 1 | 10.6 | 0.01 | 2.30103 |
| LIMD2 | 1 | 2.99 | 0.01 | 2.30103 |
| LCK | 1 | 0.40 | 0.01 | 2.30103 |
| DKC1 | 1 | 0.36 | 0.01 | 2.30103 |
| SRS11 | 1 | 0.12 | 0.01 | 2.30103 |
| DCNL1 | 2 | 0.37 | 5.0E-03 | 2.30103 |
| HNRDL | 3 | 0.64 | 5.0E-03 | 2.30103 |
| DEK | 3 | 0.52 | 5.0E-03 | 2.30103 |
| EIF3E | 3 | 0.50 | 5.0E-03 | 2.30103 |
| CPNS1 | 4 | 0.56 | 5.0E-03 | 2.30103 |
| VTDB | 6 | 1.28 | 5.0E-03 | 2.30103 |
| DX39B | 7 | 0.54 | 5.0E-03 | 2.30103 |
| RL12 | 7 | 0.48 | 5.0E-03 | 2.30103 |
| RAN | 8 | 0.54 | 5.0E-03 | 2.30103 |
| TCPH | 13 | 0.56 | 5.0E-03 | 2.30103 |
| TCPG | 14 | 0.58 | 5.0E-03 | 2.30103 |
| TCPD | 16 | 0.56 | 5.0E-03 | 2.30103 |
| TCPB | 17 | 0.58 | 5.0E-03 | 2.30103 |
| UBA1 | 24 | 0.58 | 5.0E-03 | 2.30103 |
| TIA1 | 1 | 0.61 | 0.01 | 2.22185 |
| CPSF6 | 1 | 0.51 | 0.01 | 2.22185 |
| PSMD5 | 1 | 0.45 | 0.01 | 2.22185 |

| | | | | |
|---|---|---|---|---|
| **ANM1** | **2** | **0.49** | **6.0E-03** | **2.22185** |
| **SMU1** | **2** | **0.46** | **6.0E-03** | **2.22185** |
| **PUF60** | **2** | **0.46** | **6.0E-03** | **2.22185** |
| **NGLY1** | **2** | **0.30** | **6.0E-03** | **2.22185** |
| **HCLS1** | **3** | **2.94** | **6.0E-03** | **2.22185** |
| **RPE** | **3** | **0.48** | **6.0E-03** | **2.22185** |
| **GSHB** | **4** | **0.54** | **6.0E-03** | **2.22185** |
| **HMGB2** | **6** | **0.52** | **6.0E-03** | **2.22185** |
| **ACPH** | **6** | **0.51** | **6.0E-03** | **2.22185** |
| **UB2V1** | **7** | **0.49** | **6.0E-03** | **2.22185** |
| **GSTP1** | **9** | **0.53** | **6.0E-03** | **2.22185** |
| **MVP** | **12** | **0.65** | **6.0E-03** | **2.22185** |
| **1433E** | **13** | **0.54** | **6.0E-03** | **2.22185** |
| **HLAE** | **1** | **29.6** | **0.01** | **2.1549** |
| **NTF2** | **1** | **0.43** | **0.01** | **2.1549** |
| **PFD5** | **1** | **0.36** | **0.01** | **2.1549** |
| **PP14A** | **1** | **0.32** | **0.01** | **2.1549** |
| **CSTF3** | **1** | **0.26** | **0.01** | **2.1549** |
| **EIF3I** | **5** | **0.53** | **7.0E-03** | **2.1549** |
| **SAE2** | **6** | **0.53** | **7.0E-03** | **2.1549** |
| **PSA2** | **7** | **0.45** | **7.0E-03** | **2.1549** |
| **RSSA** | **8** | **0.55** | **7.0E-03** | **2.1549** |
| **BUB3** | **8** | **0.47** | **7.0E-03** | **2.1549** |
| **COPD** | **8** | **0.47** | **7.0E-03** | **2.1549** |
| **HNRPC** | **12** | **0.58** | **7.0E-03** | **2.1549** |
| **PROF1** | **13** | **0.60** | **7.0E-03** | **2.1549** |
| **1433B** | **13** | **0.59** | **7.0E-03** | **2.1549** |
| **ENOA** | **20** | **0.54** | **7.0E-03** | **2.1549** |
| **DDX17** | **1** | **0.44** | **0.01** | **2.09691** |
| **WDR82** | **1** | **0.40** | **0.01** | **2.09691** |
| **SSRP1** | **2** | **0.45** | **8.0E-03** | **2.09691** |
| **OSGEP** | **2** | **0.43** | **8.0E-03** | **2.09691** |
| **CDC37** | **5** | **0.48** | **8.0E-03** | **2.09691** |
| **SYEP** | **6** | **0.39** | **8.0E-03** | **2.09691** |
| **AN32B** | **7** | **0.48** | **8.0E-03** | **2.09691** |
| **VATA** | **9** | **0.63** | **8.0E-03** | **2.09691** |
| **PSME1** | **9** | **0.57** | **8.0E-03** | **2.09691** |
| **PSA** | **12** | **0.53** | **8.0E-03** | **2.09691** |
| **2AAA** | **14** | **0.54** | **8.0E-03** | **2.09691** |
| **CLH1** | **37** | **0.61** | **8.0E-03** | **2.09691** |
| **IRF4** | **1** | **5.72** | **0.01** | **2.04576** |
| **B2MG** | **1** | **4.21** | **0.01** | **2.04576** |
| **STX12** | **1** | **0.45** | **0.01** | **2.04576** |
| **PLCB3** | **1** | **0.39** | **0.01** | **2.04576** |

| | | | | |
|---|---|---|---|---|
| DEOC | 1 | 0.36 | 0.01 | 2.04576 |
| REPI1 | 1 | 0.18 | 0.01 | 2.04576 |
| P66A | 1 | 0.18 | 0.01 | 2.04576 |
| VP26A | 2 | 0.60 | 9.0E-03 | 2.04576 |
| CPSF5 | 2 | 0.45 | 9.0E-03 | 2.04576 |
| UB2L3 | 6 | 0.61 | 9.0E-03 | 2.04576 |
| HDGF | 7 | 0.58 | 9.0E-03 | 2.04576 |
| TCPZ | 8 | 0.60 | 9.0E-03 | 2.04576 |
| PPIA | 14 | 0.60 | 9.0E-03 | 2.04576 |
| LMNB1 | 21 | 0.55 | 9.0E-03 | 2.04576 |
| GLYG | 1 | 0.48 | 0.01 | 2 |
| LSM5 | 1 | 0.44 | 0.01 | 2 |
| SART3 | 1 | 0.37 | 0.01 | 2 |
| SAP | 3 | 2.68 | 0.01 | 2 |
| UFM1 | 3 | 0.48 | 0.01 | 2 |
| XPP1 | 4 | 0.56 | 0.01 | 2 |
| EWS | 4 | 0.52 | 0.01 | 2 |
| EIF3D | 5 | 0.59 | 0.01 | 2 |
| FUBP1 | 6 | 0.58 | 0.01 | 2 |
| RBBP4 | 9 | 0.49 | 0.01 | 2 |
| PLSL | 45 | 0.57 | 0.01 | 2 |
| MYH9 | 78 | 0.63 | 0.01 | 2 |
| SMD1 | 1 | 0.36 | 0.01 | 1.95861 |
| NOLC1 | 1 | 0.29 | 0.01 | 1.95861 |
| SEC13 | 2 | 0.55 | 0.01 | 1.95861 |
| CZIB | 2 | 0.48 | 0.01 | 1.95861 |
| PFD2 | 2 | 0.46 | 0.01 | 1.95861 |
| LYPA1 | 3 | 0.53 | 0.01 | 1.95861 |
| API5 | 3 | 0.46 | 0.01 | 1.95861 |
| ARF1, ARF3 | 4 | 0.39 | 0.01 | 1.95861 |
| RS28 | 5 | 0.58 | 0.01 | 1.95861 |
| COPB | 7 | 0.52 | 0.01 | 1.95861 |
| TPIS | 15 | 0.61 | 0.01 | 1.95861 |
| IRF8 | 1 | 83.8 | 0.01 | 1.92082 |
| THMS2 | 1 | 14.2 | 0.01 | 1.92082 |
| COPZ1 | 1 | 0.51 | 0.01 | 1.92082 |
| BAP18 | 1 | 0.49 | 0.01 | 1.92082 |
| LRC59 | 1 | 0.45 | 0.01 | 1.92082 |
| BIN1 | 1 | 0.45 | 0.01 | 1.92082 |
| GLGB | 2 | 0.50 | 0.01 | 1.92082 |
| K1C17 | 2 | 0.21 | 0.01 | 1.92082 |
| GMFG | 3 | 0.57 | 0.01 | 1.92082 |
| PLPP | 3 | 0.30 | 0.01 | 1.92082 |
| DBNL | 9 | 0.69 | 0.01 | 1.92082 |

| | | | | |
|---|---|---|---|---|
| TPM4 | 11 | 0.60 | 0.01 | 1.92082 |
| IPO5 | 12 | 0.49 | 0.01 | 1.92082 |
| LDHB | 13 | 0.56 | 0.01 | 1.92082 |
| KCAB2 | 1 | 0.44 | 0.01 | 1.88606 |
| EIF3K | 2 | 0.51 | 0.01 | 1.88606 |
| IF1AX | 3 | 0.52 | 0.01 | 1.88606 |
| SPB8 | 4 | 0.49 | 0.01 | 1.88606 |
| HLAA | 5 | 2.71 | 0.01 | 1.88606 |
| GPX1 | 5 | 0.55 | 0.01 | 1.88606 |
| XPO1 | 7 | 0.59 | 0.01 | 1.88606 |
| KCD12 | 8 | 0.67 | 0.01 | 1.88606 |
| DDB1 | 9 | 0.50 | 0.01 | 1.88606 |
| ACTS | 15 | 0.77 | 0.01 | 1.88606 |
| TCPQ | 18 | 0.63 | 0.01 | 1.88606 |
| RN213 | 1 | 17.9 | 0.01 | 1.85387 |
| NC2A | 1 | 0.47 | 0.01 | 1.85387 |
| RFA3 | 1 | 0.44 | 0.01 | 1.85387 |
| GMPPB | 1 | 0.34 | 0.01 | 1.85387 |
| FCL | 2 | 0.56 | 0.01 | 1.85387 |
| THOP1 | 2 | 0.49 | 0.01 | 1.85387 |
| ABRAL | 3 | 0.50 | 0.01 | 1.85387 |
| SMD3 | 4 | 0.51 | 0.01 | 1.85387 |
| 1433F | 6 | 0.56 | 0.01 | 1.85387 |
| PARP1 | 8 | 0.41 | 0.01 | 1.85387 |
| IFIT1 | 1 | 8.42 | 0.02 | 1.82391 |
| MTNB | 1 | 0.35 | 0.02 | 1.82391 |
| ELAV1 | 3 | 0.49 | 0.02 | 1.82391 |
| GSHR | 4 | 0.59 | 0.02 | 1.82391 |
| FUBP2 | 6 | 0.63 | 0.02 | 1.82391 |
| 6PGD | 12 | 0.63 | 0.02 | 1.82391 |
| COPB2 | 14 | 0.57 | 0.02 | 1.82391 |
| TERA | 30 | 0.59 | 0.02 | 1.82391 |
| CMPK2 | 1 | 29.2 | 0.02 | 1.79588 |
| SWP70 | 1 | 2.76 | 0.02 | 1.79588 |
| APOA1 | 1 | 0.69 | 0.02 | 1.79588 |
| CNPY3 | 1 | 0.58 | 0.02 | 1.79588 |
| BROX | 1 | 0.47 | 0.02 | 1.79588 |
| LRC47 | 1 | 0.34 | 0.02 | 1.79588 |
| CATS | 2 | 6.80 | 0.02 | 1.79588 |
| GMPR2 | 2 | 0.54 | 0.02 | 1.79588 |
| IF4E | 2 | 0.53 | 0.02 | 1.79588 |
| MGN2 | 2 | 0.38 | 0.02 | 1.79588 |
| AP1M1 | 3 | 0.39 | 0.02 | 1.79588 |
| HNRPR | 5 | 0.68 | 0.02 | 1.79588 |

| | | | | |
|---|---|---|---|---|
| SET | 6 | 0.53 | 0.02 | 1.79588 |
| PGM2 | 8 | 0.50 | 0.02 | 1.79588 |
| HNRPU | 10 | 0.54 | 0.02 | 1.79588 |
| PRDX1 | 12 | 0.64 | 0.02 | 1.79588 |
| ARC1B | 15 | 0.67 | 0.02 | 1.79588 |
| TRFE | 65 | 0.75 | 0.02 | 1.79588 |
| DDX3X | 1 | 2.31 | 0.02 | 1.76955 |
| SFT2B | 1 | 0.43 | 0.02 | 1.76955 |
| PLRG1 | 1 | 0.37 | 0.02 | 1.76955 |
| SMCA4 | 1 | 0.29 | 0.02 | 1.76955 |
| CNN2 | 4 | 0.54 | 0.02 | 1.76955 |
| IF2G | 5 | 0.58 | 0.02 | 1.76955 |
| PTBP1 | 6 | 0.48 | 0.02 | 1.76955 |
| BUD31 | 1 | 0.50 | 0.02 | 1.74473 |
| LSM4 | 1 | 0.46 | 0.02 | 1.74473 |
| AP1G1 | 2 | 0.49 | 0.02 | 1.74473 |
| CHM1B | 2 | 0.45 | 0.02 | 1.74473 |
| LIS1 | 4 | 0.57 | 0.02 | 1.74473 |
| GDS1 | 5 | 0.52 | 0.02 | 1.74473 |
| RFA1 | 6 | 0.55 | 0.02 | 1.74473 |
| HSP7C | 28 | 0.63 | 0.02 | 1.74473 |
| PIPNB | 2 | 0.62 | 0.02 | 1.72125 |
| FKB15 | 3 | 0.58 | 0.02 | 1.72125 |
| IF4A3 | 3 | 0.55 | 0.02 | 1.72125 |
| 1433T | 5 | 0.59 | 0.02 | 1.72125 |
| SIAS | 5 | 0.46 | 0.02 | 1.72125 |
| MARE1 | 6 | 0.49 | 0.02 | 1.72125 |
| MYL6 | 7 | 0.66 | 0.02 | 1.72125 |
| H2B2F | 10 | 0.60 | 0.02 | 1.72125 |
| TMOD3 | 1 | 0.53 | 0.02 | 1.69897 |
| HEBP1 | 1 | 0.49 | 0.02 | 1.69897 |
| 2ABA | 2 | 0.61 | 0.02 | 1.69897 |
| SH3L3 | 3 | 0.55 | 0.02 | 1.69897 |
| EIF3H | 4 | 0.60 | 0.02 | 1.69897 |
| NDKB | 9 | 0.55 | 0.02 | 1.69897 |
| DPP3 | 14 | 0.59 | 0.02 | 1.69897 |
| PDC10 | 1 | 0.48 | 0.02 | 1.67778 |
| MTA2 | 2 | 0.39 | 0.02 | 1.67778 |
| SMC3 | 2 | 0.28 | 0.02 | 1.67778 |
| KINH | 5 | 0.66 | 0.02 | 1.67778 |
| MGN | 1 | 0.49 | 0.02 | 1.65758 |
| CSTN1 | 2 | 5.06 | 0.02 | 1.65758 |
| SF3B6 | 2 | 0.47 | 0.02 | 1.65758 |
| PPM1G | 4 | 0.46 | 0.02 | 1.65758 |

| | | | | |
|---|---|---|---|---|
| YTDC1 | 1 | 0.40 | 0.02 | 1.63827 |
| CSN4 | 2 | 0.54 | 0.02 | 1.63827 |
| UGPA | 3 | 0.52 | 0.02 | 1.63827 |
| ADK | 3 | 0.52 | 0.02 | 1.63827 |
| DEST | 3 | 0.50 | 0.02 | 1.63827 |
| EMAL2 | 4 | 0.60 | 0.02 | 1.63827 |
| SYK | 6 | 0.61 | 0.02 | 1.63827 |
| MAT2B | 6 | 0.47 | 0.02 | 1.63827 |
| PSME2 | 10 | 0.60 | 0.02 | 1.63827 |
| RHG01 | 1 | 0.81 | 0.02 | 1.61979 |
| DHSO | 2 | 0.47 | 0.02 | 1.61979 |
| DHPR | 4 | 0.41 | 0.02 | 1.61979 |
| GBP3 | 1 | 21.3 | 0.03 | 1.60206 |
| MATR3 | 1 | 0.53 | 0.03 | 1.60206 |
| EF1D | 1 | 0.49 | 0.03 | 1.60206 |
| PLST | 1 | 0.41 | 0.03 | 1.60206 |
| DP13A | 2 | 0.42 | 0.03 | 1.60206 |
| H33 | 3 | 0.60 | 0.03 | 1.60206 |
| MIF | 3 | 0.51 | 0.03 | 1.60206 |
| TCPE | 9 | 0.61 | 0.03 | 1.60206 |
| CAND1 | 10 | 0.52 | 0.03 | 1.60206 |
| SYVC | 1 | 0.49 | 0.03 | 1.58503 |
| H13 | 2 | 0.31 | 0.03 | 1.58503 |
| SND1 | 9 | 0.68 | 0.03 | 1.58503 |
| SRP72 | 1 | 5.71 | 0.03 | 1.56864 |
| VATE1 | 1 | 0.56 | 0.03 | 1.56864 |
| RL7 | 2 | 0.57 | 0.03 | 1.56864 |
| GSLG1 | 3 | 3.18 | 0.03 | 1.56864 |
| F16P1 | 6 | 0.57 | 0.03 | 1.56864 |
| TRYP_PIG | 10 | 1.69 | 0.03 | 1.56864 |
| ALDOA | 22 | 0.66 | 0.03 | 1.56864 |
| CSN5 | 1 | 0.57 | 0.03 | 1.55284 |
| DNJC9 | 1 | 0.49 | 0.03 | 1.55284 |
| RUXF | 2 | 0.50 | 0.03 | 1.55284 |
| NP1L4 | 5 | 0.48 | 0.03 | 1.55284 |
| AK1A1 | 7 | 0.60 | 0.03 | 1.55284 |
| PTN6 | 16 | 0.64 | 0.03 | 1.55284 |
| TALDO | 16 | 0.59 | 0.03 | 1.55284 |
| TPM2 | 1 | 0.45 | 0.03 | 1.5376 |
| PTPA | 5 | 0.58 | 0.03 | 1.5376 |
| ACTB | 21 | 0.70 | 0.03 | 1.5376 |
| UBL4A | 1 | 0.37 | 0.03 | 1.52288 |
| MTAP | 1 | 0.36 | 0.03 | 1.52288 |
| AP1B1 | 1 | 0.16 | 0.03 | 1.52288 |

| | | | | |
|---|---|---|---|---|
| GBRL2 | 2 | 0.57 | 0.03 | 1.52288 |
| LZTL1 | 2 | 0.43 | 0.03 | 1.52288 |
| CD2AP | 5 | 0.58 | 0.03 | 1.52288 |
| RACK1 | 8 | 0.63 | 0.03 | 1.52288 |
| NPC2 | 1 | 2.59 | 0.03 | 1.50864 |
| CRYM | 2 | 0.26 | 0.03 | 1.50864 |
| GDIR1 | 7 | 0.61 | 0.03 | 1.50864 |
| RLA0 | 9 | 0.57 | 0.03 | 1.49485 |
| TPM3 | 20 | 0.65 | 0.03 | 1.49485 |
| STAT3 | 1 | 6.31 | 0.03 | 1.48149 |
| LXN | 1 | 0.45 | 0.03 | 1.48149 |
| TMA7 | 2 | 0.57 | 0.03 | 1.48149 |
| EIF3G | 2 | 0.53 | 0.03 | 1.48149 |
| IMPA1 | 7 | 0.50 | 0.03 | 1.48149 |
| LKHA4 | 18 | 0.61 | 0.03 | 1.48149 |
| TXD17 | 2 | 0.58 | 0.03 | 1.46852 |
| GFRP | 2 | 0.26 | 0.03 | 1.46852 |
| SPF27 | 1 | 0.53 | 0.04 | 1.45593 |
| SUMO3 | 1 | 0.51 | 0.04 | 1.45593 |
| SC31A | 2 | 0.57 | 0.04 | 1.45593 |
| TSP1 | 1 | 0.74 | 0.04 | 1.4437 |
| H2A1B | 1 | 0.58 | 0.04 | 1.4437 |
| OAS3 | 2 | 2.91 | 0.04 | 1.4437 |
| PRDX5 | 3 | 0.56 | 0.04 | 1.4437 |
| PSB6 | 3 | 0.51 | 0.04 | 1.4437 |
| OTUB1 | 4 | 0.55 | 0.04 | 1.4437 |
| DNJC8 | 7 | 0.53 | 0.04 | 1.4437 |
| K2C6B | 9 | 0.35 | 0.04 | 1.4437 |
| RCC2 | 13 | 0.57 | 0.04 | 1.4437 |
| PHF5A | 1 | 0.26 | 0.04 | 1.4318 |
| CDC42 | 4 | 0.65 | 0.04 | 1.4318 |
| SYSC | 5 | 0.65 | 0.04 | 1.4318 |
| PGM1 | 9 | 0.54 | 0.04 | 1.42022 |
| ARC1A | 2 | 0.55 | 0.04 | 1.40894 |
| DAZP1 | 2 | 0.37 | 0.04 | 1.40894 |
| HLAB | 4 | 2.56 | 0.04 | 1.40894 |
| IDHC | 9 | 0.64 | 0.04 | 1.40894 |
| CADH1 | 1 | 3.22 | 0.04 | 1.39794 |
| DENR | 1 | 0.56 | 0.04 | 1.39794 |
| RGS10 | 1 | 0.56 | 0.04 | 1.39794 |
| SASH3 | 1 | 0.54 | 0.04 | 1.39794 |
| HNRPQ | 4 | 0.65 | 0.04 | 1.39794 |
| ARK72 | 1 | 0.41 | 0.04 | 1.38722 |
| SPB6 | 2 | 0.55 | 0.04 | 1.38722 |

| | | | | |
|---|---|---|---|---|
| ECHD1 | 2 | 0.49 | 0.04 | 1.38722 |
| SMC1A | 3 | 0.50 | 0.04 | 1.38722 |
| CBX5 | 3 | 0.46 | 0.04 | 1.38722 |
| LPXN | 1 | 0.62 | 0.04 | 1.37675 |
| EIF3F | 2 | 0.66 | 0.04 | 1.37675 |
| DDX5 | 2 | 0.60 | 0.04 | 1.37675 |
| ML12A | 5 | 0.75 | 0.04 | 1.37675 |
| PP1A | 7 | 0.63 | 0.04 | 1.37675 |
| UFC1 | 3 | 0.53 | 0.04 | 1.35655 |
| TPM3 | 1 | 0.67 | 0.05 | 1.34679 |
| FMC1 | 1 | 0.64 | 0.05 | 1.34679 |
| AP3D1 | 2 | 0.17 | 0.05 | 1.34679 |
| PTMA | 4 | 3.23 | 0.05 | 1.34679 |
| IMA3 | 1 | 2.33 | 0.05 | 1.33724 |
| RBBP7 | 1 | 0.55 | 0.05 | 1.33724 |
| VAMP7 | 1 | 0.41 | 0.05 | 1.33724 |
| STRAP | 3 | 0.58 | 0.05 | 1.33724 |
| RAB8A | 3 | 0.57 | 0.05 | 1.33724 |
| FKB1A | 3 | 0.41 | 0.05 | 1.33724 |
| IF16 | 5 | 0.56 | 0.05 | 1.33724 |
| HSP74 | 12 | 0.62 | 0.05 | 1.33724 |
| TFIP8 | 1 | 0.64 | 0.05 | 1.3279 |
| MOB1A | 1 | 0.40 | 0.05 | 1.3279 |
| PEBB | 1 | 0.18 | 0.05 | 1.3279 |
| GLRX3 | 1 | 3.93 | 0.05 | 1.31876 |
| PRPS2 | 3 | 0.56 | 0.05 | 1.31876 |
| G3BP1 | 5 | 0.67 | 0.05 | 1.31876 |
| GSTO1 | 6 | 0.57 | 0.05 | 1.31876 |
| GYS1 | 1 | 3.86 | 0.05 | 1.3098 |
| CSTF2 | 1 | 0.51 | 0.05 | 1.3098 |
| LC7L3 | 1 | 0.50 | 0.05 | 1.3098 |
| PP2AA | 3 | 0.54 | 0.05 | 1.3098 |
| CSN2 | 1 | 0.49 | 0.05 | 1.30103 |
| PURA | 2 | 0.58 | 0.05 | 1.30103 |
| ACLY | 14 | 0.66 | 0.05 | 1.30103 |
| HNRPK | 15 | 0.71 | 0.05 | 1.30103 |
| CY24B | 1 | 0.58 | 0.05 | 1.29243 |
| RBM12 | 1 | 0.45 | 0.05 | 1.29243 |
| SMD2 | 2 | 0.50 | 0.05 | 1.29243 |
| H2A2B | 4 | 0.56 | 0.05 | 1.29243 |
| BCLF1 | 2 | 0.55 | 0.05 | 1.284 |
| PAIP1 | 3 | 0.54 | 0.05 | 1.284 |
| SRSF6 | 2 | 3.15 | 0.05 | 1.27572 |
| EIF3L | 6 | 0.59 | 0.05 | 1.27572 |

| | | | | |
|---|---|---|---|---|
| MX1 | 15 | 2.46 | 0.05 | 1.27572 |
| FUS | 1 | 2.11 | 0.05 | 1.26761 |
| SYCC | 1 | 0.56 | 0.05 | 1.26761 |
| DCTD | 1 | 0.43 | 0.05 | 1.26761 |
| ARPC5 | 4 | 0.51 | 0.05 | 1.26761 |
| GRB2 | 7 | 0.66 | 0.05 | 1.26761 |
| CFAH | 1 | 1.55 | 0.06 | 1.25964 |
| HMCN1 | 1 | 1.32 | 0.06 | 1.25964 |
| PP1B | 1 | 0.62 | 0.06 | 1.25964 |
| DC1I2 | 1 | 0.27 | 0.06 | 1.25964 |
| NOP56 | 2 | 0.55 | 0.06 | 1.25964 |
| TSTD1 | 2 | 0.51 | 0.06 | 1.25964 |
| IST1 | 2 | 0.57 | 0.06 | 1.25181 |
| 1433B | 1 | 0.62 | 0.06 | 1.24413 |
| LYRIC | 1 | 0.11 | 0.06 | 1.24413 |
| NUCKS | 2 | 0.40 | 0.06 | 1.24413 |
| ARHL2 | 3 | 0.52 | 0.06 | 1.23657 |
| GMPPA | 1 | 0.51 | 0.06 | 1.22915 |
| EXOS4 | 1 | 0.24 | 0.06 | 1.22915 |
| THOC2 | 1 | 0.35 | 0.06 | 1.22185 |
| IF2P | 2 | 0.37 | 0.06 | 1.22185 |
| VPS35 | 4 | 0.64 | 0.06 | 1.22185 |
| CFAB | 1 | 7.00 | 0.06 | 1.20761 |
| SRGN | 1 | 116 | 0.06 | 1.20066 |
| DCUP | 1 | 0.27 | 0.06 | 1.20066 |
| OLA1 | 2 | 0.55 | 0.06 | 1.20066 |
| GBB1 | 9 | 0.67 | 0.06 | 1.20066 |
| HARS1 | 11 | 0.59 | 0.06 | 1.20066 |
| XRCC5 | 14 | 0.67 | 0.06 | 1.20066 |
| SYWC | 15 | 1.76 | 0.06 | 1.20066 |
| HDGR2 | 1 | 0.10 | 0.06 | 1.19382 |
| CPPED | 2 | 0.52 | 0.06 | 1.19382 |
| RRBP1 | 21 | 0.62 | 0.06 | 1.19382 |
| PYGB | 1 | 0.62 | 0.07 | 1.18709 |
| SYDC | 5 | 0.71 | 0.07 | 1.18709 |
| UBL5 | 1 | 1.86 | 0.07 | 1.18046 |
| K1C16 | 11 | 0.23 | 0.07 | 1.18046 |
| LA | 9 | 0.63 | 0.07 | 1.17393 |
| UB2V2 | 1 | 0.48 | 0.07 | 1.1549 |
| GGCT | 3 | 0.49 | 0.07 | 1.1549 |
| NUCL | 17 | 0.61 | 0.07 | 1.1549 |
| DCD | 1 | 1.47 | 0.07 | 1.14874 |
| PFD3 | 2 | 0.52 | 0.07 | 1.14874 |
| OSBP1 | 1 | 0.51 | 0.07 | 1.14267 |

| | | | | |
|---|---|---|---|---|
| ERF1 | 2 | 0.55 | 0.07 | 1.14267 |
| NP1L1 | 5 | 0.58 | 0.07 | 1.14267 |
| EPS15 | 1 | 0.44 | 0.07 | 1.13668 |
| RTRAF | 2 | 0.61 | 0.07 | 1.13077 |
| SRSF1 | 7 | 0.65 | 0.07 | 1.13077 |
| EIF3J | 1 | 0.53 | 0.08 | 1.12494 |
| ACTG | 2 | 0.39 | 0.08 | 1.12494 |
| CBR1 | 3 | 0.34 | 0.08 | 1.12494 |
| XRCC6 | 12 | 0.66 | 0.08 | 1.12494 |
| PUR6 | 5 | 0.62 | 0.08 | 1.11919 |
| TRXR1 | 7 | 0.61 | 0.08 | 1.11919 |
| IF4G2 | 1 | 23.0 | 0.08 | 1.10791 |
| 3BP1 | 2 | 2.05 | 0.08 | 1.10791 |
| TBCA | 4 | 0.68 | 0.08 | 1.10791 |
| IF4G1 | 1 | 4.24 | 0.08 | 1.10237 |
| CCAR2 | 1 | 0.60 | 0.08 | 1.10237 |
| U2AF2 | 2 | 0.53 | 0.08 | 1.09691 |
| TIMP1 | 3 | 0.64 | 0.08 | 1.09691 |
| FLII | 1 | 0.75 | 0.08 | 1.09151 |
| ELOC | 1 | 0.58 | 0.08 | 1.09151 |
| LDHA | 16 | 0.61 | 0.08 | 1.09151 |
| RMD1 | 1 | 0.43 | 0.08 | 1.08619 |
| MAP1A | 1 | 0.35 | 0.08 | 1.08619 |
| NECP2 | 1 | 0.67 | 0.08 | 1.08092 |
| 1433G | 6 | 0.62 | 0.08 | 1.08092 |
| PTMS | 2 | 0.43 | 0.08 | 1.07572 |
| ANR44 | 1 | 0.65 | 0.09 | 1.07058 |
| G3P | 18 | 0.68 | 0.09 | 1.0655 |
| ERP29 | 6 | 0.73 | 0.09 | 1.05552 |
| TKT | 29 | 0.65 | 0.09 | 1.05552 |
| STK24 | 1 | 0.54 | 0.09 | 1.05061 |
| HAT1 | 1 | 0.38 | 0.09 | 1.05061 |
| THIC | 2 | 0.58 | 0.09 | 1.05061 |
| PGAM1 | 16 | 0.65 | 0.09 | 1.05061 |
| FNTA | 1 | 0.55 | 0.09 | 1.04576 |
| B4GT1 | 2 | 2.19 | 0.09 | 1.04576 |
| ENOPH | 2 | 0.58 | 0.09 | 1.04576 |
| TCP4 | 8 | 0.58 | 0.09 | 1.04096 |
| TOP1 | 1 | 0.46 | 0.09 | 1.03621 |
| DNJA2 | 2 | 1.59 | 0.09 | 1.03621 |
| AMPB | 8 | 0.58 | 0.09 | 1.03152 |
| PUR8 | 1 | 0.54 | 0.09 | 1.02687 |
| PLEK | 7 | 1.90 | 0.09 | 1.02687 |
| ILEU | 13 | 0.72 | 0.09 | 1.02687 |

| | | | | |
|---|---|---|---|---|
| TRY1 | 1 | 1.49 | 0.10 | 1.02228 |
| CK054 | 1 | 0.41 | 0.10 | 1.01773 |
| DNM1L | 1 | 0.71 | 0.10 | 1.01323 |
| DDX23 | 1 | 0.19 | 0.10 | 1.01323 |
| PDIA3 | 13 | 0.72 | 0.10 | 1.01323 |
| SF3B2 | 4 | 0.62 | 0.10 | 1.00877 |
| NDRG1 | 2 | 0.39 | 0.10 | 1.00436 |
| HLAC | 3 | 3.17 | 0.10 | 1 |
| H31 | 2 | 0.65 | 0.10 | 0.99568 |
| CUTA | 3 | 0.62 | 0.10 | 0.9914 |
| U5S1 | 1 | 0.60 | 0.10 | 0.98716 |
| NRDC | 1 | 0.49 | 0.10 | 0.98716 |
| PIMT | 4 | 0.38 | 0.10 | 0.98716 |
| PLPHP | 2 | 0.63 | 0.10 | 0.98297 |
| TEBP | 3 | 1.84 | 0.10 | 0.98297 |
| ALDOC | 3 | 0.65 | 0.10 | 0.98297 |
| EF2 | 32 | 0.73 | 0.10 | 0.98297 |
| E2AK2 | 1 | 65.3 | 0.11 | 0.97469 |
| RS30 | 1 | 2.38 | 0.11 | 0.97469 |
| UBP14 | 4 | 0.69 | 0.11 | 0.97469 |
| MYDGF | 2 | 0.44 | 0.11 | 0.96257 |
| FKBP4 | 5 | 0.63 | 0.11 | 0.96257 |
| GDIR2 | 10 | 0.70 | 0.11 | 0.96257 |
| B2CL2 | 1 | 0.30 | 0.11 | 0.95861 |
| K2C6C | 1 | 0.42 | 0.11 | 0.95468 |
| FUMH | 2 | 0.66 | 0.11 | 0.95078 |
| DUT | 1 | 2.73 | 0.12 | 0.9393 |
| SPTN1 | 4 | 0.61 | 0.12 | 0.9393 |
| IF2A | 3 | 0.65 | 0.12 | 0.93554 |
| AP2A1 | 5 | 0.61 | 0.12 | 0.93554 |
| WDR48 | 1 | 0.25 | 0.12 | 0.93181 |
| BPNT1 | 2 | 0.66 | 0.12 | 0.92812 |
| TBA1A | 1 | 1.82 | 0.12 | 0.92445 |
| ULA1 | 1 | 0.52 | 0.12 | 0.92445 |
| INS | 2 | 1.16 | 0.12 | 0.92445 |
| IL16 | 2 | 0.62 | 0.12 | 0.92445 |
| GLRX1 | 2 | 0.57 | 0.12 | 0.92445 |
| HPRT | 3 | 0.62 | 0.12 | 0.92445 |
| GUAA | 1 | 0.44 | 0.12 | 0.91009 |
| SCRN1 | 3 | 1.55 | 0.12 | 0.91009 |
| HORN | 4 | 0.43 | 0.12 | 0.91009 |
| MESD | 1 | 0.68 | 0.12 | 0.90658 |
| PTGDS | 3 | 3.78 | 0.12 | 0.90658 |
| TSSK4 | 1 | 1.25 | 0.13 | 0.90309 |

| | | | | |
|---|---|---|---|---|
| ELMO1 | 2 | 0.74 | 0.13 | 0.90309 |
| FHL1 | 3 | 0.22 | 0.13 | 0.90309 |
| ABI1 | 1 | 0.59 | 0.13 | 0.89963 |
| CAB39 | 1 | 0.51 | 0.13 | 0.8962 |
| SH3L1 | 2 | 0.62 | 0.13 | 0.8962 |
| EDF1 | 3 | 0.64 | 0.13 | 0.8962 |
| LSP1 | 8 | 0.64 | 0.13 | 0.8962 |
| EIF1 | 1 | 1.51 | 0.13 | 0.89279 |
| 5NTC | 1 | 0.52 | 0.13 | 0.89279 |
| CCL19 | 1 | 1,000 | 0.13 | 0.88941 |
| PTPRS | 5 | 1.54 | 0.13 | 0.88273 |
| CIRBP | 1 | 0.57 | 0.13 | 0.87943 |
| CYFP2 | 3 | 0.68 | 0.13 | 0.87943 |
| DCTN1 | 3 | 0.55 | 0.13 | 0.87943 |
| UBP5 | 3 | 0.52 | 0.13 | 0.87615 |
| LEG1 | 8 | 0.69 | 0.13 | 0.87615 |
| PABP1 | 10 | 0.71 | 0.13 | 0.8729 |
| FKBP3 | 2 | 0.64 | 0.14 | 0.86967 |
| K1C14 | 5 | 0.19 | 0.14 | 0.86967 |
| PUR2 | 6 | 1.69 | 0.14 | 0.86967 |
| PPCE | 2 | 0.66 | 0.14 | 0.85699 |
| QOR | 4 | 0.46 | 0.14 | 0.85699 |
| NMT1 | 3 | 0.66 | 0.14 | 0.85387 |
| K2C5 | 14 | 0.29 | 0.14 | 0.85387 |
| AP3S1 | 1 | 0.59 | 0.14 | 0.84466 |
| ALDR | 5 | 0.60 | 0.14 | 0.84466 |
| RBM14 | 1 | 1.79 | 0.15 | 0.83565 |
| PSPC1 | 1 | 0.63 | 0.15 | 0.83565 |
| CHM4B | 2 | 1.48 | 0.15 | 0.83565 |
| TSYL2 | 1 | 2.14 | 0.15 | 0.83268 |
| NACAM | 2 | 0.69 | 0.15 | 0.83268 |
| RS20 | 2 | 0.70 | 0.15 | 0.82974 |
| DDX6 | 4 | 0.72 | 0.15 | 0.82974 |
| TCEA1 | 4 | 0.45 | 0.15 | 0.82974 |
| KSYK | 1 | 1.48 | 0.15 | 0.82681 |
| PCBP2 | 6 | 0.76 | 0.15 | 0.82681 |
| IF6 | 4 | 0.81 | 0.15 | 0.82391 |
| HMGB1 | 6 | 0.59 | 0.15 | 0.82102 |
| EXOS9 | 1 | 0.64 | 0.15 | 0.81816 |
| MICA1 | 1 | 0.60 | 0.15 | 0.81816 |
| DCXR | 2 | 0.67 | 0.16 | 0.80967 |
| SPTB2 | 8 | 0.61 | 0.16 | 0.80967 |
| G6PI | 15 | 0.71 | 0.16 | 0.80967 |
| YBOX1 | 7 | 0.66 | 0.16 | 0.80688 |

| | | | | |
|---|---|---|---|---|
| IN35 | 1 | 1.32 | 0.16 | 0.80134 |
| AP1G2 | 2 | 0.31 | 0.16 | 0.7986 |
| BTF3 | 1 | 0.66 | 0.16 | 0.79588 |
| 4F2 | 1 | 0.55 | 0.16 | 0.79588 |
| K2C78 | 3 | 0.25 | 0.16 | 0.79588 |
| RPR1B | 1 | 0.60 | 0.16 | 0.79048 |
| RS8 | 7 | 0.66 | 0.16 | 0.79048 |
| ANXA5 | 17 | 0.78 | 0.16 | 0.79048 |
| ROA2 | 19 | 0.76 | 0.16 | 0.78781 |
| PPARD | 1 | 1.18 | 0.16 | 0.78516 |
| C19L1 | 1 | 0.43 | 0.16 | 0.78516 |
| MP2K2 | 1 | 1.60 | 0.17 | 0.78252 |
| HGFA | 1 | 1.65 | 0.17 | 0.77989 |
| MFAP1 | 1 | 0.51 | 0.17 | 0.77728 |
| SUMO1 | 1 | 0.45 | 0.17 | 0.77728 |
| S10AA | 2 | 0.69 | 0.17 | 0.77211 |
| IFNG | 1 | 24.2 | 0.17 | 0.76955 |
| AGFG1 | 2 | 0.60 | 0.17 | 0.76955 |
| RAB8B | 1 | 2.12 | 0.17 | 0.76195 |
| ACL6A | 1 | 0.64 | 0.17 | 0.75945 |
| ISOC1 | 2 | 0.55 | 0.17 | 0.75945 |
| WASP | 4 | 0.61 | 0.17 | 0.75945 |
| HV373 | 1 | 4.68 | 0.18 | 0.75696 |
| DESP | 3 | 0.16 | 0.18 | 0.75696 |
| HS90A | 33 | 0.69 | 0.18 | 0.75696 |
| RS27A | 2 | 0.64 | 0.18 | 0.74232 |
| ROA1 | 12 | 0.74 | 0.18 | 0.73993 |
| EIF2A | 1 | 0.41 | 0.18 | 0.73518 |
| RS16 | 4 | 0.61 | 0.18 | 0.73518 |
| THUM1 | 1 | 0.63 | 0.19 | 0.73283 |
| TIGAR | 1 | 0.61 | 0.19 | 0.73283 |
| UB2J1 | 1 | 0.11 | 0.19 | 0.73283 |
| PPM1F | 2 | 0.65 | 0.19 | 0.72584 |
| H32 | 2 | 0.73 | 0.19 | 0.7167 |
| PTN11 | 2 | 0.63 | 0.19 | 0.71444 |
| RS11 | 2 | 0.64 | 0.20 | 0.70997 |
| PCBP1 | 11 | 0.80 | 0.20 | 0.70997 |
| RL32 | 1 | 0.56 | 0.20 | 0.70553 |
| HGS | 1 | 0.67 | 0.20 | 0.70333 |
| RL17 | 1 | 0.65 | 0.20 | 0.69897 |
| TRA2B | 2 | 0.75 | 0.20 | 0.69897 |
| GMIP | 1 | 0.52 | 0.20 | 0.6968 |
| RL23A | 3 | 0.65 | 0.20 | 0.69465 |
| HEBP2 | 1 | 0.64 | 0.20 | 0.6925 |

| | | | | |
|---|---|---|---|---|
| **ABRX2** | **3** | **0.67** | **0.20** | **0.6925** |
| **K1C9** | **35** | **0.48** | **0.20** | **0.69037** |
| **K1C10** | **27** | **0.30** | **0.21** | **0.68613** |
| **PLCG2** | **1** | **0.65** | **0.21** | **0.68403** |
| **RU2A** | **1** | **0.51** | **0.21** | **0.67985** |
| **DSC1** | **1** | **0.23** | **0.21** | **0.67985** |
| **SARNP** | **2** | **0.68** | **0.21** | **0.67778** |
| **ADHX** | **5** | **0.64** | **0.21** | **0.67778** |
| **TPD54** | **1** | **0.69** | **0.21** | **0.67572** |
| **NIT2** | **1** | **0.49** | **0.21** | **0.67366** |
| **RL18** | **2** | **0.67** | **0.21** | **0.67366** |
| **GIT2** | **2** | **0.50** | **0.21** | **0.67162** |
| **CATG** | **1** | **2.80** | **0.21** | **0.66959** |
| **ZRAB2** | **1** | **0.60** | **0.22** | **0.66756** |
| **RS10** | **1** | **0.78** | **0.22** | **0.65758** |
| **HS71A** | **21** | **1.21** | **0.22** | **0.65758** |
| **K22E** | **20** | **0.31** | **0.22** | **0.65561** |
| **CYTC** | **3** | **0.75** | **0.22** | **0.65365** |
| **AHNK** | **5** | **0.74** | **0.22** | **0.65365** |
| **K2C1** | **38** | **0.37** | **0.22** | **0.65365** |
| **PSDE** | **1** | **0.77** | **0.22** | **0.64975** |
| **H2B1B** | **2** | **0.70** | **0.23** | **0.64589** |
| **SGTA** | **1** | **0.54** | **0.23** | **0.64016** |
| **BST2** | **2** | **2.46** | **0.23** | **0.63827** |
| **TBB6** | **4** | **0.61** | **0.23** | **0.63451** |
| **CCD50** | **2** | **0.60** | **0.23** | **0.63264** |
| **TYB4** | **1** | **0.78** | **0.24** | **0.62893** |
| **CIAO1** | **1** | **0.60** | **0.24** | **0.62893** |
| **EIF3M** | **2** | **0.67** | **0.24** | **0.62893** |
| **PFKAL** | **2** | **0.80** | **0.24** | **0.62709** |
| **BZW1** | **1** | **1.62** | **0.24** | **0.62525** |
| **DYH8** | **1** | **1.12** | **0.24** | **0.62525** |
| **BLMH** | **1** | **0.72** | **0.24** | **0.6216** |
| **RL36** | **1** | **0.60** | **0.24** | **0.6216** |
| **MTND** | **1** | **0.38** | **0.24** | **0.61798** |
| **TPP2** | **4** | **0.72** | **0.24** | **0.61618** |
| **LMNA** | **5** | **0.63** | **0.24** | **0.61618** |
| **HEM2** | **1** | **0.52** | **0.24** | **0.61439** |
| **DIAP1** | **4** | **0.78** | **0.24** | **0.61439** |
| **TPD52** | **1** | **0.44** | **0.24** | **0.61261** |
| **RS3A** | **9** | **0.72** | **0.24** | **0.61261** |
| **RL27** | **2** | **0.72** | **0.25** | **0.60906** |
| **MYPT1** | **1** | **0.59** | **0.25** | **0.60555** |
| **RS3** | **6** | **0.72** | **0.25** | **0.60555** |

| | | | | |
|---|---|---|---|---|
| **FBX6** | **1** | **2.89** | **0.25** | **0.60033** |
| **SGT1** | **2** | **0.71** | **0.25** | **0.59688** |
| **AMY1** | **2** | **0.77** | **0.26** | **0.59346** |
| **PSMD9** | **3** | **0.75** | **0.26** | **0.59176** |
| **PA1B2** | **1** | **0.77** | **0.26** | **0.59007** |
| **LTOR1** | **1** | **0.48** | **0.26** | **0.59007** |
| **NIBA1** | **1** | **2.57** | **0.26** | **0.5867** |
| **RAGP1** | **2** | **0.65** | **0.26** | **0.5867** |
| **SORCN** | **1** | **0.73** | **0.26** | **0.58503** |
| **PTGR1** | **2** | **0.37** | **0.26** | **0.58503** |
| **TXNL1** | **2** | **0.76** | **0.26** | **0.5817** |
| **SAMH1** | **19** | **0.85** | **0.27** | **0.57675** |
| **USP9X** | **1** | **0.55** | **0.27** | **0.57349** |
| **OSTF1** | **2** | **0.79** | **0.27** | **0.56543** |
| **AL9A1** | **1** | **0.63** | **0.27** | **0.56225** |
| **RL4** | **5** | **0.72** | **0.28** | **0.55909** |
| **HV64D** | **1** | **0.72** | **0.28** | **0.55596** |
| **RPAB3** | **1** | **0.37** | **0.28** | **0.5544** |
| **PAIRB** | **2** | **0.79** | **0.28** | **0.5544** |
| **RL30** | **2** | **0.70** | **0.28** | **0.5544** |
| **PDIA1** | **12** | **0.72** | **0.28** | **0.55129** |
| **EZRI** | **17** | **0.73** | **0.28** | **0.54975** |
| **DYHC1** | **1** | **0.76** | **0.28** | **0.54668** |
| **PRS8** | **1** | **1.43** | **0.29** | **0.5391** |
| **RL11** | **3** | **1.38** | **0.29** | **0.5391** |
| **CSK2B** | **1** | **0.76** | **0.29** | **0.5376** |
| **HMGN2** | **2** | **0.65** | **0.30** | **0.52578** |
| **NONO** | **3** | **0.80** | **0.30** | **0.52578** |
| **ANXA6** | **6** | **0.78** | **0.30** | **0.52578** |
| **SYFA** | **1** | **0.71** | **0.30** | **0.52288** |
| **RL21** | **1** | **0.69** | **0.30** | **0.52288** |
| **RL10A** | **5** | **0.78** | **0.30** | **0.52288** |
| **PSME3** | **2** | **0.71** | **0.30** | **0.52143** |
| **LV469** | **1** | **2.47** | **0.30** | **0.51999** |
| **PHP14** | **1** | **0.37** | **0.30** | **0.51999** |
| **CNTN1** | **2** | **1.08** | **0.30** | **0.51856** |
| **U2AF5** | **1** | **0.74** | **0.30** | **0.51713** |
| **SKP1** | **1** | **0.48** | **0.31** | **0.5157** |
| **A2MG** | **1** | **1.41** | **0.31** | **0.51286** |
| **RS7** | **2** | **0.67** | **0.31** | **0.51145** |
| **OXSR1** | **1** | **0.72** | **0.31** | **0.50864** |
| **RL35A** | **1** | **2.00** | **0.31** | **0.50724** |
| **HYOU1** | **3** | **1.34** | **0.31** | **0.50724** |
| **BGH3** | **7** | **0.81** | **0.31** | **0.50585** |

| | | | | |
|---|---|---|---|---|
| CO4A | 2 | 1.16 | 0.31 | 0.50446 |
| ABCE1 | 1 | 0.61 | 0.31 | 0.50307 |
| PIN1 | 1 | 0.66 | 0.32 | 0.50169 |
| HNRH1 | 9 | 0.84 | 0.32 | 0.50169 |
| SPB12 | 1 | 0.40 | 0.32 | 0.50031 |
| LRRF1 | 1 | 0.86 | 0.32 | 0.49894 |
| GILT | 1 | 1.34 | 0.32 | 0.49757 |
| RAB10 | 1 | 0.79 | 0.32 | 0.48945 |
| NPL4 | 1 | 0.75 | 0.32 | 0.48945 |
| DYN2 | 3 | 0.70 | 0.32 | 0.48945 |
| RL3 | 6 | 0.75 | 0.32 | 0.48945 |
| PDCD6 | 1 | 0.72 | 0.33 | 0.48812 |
| BLVRB | 1 | 1.71 | 0.33 | 0.48413 |
| RS4X | 2 | 0.67 | 0.33 | 0.48413 |
| HS90B | 17 | 0.79 | 0.33 | 0.4828 |
| CH60 | 3 | 1.63 | 0.33 | 0.48149 |
| SAFB1 | 2 | 0.67 | 0.33 | 0.48017 |
| URP2 | 11 | 0.81 | 0.33 | 0.48017 |
| FRIL | 1 | 1.88 | 0.33 | 0.47886 |
| AIF1 | 2 | 0.81 | 0.33 | 0.47625 |
| SAP18 | 1 | 0.63 | 0.34 | 0.47237 |
| PSIP1 | 1 | 0.50 | 0.34 | 0.47237 |
| S10AB | 1 | 0.75 | 0.34 | 0.46852 |
| GRHPR | 1 | 0.68 | 0.34 | 0.46725 |
| UBR4 | 1 | 2.28 | 0.34 | 0.46597 |
| GLU2B | 1 | 0.74 | 0.34 | 0.46597 |
| NUDC | 1 | 0.74 | 0.34 | 0.46471 |
| NT5D1 | 1 | 0.67 | 0.34 | 0.46471 |
| USO1 | 7 | 0.78 | 0.34 | 0.46471 |
| CAR11 | 1 | 0.46 | 0.35 | 0.46218 |
| IPP2B | 1 | 1.98 | 0.35 | 0.46092 |
| HSPB1 | 1 | 0.64 | 0.35 | 0.45842 |
| PSD11 | 3 | 1.31 | 0.35 | 0.45842 |
| FPPS | 1 | 1.69 | 0.35 | 0.45223 |
| FYB1 | 1 | 0.41 | 0.35 | 0.451 |
| RHG25 | 1 | 0.71 | 0.36 | 0.44855 |
| AK1C3 | 3 | 0.62 | 0.36 | 0.44855 |
| DRB5 | 2 | 1.51 | 0.36 | 0.44249 |
| ITB2 | 3 | 1.53 | 0.36 | 0.4389 |
| RS26 | 2 | 0.65 | 0.37 | 0.43533 |
| CLIC1 | 12 | 0.82 | 0.37 | 0.43533 |
| NNRE | 1 | 0.66 | 0.37 | 0.43297 |
| RL15 | 1 | 0.63 | 0.37 | 0.43297 |
| THIO | 5 | 1.25 | 0.37 | 0.43297 |

| | | | | |
|---|---|---|---|---|
| **COMD2** | **1** | **1.44** | **0.37** | **0.4318** |
| **RBM3** | **1** | **0.56** | **0.37** | **0.4318** |
| **F120A** | **1** | **2.39** | **0.37** | **0.43063** |
| **CFAI** | **4** | **1.09** | **0.37** | **0.43063** |
| **G6PD** | **2** | **0.80** | **0.37** | **0.42829** |
| **MOES** | **43** | **0.82** | **0.38** | **0.42597** |
| **CAPR1** | **1** | **0.80** | **0.38** | **0.41908** |
| **RL7A** | **2** | **0.74** | **0.38** | **0.41908** |
| **KV315** | **2** | **1.40** | **0.38** | **0.41794** |
| **VPS29** | **1** | **0.66** | **0.38** | **0.4168** |
| **IGJ** | **8** | **1.28** | **0.38** | **0.41567** |
| **PSMD2** | **3** | **0.82** | **0.39** | **0.41454** |
| **PP1R7** | **1** | **0.68** | **0.39** | **0.41117** |
| **LV310** | **1** | **1.24** | **0.39** | **0.40894** |
| **RS2** | **2** | **0.81** | **0.39** | **0.40561** |
| **PTPRC** | **2** | **0.69** | **0.40** | **0.4023** |
| **PSB5** | **1** | **0.56** | **0.40** | **0.40121** |
| **UBP15** | **1** | **1.40** | **0.40** | **0.39903** |
| **KPYM** | **28** | **0.85** | **0.40** | **0.39903** |
| **KV401** | **2** | **1.52** | **0.40** | **0.39794** |
| **SAMP** | **1** | **1.14** | **0.40** | **0.39577** |
| **BIN2** | **4** | **0.77** | **0.40** | **0.39577** |
| **IMDH2** | **1** | **1.30** | **0.40** | **0.39469** |
| **JAML** | **1** | **0.68** | **0.40** | **0.39469** |
| **CO6** | **1** | **1.71** | **0.40** | **0.39362** |
| **UBE2K** | **1** | **0.61** | **0.41** | **0.39147** |
| **EF1B** | **3** | **0.79** | **0.41** | **0.38722** |
| **P85A** | **1** | **0.67** | **0.42** | **0.38091** |
| **ZCCHV** | **2** | **1.41** | **0.42** | **0.38091** |
| **SC23B** | **4** | **0.83** | **0.42** | **0.38091** |
| **HNRPF** | **5** | **0.84** | **0.42** | **0.38091** |
| **FAS** | **2** | **1.33** | **0.42** | **0.37469** |
| **RL19** | **2** | **0.75** | **0.42** | **0.37469** |
| **PPIB** | **7** | **0.82** | **0.42** | **0.37469** |
| **RAVR1** | **1** | **0.62** | **0.42** | **0.37366** |
| **RS17** | **3** | **0.75** | **0.42** | **0.37366** |
| **ALBU_BOVIN** | **95** | **1.08** | **0.42** | **0.37263** |
| **NASP** | **5** | **0.82** | **0.43** | **0.37059** |
| **ANT3** | **4** | **1.16** | **0.43** | **0.36552** |
| **PP1G** | **1** | **0.69** | **0.43** | **0.36351** |
| **LAMC1** | **3** | **0.78** | **0.44** | **0.36051** |
| **NUDC1** | **1** | **0.68** | **0.44** | **0.35952** |
| **IDHP** | **1** | **1.71** | **0.44** | **0.35853** |
| **TES** | **1** | **0.79** | **0.44** | **0.35754** |

| | | | | |
|---|---|---|---|---|
| PPIH | 1 | 0.70 | 0.44 | 0.35458 |
| S10A4 | 3 | 0.83 | 0.45 | 0.35164 |
| UBE3A | 1 | 0.70 | 0.45 | 0.35067 |
| AB1IP | 2 | 0.81 | 0.45 | 0.34486 |
| PDIA6 | 4 | 0.81 | 0.46 | 0.33724 |
| SPB9 | 12 | 0.78 | 0.47 | 0.33255 |
| UBA6 | 1 | 0.49 | 0.47 | 0.33068 |
| KVD11 | 2 | 1.52 | 0.47 | 0.32606 |
| FA98B | 2 | 0.71 | 0.48 | 0.32148 |
| UBCP1 | 1 | 0.69 | 0.48 | 0.31695 |
| FLNA | 49 | 0.85 | 0.48 | 0.31695 |
| TCOF | 4 | 1.16 | 0.48 | 0.31515 |
| TCTP | 3 | 0.84 | 0.49 | 0.31247 |
| EF1A1 | 19 | 1.13 | 0.49 | 0.31158 |
| KV116 | 1 | 1.94 | 0.49 | 0.31069 |
| RINI | 10 | 0.86 | 0.49 | 0.30715 |
| IF4A1 | 9 | 1.19 | 0.50 | 0.30364 |
| RAP1A | 1 | 0.82 | 0.50 | 0.3019 |
| FETUA | 1 | 1.22 | 0.50 | 0.29843 |
| IREB2 | 1 | 0.75 | 0.50 | 0.29843 |
| RALY | 2 | 1.16 | 0.50 | 0.29757 |
| PSB7 | 1 | 0.61 | 0.51 | 0.29585 |
| MPRI | 1 | 1.43 | 0.51 | 0.29328 |
| APOH | 1 | 1.09 | 0.51 | 0.29328 |
| CREG1 | 1 | 1.20 | 0.51 | 0.29158 |
| RS6 | 3 | 0.83 | 0.52 | 0.28735 |
| KAP2 | 1 | 0.84 | 0.52 | 0.28567 |
| ANXA2 | 16 | 0.90 | 0.52 | 0.28567 |
| PSMD3 | 3 | 0.85 | 0.52 | 0.284 |
| SF3A3 | 1 | 0.72 | 0.52 | 0.28316 |
| EHD1 | 4 | 0.79 | 0.52 | 0.28067 |
| ACAP1 | 1 | 0.47 | 0.53 | 0.27901 |
| GARS | 1 | 0.83 | 0.53 | 0.27737 |
| MAP4 | 4 | 0.79 | 0.53 | 0.27737 |
| MYO1E | 1 | 0.82 | 0.53 | 0.27654 |
| HV69D | 1 | 1.30 | 0.53 | 0.27572 |
| APT | 1 | 0.83 | 0.54 | 0.27084 |
| TRFL | 3 | 0.89 | 0.54 | 0.26922 |
| TBA1C | 18 | 1.17 | 0.54 | 0.26841 |
| DRA | 1 | 1.28 | 0.54 | 0.26761 |
| MY18A | 3 | 0.79 | 0.54 | 0.26761 |
| CYBP | 2 | 0.85 | 0.54 | 0.2668 |
| PCNA | 5 | 0.82 | 0.54 | 0.2652 |
| EMAL4 | 1 | 0.74 | 0.55 | 0.2636 |

| | | | | |
|---|---|---|---|---|
| ABC3C | 1 | 1.41 | 0.55 | 0.26122 |
| HV307 | 2 | 1.18 | 0.55 | 0.26043 |
| A1BG | 1 | 1.10 | 0.55 | 0.25727 |
| SYTC | 3 | 0.85 | 0.56 | 0.25259 |
| SPSY | 1 | 1.16 | 0.57 | 0.24718 |
| RS25 | 2 | 0.79 | 0.57 | 0.24718 |
| BIEA | 2 | 1.21 | 0.57 | 0.24336 |
| RS18 | 2 | 1.30 | 0.57 | 0.2426 |
| IMB1 | 5 | 0.85 | 0.57 | 0.24185 |
| SRSF7 | 2 | 0.86 | 0.57 | 0.24109 |
| HNRPM | 4 | 1.14 | 0.58 | 0.23958 |
| CD44 | 1 | 0.78 | 0.58 | 0.23882 |
| YTHD2 | 2 | 0.86 | 0.58 | 0.23882 |
| HV118 | 1 | 0.73 | 0.58 | 0.23508 |
| LV321 | 2 | 1.17 | 0.58 | 0.23508 |
| A2AP | 1 | 1.07 | 0.58 | 0.23359 |
| SRSF3 | 1 | 0.86 | 0.58 | 0.23359 |
| ACTN1 | 3 | 0.86 | 0.58 | 0.23359 |
| IGHG3 | 6 | 0.87 | 0.58 | 0.23359 |
| PSMD4 | 3 | 0.90 | 0.59 | 0.2321 |
| NHRF1 | 1 | 1.33 | 0.59 | 0.22621 |
| SRSF8 | 1 | 0.85 | 0.60 | 0.22113 |
| ERP44 | 1 | 1.19 | 0.60 | 0.2204 |
| HV321 | 2 | 1.27 | 0.60 | 0.21968 |
| LV147 | 1 | 1.19 | 0.60 | 0.21896 |
| RS5 | 4 | 0.92 | 0.60 | 0.21896 |
| PCNP | 1 | 1.18 | 0.61 | 0.21681 |
| GBB2 | 2 | 0.86 | 0.61 | 0.21467 |
| CO3A1 | 1 | 1.07 | 0.61 | 0.21325 |
| IGHM | 19 | 1.29 | 0.61 | 0.21325 |
| DDI2 | 1 | 1.16 | 0.61 | 0.21183 |
| METK2 | 3 | 0.93 | 0.62 | 0.21042 |
| HV102 | 1 | 0.64 | 0.62 | 0.20691 |
| GSH1 | 1 | 0.83 | 0.63 | 0.20412 |
| RL24 | 2 | 0.84 | 0.63 | 0.20412 |
| H2AZ | 2 | 0.77 | 0.63 | 0.20273 |
| SYAC | 5 | 0.80 | 0.63 | 0.20273 |
| TRIR | 1 | 1.28 | 0.63 | 0.19928 |
| IF4B | 3 | 1.17 | 0.63 | 0.1986 |
| ARI1 | 1 | 1.49 | 0.64 | 0.19723 |
| DDX42 | 2 | 1.22 | 0.64 | 0.19723 |
| RL37 | 1 | 1.39 | 0.64 | 0.1945 |
| KVD24 | 1 | 1.24 | 0.64 | 0.19246 |
| AOC3 | 1 | 0.89 | 0.65 | 0.18977 |

| | | | | |
|---|---|---|---|---|
| RL8 | 3 | 0.88 | 0.65 | 0.18709 |
| MK14 | 1 | 0.82 | 0.65 | 0.18575 |
| FIBB | 1 | 0.90 | 0.66 | 0.18243 |
| PDLI5 | 1 | 1.23 | 0.66 | 0.18177 |
| RANG | 2 | 1.11 | 0.66 | 0.18111 |
| LV151 | 1 | 1.35 | 0.67 | 0.17718 |
| PDIA4 | 10 | 1.11 | 0.67 | 0.17522 |
| PK3CD | 1 | 0.82 | 0.67 | 0.17393 |
| SF3A2 | 1 | 0.82 | 0.68 | 0.1707 |
| TETN | 3 | 0.96 | 0.68 | 0.16877 |
| S10A9 | 3 | 1.18 | 0.68 | 0.16813 |
| THBG | 2 | 1.07 | 0.68 | 0.16685 |
| RL28 | 1 | 1.37 | 0.68 | 0.16558 |
| IF5A1 | 4 | 0.92 | 0.69 | 0.16368 |
| RLA2 | 4 | 0.89 | 0.69 | 0.16368 |
| LASP1 | 2 | 0.87 | 0.69 | 0.16178 |
| TBB5 | 23 | 1.11 | 0.69 | 0.16052 |
| RL37A | 2 | 1.19 | 0.69 | 0.15927 |
| ICAL | 3 | 0.89 | 0.69 | 0.15864 |
| GAPR1 | 1 | 1.11 | 0.70 | 0.1549 |
| TBB4B | 1 | 0.90 | 0.70 | 0.1549 |
| PICAL | 1 | 0.86 | 0.70 | 0.15304 |
| PSMD6 | 1 | 0.91 | 0.71 | 0.15181 |
| NSF1C | 2 | 0.92 | 0.71 | 0.15181 |
| KV320 | 2 | 1.21 | 0.71 | 0.1512 |
| LAMB1 | 1 | 0.85 | 0.71 | 0.14997 |
| CLIC3 | 2 | 1.15 | 0.71 | 0.14935 |
| CORO7 | 1 | 1.22 | 0.71 | 0.14874 |
| DTD1 | 1 | 0.74 | 0.71 | 0.14874 |
| IGHG2 | 11 | 1.12 | 0.71 | 0.14874 |
| RTCB | 1 | 0.88 | 0.71 | 0.14813 |
| CSK | 1 | 1.11 | 0.71 | 0.14752 |
| PRS7 | 2 | 1.12 | 0.71 | 0.14752 |
| ENPL | 20 | 0.87 | 0.71 | 0.14752 |
| S10AC | 1 | 0.83 | 0.71 | 0.1463 |
| HV374 | 4 | 1.16 | 0.71 | 0.1463 |
| CAP7 | 1 | 1.21 | 0.72 | 0.14448 |
| LEUK | 1 | 0.87 | 0.72 | 0.14388 |
| KCY | 3 | 0.89 | 0.73 | 0.13966 |
| ITIH4 | 5 | 0.97 | 0.73 | 0.13847 |
| ACINU | 1 | 1.30 | 0.73 | 0.13549 |
| RECQ1 | 1 | 0.86 | 0.73 | 0.1343 |
| PEDF | 1 | 1.23 | 0.74 | 0.13312 |
| ZC3H4 | 1 | 0.86 | 0.74 | 0.13312 |

| | | | | |
|---|---|---|---|---|
| KV139 | 3 | 1.32 | 0.74 | 0.13312 |
| RUVB2 | 1 | 0.85 | 0.74 | 0.13194 |
| RL14 | 1 | 0.89 | 0.74 | 0.12901 |
| GIMA4 | 1 | 1.23 | 0.75 | 0.12784 |
| THRB | 1 | 1.15 | 0.75 | 0.12784 |
| IGLL5 | 9 | 1.22 | 0.75 | 0.12784 |
| PRDX3 | 1 | 1.10 | 0.76 | 0.12205 |
| IGKC | 7 | 1.12 | 0.76 | 0.11862 |
| HV330 | 4 | 1.16 | 0.76 | 0.11748 |
| RS19 | 5 | 1.09 | 0.76 | 0.11748 |
| RL6 | 3 | 0.90 | 0.76 | 0.11691 |
| CAN2 | 2 | 1.09 | 0.77 | 0.11634 |
| EVL | 3 | 1.10 | 0.77 | 0.11634 |
| TYPH | 11 | 0.93 | 0.77 | 0.11238 |
| CLIC4 | 1 | 0.85 | 0.78 | 0.10958 |
| TRI25 | 2 | 0.91 | 0.78 | 0.10846 |
| ERAP1 | 6 | 1.17 | 0.78 | 0.10846 |
| PUR9 | 4 | 0.94 | 0.79 | 0.10237 |
| HV349 | 1 | 0.94 | 0.79 | 0.10182 |
| RL10 | 2 | 0.92 | 0.79 | 0.10182 |
| VNN1 | 1 | 0.93 | 0.80 | 0.09854 |
| HV551 | 1 | 0.87 | 0.80 | 0.098 |
| IPYR | 3 | 1.06 | 0.80 | 0.09691 |
| NCKPL | 2 | 0.91 | 0.80 | 0.09528 |
| KV105 | 1 | 1.16 | 0.80 | 0.09474 |
| BIP | 13 | 0.94 | 0.81 | 0.0942 |
| NAMPT | 7 | 1.04 | 0.81 | 0.08938 |
| LAMA1 | 1 | 1.13 | 0.82 | 0.0846 |
| VPS4B | 2 | 1.09 | 0.82 | 0.08407 |
| MDHM | 2 | 0.97 | 0.82 | 0.08407 |
| IGLC2 | 3 | 1.15 | 0.83 | 0.08302 |
| TXND5 | 11 | 0.94 | 0.83 | 0.08249 |
| KVD29 | 2 | 1.05 | 0.83 | 0.0804 |
| RS24 | 1 | 0.92 | 0.83 | 0.07988 |
| CO3 | 7 | 0.98 | 0.83 | 0.07988 |
| TIF1B | 1 | 0.88 | 0.84 | 0.07831 |
| OAS1 | 1 | 0.87 | 0.84 | 0.07624 |
| LSM6 | 1 | 0.88 | 0.84 | 0.07572 |
| EFHD2 | 4 | 1.04 | 0.84 | 0.07572 |
| ERF3B | 1 | 1.06 | 0.84 | 0.0752 |
| STMN1 | 2 | 1.06 | 0.84 | 0.07366 |
| IGHA2 | 4 | 1.10 | 0.84 | 0.07366 |
| SNX6 | 1 | 0.95 | 0.85 | 0.07212 |
| PPID | 1 | 0.92 | 0.85 | 0.07212 |

| | | | | |
|---|---|---|---|---|
| GANAB | 4 | 0.95 | 0.85 | 0.07109 |
| ALBU | 29 | 0.95 | 0.85 | 0.07007 |
| COTL1 | 10 | 1.04 | 0.86 | 0.06803 |
| ARHG6 | 1 | 0.93 | 0.86 | 0.06753 |
| SPB10 | 1 | 0.89 | 0.86 | 0.06753 |
| RS14 | 3 | 1.05 | 0.86 | 0.06651 |
| RS13 | 2 | 1.06 | 0.87 | 0.05948 |
| CH10 | 1 | 1.04 | 0.87 | 0.05899 |
| AHSA1 | 1 | 0.96 | 0.87 | 0.05849 |
| RL27A | 2 | 0.93 | 0.88 | 0.0575 |
| ARHG1 | 1 | 0.93 | 0.89 | 0.05306 |
| HPT | 4 | 1.03 | 0.89 | 0.05306 |
| ROA0 | 2 | 1.05 | 0.89 | 0.05257 |
| HXK3 | 3 | 0.96 | 0.89 | 0.05257 |
| VASP | 10 | 0.97 | 0.89 | 0.05257 |
| S10A8 | 2 | 1.05 | 0.89 | 0.05061 |
| HEMO | 20 | 0.99 | 0.89 | 0.04964 |
| IGHD | 3 | 1.12 | 0.89 | 0.04915 |
| PLD4 | 4 | 1.04 | 0.90 | 0.04769 |
| TFG | 1 | 0.95 | 0.90 | 0.04479 |
| BAF | 1 | 0.96 | 0.91 | 0.04335 |
| PPP5 | 1 | 1.07 | 0.91 | 0.04287 |
| IGHA1 | 17 | 1.03 | 0.91 | 0.04239 |
| DEF3 | 1 | 0.96 | 0.92 | 0.0381 |
| AL1B1 | 1 | 1.07 | 0.92 | 0.03716 |
| KAP0 | 2 | 1.03 | 0.92 | 0.03574 |
| C1TC | 2 | 1.03 | 0.92 | 0.03527 |
| CD97 | 1 | 1.09 | 0.92 | 0.0348 |
| RL13 | 5 | 0.97 | 0.92 | 0.03433 |
| FIBG | 1 | 1.02 | 0.93 | 0.03245 |
| ANXA1 | 14 | 0.97 | 0.93 | 0.03198 |
| RS15A | 2 | 0.98 | 0.93 | 0.02965 |
| IGHG4 | 2 | 0.94 | 0.93 | 0.02965 |
| GNAI2 | 3 | 1.03 | 0.94 | 0.02641 |
| SNX1 | 1 | 0.98 | 0.95 | 0.02411 |
| SHBG | 1 | 1.02 | 0.95 | 0.02365 |
| PSD12 | 1 | 0.98 | 0.95 | 0.02182 |
| ELNE | 4 | 0.97 | 0.95 | 0.02182 |
| DBNL | 1 | 1.02 | 0.96 | 0.01728 |
| RS9 | 1 | 0.97 | 0.96 | 0.01592 |
| MNDA | 7 | 1.02 | 0.96 | 0.01592 |
| PRS6A | 1 | 1.02 | 0.97 | 0.01457 |
| APOC3 | 1 | 1.00 | 0.97 | 0.01457 |
| IGHG1 | 10 | 0.98 | 0.97 | 0.01412 |

| | | | | |
|---|---|---|---|---|
| **HV43D** | **1** | **0.99** | **0.97** | **0.01323** |
| **AN32E** | **2** | **0.99** | **0.97** | **0.01323** |
| **PDCD5** | **1** | **1.01** | **0.98** | **0.01055** |
| **AMPL** | **14** | **0.99** | **0.98** | **0.01055** |
| **S10A6** | **3** | **1.00** | **0.98** | **0.00745** |
| **CALR** | **6** | **0.99** | **0.98** | **0.00745** |
| **HV439** | **3** | **0.99** | **0.98** | **0.007** |
| **ANXA7** | **3** | **0.99** | **0.99** | **0.00656** |
| **RAC2** | **3** | **1.00** | **0.99** | **0.00612** |
| **PRTN3** | **3** | **0.99** | **0.99** | **0.00612** |
| **HV601** | **1** | **1.01** | **0.99** | **0.00568** |
| **RL26** | **2** | **1.01** | **0.99** | **0.00524** |
| **RS27L** | **2** | **1.01** | **0.99** | **0.00524** |
| **ROA3** | **5** | **1.00** | **0.99** | **0.00524** |
| **TBA1B** | **3** | **1.00** | **0.99** | **0.0048** |
| **FRIH** | **4** | **1.00** | **0.99** | **0.00349** |
| **PERM** | **18** | **1.00** | **1.00** | **0.00218** |

Table S2: Cell input and sequencing output of the scRNA-seq experiment.

| Subject | *Ex vivo* | Flu 6hrs | Flu 24hrs |
|---|---|---|---|
| A | 2759 | 2102 | 1871 |
| B | 2794 | 2758 | 2796 |
| C | 2535 | 2450 | 2206 |
| Total Cells per Condition | 8088 | 7310 | 6873 |
| Mean reads per cell | 55,375 | 58,215 | 62,944 |
| Median genes per cell | 2,670 | 2,786 | 3,758 |
| Doublets | 2152 | (excluded from analysis) | |
| Ambiguous | 151 | (excluded from analysis) | |

Table S3: Top 25 enriched genes in the pDC scRNA-seq clusters, by cluster

| Gene | avg_logFC | p_val_adj |
|---|---|---|
| CLUSTER 0 | | |
| FCER1G | 2.1024666 | 0 |
| C12orf75 | 2.0461347 | 0 |
| MS4A6A | 1.7763833 | 0 |
| PTPRE | 1.6217105 | 0 |
| TAGLN2 | 1.5492633 | 0 |
| ALOX5AP | 1.5456969 | 0 |
| UCP2 | 1.5376786 | 0 |
| RNASE6 | 1.5040914 | 0 |
| ZFP36L2 | 1.4492112 | 0 |
| CST3 | 1.4329711 | 0 |
| DERL3 | 1.430853 | 0 |
| PLD4 | 1.3906609 | 0 |
| PPP1R14B | 1.3824086 | 0 |
| GZMB | 1.3781429 | 0 |
| UGCG | 1.3781111 | 0 |
| TPM2 | 1.3715611 | 0 |
| CORO1A | 1.3562062 | 0 |
| CCDC50 | 1.3482743 | 0 |
| APP | 1.3357869 | 0 |
| SERPINF1 | 1.3109063 | 0 |
| SAMHD1 | 1.2736115 | 0 |
| IRF2BP2 | 1.2652994 | 0 |
| BCL11A | 1.2436351 | 0 |
| GAPT | 1.2419787 | 0 |

| Gene | avg_logFC | p_val_adj |
|---|---|---|
| CLUSTER 1 | | |
| IFNA2 | 3.549002 | 0 |
| IFNA14 | 3.36561 | 0 |
| IFNA6 | 3.331724 | 0 |
| IFNB1 | 3.189481 | 0 |
| IFNW1 | 3.158891 | 0 |
| IFNA17 | 3.157789 | 0 |
| IFNA10 | 3.105879 | 0 |
| IFNA8 | 3.069484 | 0 |
| IFNA5 | 3.052137 | 0 |
| IFNA4 | 2.975805 | 0 |
| IFNA21 | 2.94575 | 0 |
| IFNA1 | 2.645179 | 0 |
| IFNA16 | 2.627565 | 0 |
| IFNA7 | 2.594752 | 0 |
| IFNL1 | 2.532747 | 0 |
| PPP1R15A | 2.506891 | 0 |
| PUS10 | 2.17797 | 0 |
| IL12A | 2.163395 | 0 |
| CCL3 | 2.132322 | 0 |
| CCL4L2 | 2.089998 | 0 |
| BCL2A1 | 2.05844 | 0 |
| CCL4 | 2.050363 | 0 |
| TNF | 1.927204 | 0 |
| NR4A3 | 1.847532 | 0 |

| Gene | avg_logFC | p_val_adj |
|---|---|---|
| CLUSTER 2 | | |
| CCL4 | 1.702038 | 0 |
| NCF1 | 1.598255 | 0 |
| CCL4L2 | 1.555877 | 0 |
| TNF | 1.550849 | 0 |
| NABP1 | 1.539063 | 0 |
| HSPA1A | 1.527175 | 0 |
| CD83 | 1.503993 | 0 |
| LTB | 1.499888 | 0 |
| HSPA1B | 1.280324 | 0 |
| IFIT2 | 1.265776 | 0 |
| CCL3 | 1.229196 | 0 |
| CDKN1A | 1.226971 | 0 |
| LTA | 1.207946 | 0 |
| CD40 | 1.157032 | 0 |
| LDLRAD4 | 1.143193 | 0 |
| HSP90AA1 | 1.099246 | 0 |
| CCND2 | 1.086032 | 0 |
| TYW3 | 1.013196 | 0 |
| RGS1 | 1.003052 | 5.5E-284 |
| PPP1R15A | 0.996728 | 0 |
| CALCRL | 0.986798 | 5.6E-192 |
| ATF3 | 0.967541 | 1.5E-222 |
| HSPA8 | 0.92811 | 0 |
| CCL3L1 | 0.924779 | 1.5E-108 |

| Gene | avg_logFC | p_val_adj |
|---|---|---|
| CLUSTER 3 | | |
| IFI27 | 1.774985 | 0 |
| PNOC | 1.45832 | 5.1E-236 |
| TNFSF10 | 1.341827 | 0 |
| NEAT1 | 1.303959 | 0 |
| IFI6 | 1.268901 | 0 |
| CXCL13 | 1.261145 | 3.4E-165 |
| FTL | 1.205017 | 0 |
| DUSP5 | 1.184991 | 0 |
| CLEC2D | 1.167247 | 0 |
| HIST1H1C | 1.135166 | 0 |
| GAPDH | 1.131192 | 0 |
| ATOX1 | 1.07544 | 0 |
| LY6E | 1.022173 | 0 |
| ISG15 | 1.007866 | 0 |
| CD38 | 0.953347 | 0 |
| SCT | 0.948665 | 1.4E-102 |
| STAT1 | 0.940203 | 0 |
| COX5A | 0.911369 | 0 |
| PRDM1 | 0.8714 | 0 |
| CASP3 | 0.869088 | 0 |
| RSAD2 | 0.846212 | 0 |
| SAT1 | 0.824952 | 1.9E-293 |
| P2RY10 | 0.80955 | 0 |
| TRIB1 | 0.804212 | 0 |

| Gene | avg_logFC | p_val_adj |
|---|---|---|
| **CLUSTER 4** | | |
| CD44 | 1.59138 | 0 |
| CXCL10 | 1.428058 | 0 |
| NMB | 1.424394 | 1.79E-88 |
| LTA | 1.221621 | 0 |
| CXCL11 | 1.20654 | 5.9E-127 |
| CD40 | 1.09721 | 0 |
| CCL4L2 | 1.073064 | 0 |
| SLAMF7 | 1.063316 | 0 |
| ID2 | 1.058091 | 0 |
| CD83 | 1.049015 | 0 |
| CALCRL | 1.02762 | 0 |
| FAM129A | 1.025988 | 0 |
| HSPA1A | 1.025384 | 0 |
| ALCAM | 1.018413 | 0 |
| REL | 1.003306 | 2.5E-303 |
| BCL2A1 | 0.999812 | 0 |
| MIR155HG | 0.986837 | 0 |
| TFEC | 0.926733 | 0 |
| CFLAR | 0.89305 | 0 |
| BCL2L1 | 0.891324 | 0 |
| CCL4 | 0.881024 | 0 |
| NFKB1 | 0.859435 | 0 |
| HERC5 | 0.85307 | 0 |
| C15orf48 | 0.843186 | 1.7E-233 |

| Gene | avg_logFC | p_val_adj |
|---|---|---|
| **CLUSTER 5** | | |
| CXCL13 | 1.800695 | 0 |
| CCR7 | 1.710081 | 0 |
| TMSB4X | 1.693879 | 0 |
| TUBA1A | 1.626864 | 0 |
| SOX4 | 1.622532 | 0 |
| BASP1 | 1.596734 | 0 |
| FSCN1 | 1.446334 | 0 |
| ZFP36L1 | 1.398223 | 0 |
| CD70 | 1.386201 | 0 |
| IFI27 | 1.374846 | 0 |
| LMNB1 | 1.326046 | 0 |
| MARCKS | 1.299956 | 0 |
| IFITM3 | 1.277995 | 0 |
| CKB | 1.273303 | 0 |
| LY6E | 1.206548 | 0 |
| GAPDH | 1.191684 | 0 |
| TUBB | 1.163445 | 0 |
| SERPINB1 | 1.158247 | 0 |
| RAB9A | 1.135876 | 0 |
| S100A11 | 1.119249 | 0 |
| PKM | 1.11292 | 0 |
| CLEC2D | 1.112227 | 0 |
| BID | 1.1038 | 0 |
| TNFSF4 | 1.0764 | 0 |

| Gene | avg_logFC | p_val_adj |
|---|---|---|
| | CLUSTER 6 | |
| MT2A | 1.32218 | 0 |
| EAF2 | 1.144386 | 0 |
| GAPDH | 1.141306 | 0 |
| TPI1 | 1.079615 | 0 |
| CCR7 | 1.077027 | 0 |
| NEAT1 | 1.016605 | 0 |
| LDHA | 0.976995 | 0 |
| CLEC2D | 0.951816 | 0 |
| HIST1H1C | 0.939304 | 0 |
| CLEC2B | 0.921002 | 0 |
| MPC2 | 0.910105 | 0 |
| TALDO1 | 0.907717 | 0 |
| GBP1 | 0.902111 | 0 |
| UCHL1 | 0.899496 | 0 |
| SPATS2L | 0.898494 | 0 |
| CSRP2 | 0.898457 | 0 |
| CMTM7 | 0.880067 | 0 |
| PKM | 0.879726 | 0 |
| IFIT1 | 0.865848 | 0 |
| IFI27 | 0.845355 | 0 |
| CTSC | 0.844856 | 4.2E-275 |
| PRDX1 | 0.841203 | 0 |
| FTL | 0.839755 | 0 |
| TYMP | 0.829577 | 0 |

| Gene | avg_logFC | p_val_adj |
|---|---|---|
| | CLUSTER 7 | |
| LTB | 1.369874 | 0 |
| TYW3 | 1.01182 | 0 |
| HSPA1A | 0.995985 | 0 |
| BTG1 | 0.91905 | 0 |
| HSPA8 | 0.870512 | 0 |
| BBX | 0.860342 | 0 |
| LAP3 | 0.833926 | 0 |
| DCPS | 0.802439 | 2.1E-250 |
| CSF2RB | 0.795061 | 3.1E-300 |
| PAG1 | 0.779625 | 0 |
| HSP90AA1 | 0.77947 | 0 |
| SMC6 | 0.772563 | 0 |
| FYB1 | 0.769351 | 1.7E-303 |
| SLC12A2 | 0.768546 | 5.4E-266 |
| EIF2AK2 | 0.762227 | 0 |
| SERPING1 | 0.759044 | 0 |
| CRYZ | 0.751272 | 0 |
| TSPAN13 | 0.745545 | 0 |
| TCL1A | 0.738809 | 4.82E-61 |
| UBE2L6 | 0.738395 | 0 |
| PIM3 | 0.734997 | 0 |
| SAMD9 | 0.730834 | 0 |
| DDX60 | 0.725135 | 0 |
| DNASE1L3 | 0.721615 | 0 |

| Gene | avg_logFC | p_val_adj |
|---|---|---|
| CLUSTER 8 | | |
| CCND2 | 1.643757 | 0 |
| KHK | 1.294032 | 7.8E-208 |
| CCL19 | 1.163064 | 2.12E-94 |
| MYBL2 | 1.13259 | 0 |
| BTG1 | 1.108356 | 0 |
| ERICH3 | 0.859103 | 5.5E-185 |
| TCL1A | 0.827502 | 3.87E-95 |
| CALR | 0.774469 | 0 |
| NLRP7 | 0.764753 | 0 |
| NKG7 | 0.731817 | 5.4E-129 |
| SYNGR2 | 0.716912 | 1.2E-235 |
| IFI27 | 0.712686 | 2.3E-159 |
| CTSZ | 0.699129 | 7.1E-223 |
| GPX4 | 0.699015 | 2.3E-254 |
| HSP90B1 | 0.692583 | 1.1E-255 |
| WNT10A | 0.688987 | 5.9E-284 |
| TXN | 0.686913 | 2.3E-146 |
| PNOC | 0.64691 | 9.77E-46 |
| COX5A | 0.646351 | 3.3E-268 |
| SCT | 0.643599 | 8.3E-114 |
| VOPP1 | 0.626633 | 1.4E-234 |
| BTG2 | 0.621961 | 3.7E-132 |
| P2RY6 | 0.616117 | 6.4E-221 |
| IGHM | 0.61362 | 2.1E-278 |

| Gene | avg_logFC | p_val_adj |
|---|---|---|
| CLUSTER 9 | | |
| CXCL11 | 1.83922 | 2.1E-253 |
| RSAD2 | 1.697238 | 3.1E-295 |
| CXCL10 | 1.634519 | 1.1E-204 |
| HSPA1A | 1.127474 | 1.9E-192 |
| BTG1 | 1.092461 | 1.5E-159 |
| HMOX1 | 1.081575 | 1.58E-61 |
| PARP14 | 1.023822 | 2.5E-245 |
| TFEC | 1.014234 | 1.3E-215 |
| AKR1C3 | 0.996355 | 2.2E-153 |
| EMP3 | 0.990186 | 1.9E-197 |
| ENDOG | 0.962219 | 3.93E-94 |
| CSF2RB | 0.949516 | 2.4E-141 |
| RNF213 | 0.916677 | 1.2E-224 |
| HSPA1B | 0.894641 | 1E-132 |
| OAS3 | 0.883837 | 2.9E-206 |
| HMGCS1 | 0.883055 | 2E-128 |
| CXorf21 | 0.87998 | 2.25E-81 |
| SQSTM1 | 0.830337 | 1.4E-103 |
| MX1 | 0.810703 | 9.9E-192 |
| PIM3 | 0.808803 | 3.3E-166 |
| CMPK2 | 0.801006 | 3.9E-189 |
| LAP3 | 0.798204 | 2.1E-202 |
| DENND1B | 0.794405 | 6.4E-187 |
| CXCL9 | 0.794203 | 1.8E-75 |

| Gene | avg_logFC | p_val_adj |
|---|---|---|
| CLUSTER 10 | | |
| MALAT1 | 1.213465 | 2.18E-65 |
| XIST | 0.930008 | 3.13E-83 |
| NEAT1 | 0.816518 | 4.92E-39 |
| N4BP2L2 | 0.689664 | 3.67E-62 |
| MT-ND5 | 0.657111 | 8.48E-56 |
| POLR2J3.1 | 0.648099 | 6.96E-39 |
| CHD9 | 0.622404 | 3.93E-35 |
| DDX17 | 0.613559 | 4.82E-50 |
| MT-CO1 | 0.580231 | 7.42E-23 |
| PNISR | 0.579236 | 5.13E-43 |
| HNRNPH1 | 0.566843 | 8.18E-44 |
| MT-ND2 | 0.55709 | 1.23E-51 |
| ARID1B | 0.551148 | 1.61E-37 |
| ANKRD12 | 0.545767 | 1.94E-39 |
| RSRP1 | 0.543745 | 1.48E-35 |
| FTX | 0.541207 | 2.98E-36 |
| RBM39 | 0.532532 | 1.34E-39 |
| PNN | 0.525924 | 5.7E-32 |
| RNF213 | 0.522187 | 8.38E-23 |
| ZNF207 | 0.521197 | 1.74E-37 |
| CCNL1 | 0.519185 | 7.36E-32 |
| INTS6 | 0.518957 | 4.69E-43 |
| ZEB2 | 0.517619 | 7.99E-31 |
| CCDC88A | 0.510893 | 1.71E-35 |

| Gene | avg_logFC | p_val_adj |
|---|---|---|
| CLUSTER 11 | | |
| IFNA1 | 3.23627 | 4.3E-270 |
| IFNA16 | 3.209599 | 3.2E-243 |
| CYTOR | 3.129515 | 4.2E-286 |
| IFNA4 | 2.93449 | 1.8E-241 |
| IFNA21 | 2.841742 | 1.1E-259 |
| IFNL1 | 2.831011 | 7E-175 |
| IFNA8 | 2.686882 | 1.4E-227 |
| IFNA17 | 2.634333 | 9.1E-205 |
| IFNA5 | 2.622508 | 6.6E-286 |
| MIR4435-2HG | 2.60344 | 0 |
| CCL5 | 2.537599 | 1.1E-156 |
| IFNW1 | 2.459461 | 3E-239 |
| IFNB1 | 2.435702 | 2.8E-228 |
| IFNA10 | 2.431466 | 2.5E-222 |
| GPR34 | 2.346484 | 0 |
| IFNA6 | 2.338089 | 4.1E-118 |
| DKK1 | 2.331726 | 0 |
| CXCL8 | 2.218949 | 1.8E-295 |
| IFNA14 | 2.205407 | 2.6E-209 |
| IFNA7 | 2.200717 | 0 |
| IFNE | 2.190019 | 0 |
| CCL3L1 | 2.161842 | 2.3E-186 |
| BET1 | 2.15487 | 5.1E-254 |
| CCL3 | 2.075043 | 4.2E-173 |

| Gene | avg_logFC | p_val_adj |
|---|---|---|
| CLUSTER 12 | | |
| TMEM267 | 1.622947 | 7.6E-122 |
| CTSC | 1.412497 | 7.8E-144 |
| DUSP11 | 1.365048 | 9.2E-156 |
| IL12A | 1.318169 | 1.9E-135 |
| GRSF1 | 1.187564 | 1.2E-157 |
| PNOC | 1.167879 | 1.3E-42 |
| CCR7 | 1.152894 | 2.5E-126 |
| RGS1 | 1.14512 | 7.33E-48 |
| IFI27 | 1.137594 | 1.1E-107 |
| IGFBP4 | 1.092764 | 1.2E-146 |
| HARS | 1.083338 | 3.1E-105 |
| MPC2 | 1.05995 | 6.7E-112 |
| CMTM7 | 1.050202 | 1.4E-139 |
| CLECL1 | 1.034513 | 1.6E-121 |
| CAB39L | 1.028687 | 1.8E-197 |
| MARCKSL1 | 1.024607 | 1.8E-80 |
| RNF115 | 1.019902 | 7.58E-82 |
| GPR34 | 0.960441 | 9.67E-57 |
| CD48 | 0.943073 | 1.59E-40 |
| GPR160 | 0.934233 | 2.3E-174 |
| CLEC2D | 0.919263 | 7.5E-106 |
| REL | 0.917475 | 2E-110 |
| CRYBG1 | 0.900703 | 2.1E-206 |
| ENPP2 | 0.898065 | 1.41E-87 |

| Gene | avg_logFC | p_val_adj |
|---|---|---|
| CLUSTER 13 | | |
| IGHA1 | 6.122379 | 2.6E-143 |
| IGLC3 | 4.976468 | 1 |
| IGKC | 4.781452 | 1.06E-07 |
| IGHG2 | 4.599817 | 7.8E-281 |
| IGLC2 | 4.517642 | 1 |
| IGHG4 | 4.310132 | 6.6E-148 |
| IGHG1 | 4.141818 | 2.1E-167 |
| IGLC7 | 4.047892 | 4.12E-33 |
| IGHA2 | 3.653254 | 0 |
| IGHM | 3.340308 | 0.000234 |
| IGHG3 | 3.121733 | 0 |
| JCHAIN | 2.643538 | 6.5E-102 |
| CD79A | 2.092111 | 0 |
| IGLL5 | 1.981569 | 1.8E-283 |
| CD27 | 1.936166 | 0 |
| TNFRSF17 | 1.820993 | 3.1E-174 |
| POU2AF1 | 1.777515 | 0 |
| MZB1 | 1.743814 | 2.23E-98 |
| FKBP11 | 1.612301 | 6.2E-152 |
| KLF2 | 1.533383 | 0 |
| TENT5C | 1.450866 | 0 |
| IGKV4-1 | 1.360235 | 0 |
| PIM2 | 1.184434 | 1E-108 |
| CD52 | 1.177831 | 3.4E-82 |

| Gene | avg_logFC | p_val_adj |
|---|---|---|
| CLUSTER 14 | | |
| CCND2 | 1.673926 | 7.29E-63 |
| PNOC | 1.511161 | 2.7E-20 |
| PCLAF | 1.30334 | 0 |
| TYMS | 1.298088 | 0 |
| PCNA | 1.036432 | 5.1E-85 |
| IFI27 | 0.984043 | 8.59E-26 |
| CLSPN | 0.973058 | 0 |
| MCM4 | 0.956267 | 0 |
| CEP55 | 0.922884 | 2.16E-97 |
| NASP | 0.870973 | 7.03E-52 |
| GAPDH | 0.867348 | 6.01E-34 |
| GINS2 | 0.862674 | 0 |
| CKS1B | 0.852369 | 1.79E-49 |
| EDNRB | 0.845541 | 4.69E-40 |
| SCT | 0.840879 | 2.31E-13 |
| CTSC | 0.829415 | 8.46E-18 |
| MCM7 | 0.811443 | 3E-107 |
| CHEK1 | 0.807757 | 0 |
| CLECL1 | 0.802669 | 4.91E-28 |
| H2AFZ | 0.802205 | 5.25E-33 |
| IGHM | 0.793891 | 2.11E-34 |
| MCM3 | 0.789192 | 6.32E-61 |
| COX5A | 0.781364 | 8.36E-40 |
| TUBA1B | 0.781008 | 8.57E-40 |

| Gene | avg_logFC | p_val_adj |
|---|---|---|
| CLUSTER 15 | | |
| PLCG2 | 4.288975 | 8.08E-58 |
| HEXIM1 | 2.578878 | 1.11E-60 |
| JUN | 2.402232 | 4.64E-42 |
| GADD45B | 2.249662 | 1.04E-45 |
| IER3 | 2.245483 | 1.09E-27 |
| NFKBIA | 2.214461 | 7.69E-24 |
| SNHG12 | 2.176202 | 2.27E-83 |
| INTS6 | 2.146815 | 1.84E-36 |
| IER5 | 2.109135 | 8.03E-45 |
| RASD1 | 2.051184 | 4.96E-67 |
| IER2 | 1.985599 | 1.69E-29 |
| MAFB | 1.953123 | 0 |
| FOS | 1.934448 | 3.7E-144 |
| SRSF7 | 1.919737 | 4.55E-39 |
| PMAIP1 | 1.84026 | 6.3E-29 |
| SOCS1 | 1.80901 | 2.87E-43 |
| EGR1 | 1.769766 | 4.5E-264 |
| ID2 | 1.763281 | 3.44E-36 |
| PLK2 | 1.762039 | 2.1E-301 |
| DNAJB1 | 1.74626 | 1.44E-05 |
| TUBB4B | 1.724608 | 1.26E-27 |
| JUNB | 1.68239 | 5.9E-39 |
| CXCL8 | 1.644674 | 0.524263 |
| HSPA1B | 1.624778 | 3.9E-31 |

| Gene | avg_logFC | p_val_adj |
|---|---|---|
| CLUSTER 16 | | |
| GIMAP7 | 1.851219 | 0 |
| IL7R | 1.678736 | 1.7E-105 |
| TRBC2 | 1.656116 | 0 |
| GZMK | 1.640609 | 0 |
| CD79A | 1.640104 | 5.53E-14 |
| IGLC3 | 1.604881 | 1 |
| GIMAP4 | 1.549939 | 0 |
| MT2A | 1.48322 | 7.18E-14 |
| EVL | 1.457164 | 5.3E-103 |
| LIMD2 | 1.346851 | 7.97E-26 |
| KLF2 | 1.333666 | 2.7E-184 |
| IL32 | 1.263929 | 0 |
| TRAC | 1.210002 | 0 |
| TRBC1 | 1.201439 | 0 |
| CD52 | 1.186019 | 5.96E-22 |
| CXCR4 | 1.185677 | 2.12E-08 |
| AC245297.3 | 1.154835 | 1.72E-48 |
| ITM2B | 1.146894 | 4.57E-30 |
| CD3D | 1.11465 | 0 |
| TRDC | 1.091593 | 1.2E-119 |
| GBP1 | 1.084558 | 1.07E-25 |
| CD48 | 1.049761 | 1.29E-24 |
| GBP4 | 1.045723 | 3.16E-19 |
| SMCHD1 | 1.031087 | 4.5E-18 |

| Gene | avg_logFC | p_val_adj |
|---|---|---|
| CLUSTER 17 | | |
| LYZ | 3.203738 | 0 |
| S100A10 | 2.288291 | 5.96E-20 |
| S100A4 | 2.078156 | 9.89E-17 |
| COTL1 | 1.854883 | 1.88E-53 |
| CST3 | 1.586403 | 2.32E-15 |
| PPP1R14A | 1.262968 | 9.48E-31 |
| FOS | 1.190144 | 7.8E-125 |
| FGL2 | 1.179553 | 1.1E-104 |
| AXL | 1.130974 | 3.69E-93 |
| AIF1 | 1.109946 | 1.34E-13 |
| ITGB2 | 1.085394 | 3.83E-20 |
| FCGRT | 1.077133 | 3.17E-14 |
| CORO1A | 1.076163 | 6.27E-12 |
| ANXA1 | 1.025688 | 3.08E-42 |
| TXNIP | 1.021241 | 4.54E-10 |
| VIM | 1.016814 | 1.78E-08 |
| OTULINL | 0.992976 | 6.86E-19 |
| SAMHD1 | 0.974915 | 1.7E-10 |
| ACTG1 | 0.964491 | 8.87E-09 |
| S100A6 | 0.951039 | 5.65E-10 |
| DAB2 | 0.909922 | 8.88E-13 |
| CCND3 | 0.885548 | 4.27E-07 |
| LIMD2 | 0.877811 | 6.07E-11 |
| CTSH | 0.87696 | 6.38E-10 |
